# Multi-scale haplotype signatures distinguish among evolutionary processes

**DOI:** 10.64898/2026.10.30.763096

**Authors:** Mohammad Khan, Sandipan Paul Arnab, Md Ruhul Amin, Cindy G. Santander, Michael DeGiorgio

**Affiliations:** Department of Electrical Engineering and Computer Science, Florida Atlantic University, Boca Raton, FL, USA; Center for Omics technologies and Data Engineering, Florida Atlantic University, Boca Raton, FL, USA; Department of Biomedical Engineering, Florida Atlantic University, Boca Raton, FL, USA

**Author notes:** Department of Molecular and Cell Biology, University of California, Berkeley, CA, USA.

## Abstract

Natural selection embeds distinct genomic signatures as populations adapt to environmental pressures. Machine learning approaches, particularly deep neural networks, have emerged as powerful tools to detect adaptive signatures from genomic data. While machine learning has dramatically improved our ability to extract these signals from sequence variation, distinguishing among multiple evolutionary processes remains challenging. We introduce *SHADES*, a deep learning framework that employs information about sequence differences and their frequencies at various spatial resolutions to capture evolutionary signatures at different genomic scales. We found that *SHADES* has the capacity to distinguish among neutrality, selective sweeps, balancing selection, and adaptive introgression and outperformed several contemporary methods evaluated for both classification accuracy and power. In addition, *SHADES* remained robust to a number of confounding factors, with only modest changes in classification performance. Furthermore, application of *SHADES* to empirical haplotypic variation from European humans recovered previously reported signals for selective sweeps, balancing selection, and adaptive introgression, while also identifying novel genes broadly involved in immune defense, inflammatory regulation, and epithelial barrier biology.

## Introduction

Genetic variation arises primarily through mutation, which provides the raw material upon which evolutionary forces act [Kimura, 1985, Barton, 2010, Loewe and Hill, 2010]. Natural selection is one of the central mechanisms shaping this variation, favoring alleles that increase fitness while removing those that are deleterious [McVicker et al., 2009]. Over time, this process alters variation in populations and drives divergence between populations, and depending on the evolutionary context, it leaves behind distinct genomic footprints [Gillespie, 2004]. In addition to selection, other forces such as genetic drift, recombination, and demographic history interact to influence genomic variation, particularly in complex systems like human genomes [Ellegren and Galtier, 2016, Retel et al., 2019]. Natural selection itself can occur in different forms, such as positive and balancing selection, with each expected to leave characteristic genomic signatures, though these signatures may be modified by the effects of drift, recombination, and demographic history [Vitti et al., 2013]. Accurately identifying and distinguishing among these processes is essential for understanding the evolutionary dynamics that have influenced genome variation within and across populations.

Neutrality serves as a baseline against which selective events are measured, where genetic changes are governed primarily by nonadaptive processes such as mutation, recombination, genetic drift, and migration rather than selection [Duret, 2008]. Selective sweeps occur when a beneficial mutation rapidly increases in frequency via positive selection, reducing genetic variation at the selected locus and in surrounding regions due to hitchhiking effects [Smith and Haigh, 1974, Duret, 2008, Vitti et al., 2013]. Balancing selection, in contrast, maintains genetic diversity by favoring heterozygotes or multiple alleles over time [Charlesworth, 2006, Vitti et al., 2013]. Adaptive introgression introduces beneficial alleles from one population or species into another, followed by positive selection that retains these alleles in the recipient population [Racimo et al., 2015, Setter et al., 2020].

Detecting even a single mode of selection in genomic data is challenging due to overlapping genomic signatures that make it difficult to distinguish neutrality and selection, while discriminating among multiple selective processes is even more difficult because their genomic signatures can overlap substantially. The stochastic nature of genetic drift and recombination can obscure true signals, especially in regions with low diversity or sparse sampling [Star and Spencer, 2013]. Demographic events, such as bottlenecks or population structure, often confound selection signals, leading to false positives or negatives [Jensen et al., 2005, Zheng and Wiehe, 2019, Min et al., 2022]. Furthermore, some evolutionary phenomena operate within narrow genomic regions and can be easily masked by these confounding factors making them especially difficult to detect. Additionally, most existing methods are tailored to detect specific phenomena in isolation, limiting their ability to integrate multiple evolutionary processes. This fragmentation hinders the construction of a unified view of selection across the genome.

Neutral and different selective processes leave distinct yet subtle patterns in genomic variation, so statistical tests are widely used to distinguish among them [Hudson et al., 1987, Tajima, 1989, Fay and Wu, 2003]. In particular, selective sweeps are expected to reduce local diversity and elevate the frequency of one or a few haplotypes around the locus under selection [Przeworski, 2002, Hermisson and Pennings, 2005, Pennings and Hermisson, 2006]. In contrast, adaptive introgression can also create localized dips in diversity but yields a different haplotype structure, because the advantageous haplotype is imported from a donor population or species [Huerta-Sánchez et al., 2014, Rees et al., 2020]. Long-term balancing selection maintains alleles at intermediate frequencies and elevates local polymorphism, but the genomic footprint is typically confined by recombination to a relatively small interval around the selected site [Hudson and Kaplan, 1988, Charlesworth, 2006, Leffler et al., 2013, Siewert and Voight, 2017, Bitarello et al., 2018]. Consequently, statistics are most informative in narrow windows that capture excess diversity and the characteristic clustering of variants at similar frequencies, rather than in broad windows that dilute the signal [Siewert and Voight, 2017, Bitarello et al., 2018, Cheng and DeGiorgio, 2019, 2020]. Because balanced lineages persist for long periods, haplotypes linked to alternative alleles accumulate mutations independently, leading to deeper coalescence and greater sequence divergence among haplotypes than expected under neutrality [Charlesworth, 2006, DeGiorgio et al., 2014, Bitarello et al., 2018]. In contrast, neutral regions lack both the sustained excess of intermediate-frequency polymorphisms and the haplotype divergence that mirror loci under long-term balancing selection [Fijarczyk and Babik, 2015]. Machine learning methods extend these statistical approaches by learning complex, multidimensional relationships among different statistics and have been shown to distinguish among a number of evolutionary processes [Kern and Schrider, 2018, Sugden et al., 2018, Torada et al., 2019, Kumar et al., 2022, Lauterbur et al., 2023, Min et al., 2026].

More recently, deep learning architectures such as convolutional neural networks (CNNs) have demonstrated strong performance in evolutionary genomics by capturing spatial dependencies and subtle patterns in genomic variation [Cecil and Sugden, 2023, Korfmann et al., 2023, Huang et al., 2024, Xu et al., 2026]. CNNs excel at image-based classification by integrating local and contextual information, enabling them to detect signals that traditional methods may overlook [Goodfellow et al., 2016a]. Their flexibility also enables efficient training of large models via transfer learning [Pan and Yang, 2009, Goodfellow et al., 2016a, Arnab et al., 2025], where networks pretrained on large natural image datasets serve as feature extractors or initializations for fine-tuning on genomic data. This strategy enables scalable, efficient training while retaining the capacity to recognize complex evolutionary signatures [Sharif Razavian et al., 2014, Arnab et al., 2025]. Transfer learning can materially reduce computational and energy costs by replacing full training from scratch with comparatively brief fine-tuning runs, thereby lowering wall-clock time and associated carbon emissions [Hosseinzadeh Taher et al., 2021, Biswas et al., 2022, Bouza et al., 2023, Bolón-Canedo et al., 2024].

In this study, we present *SHADES* (Statistics of Haplotype frequencies And Differences for Evolutionary Signatures), which applies CNNs to image of three different summary statistics carefully selected to discriminate among four classes of evolutionary processes: neutrality, balancing selection, selective sweeps, and adaptive introgression. We developed two multiscale CNN architectures that integrate features from three summary statistic images through parallel branches, with one using compact, shallow networks designed for computational efficiency and the other using larger Inception-ResNet-v2 networks initialized with ImageNet pretrained weights to enable finetuning. Using extensive forward-time simulation data for training and validation, *SHADES* learns to recognize the characteristic genomic patterns produced by each process. It accurately predicts evolutionary parameters while remaining robust to biological and technical confounding factors such as missing genomic regions, mutation and recombination rate variation, and background selection. We benchmarked *SHADES* against three existing methods for detecting adaptive events. Our approach demonstrates superior accuracy across all comparison groups, proving highly effective in distinguishing between neutrality, balancing selection, selective sweeps, and adaptive introgression. To validate the model on empirical data, we applied *SHADES* to whole-genome sequences from European humans [The 1000 Genomes Project Consortium, 2015]. It successfully recapitulated established candidates for all defined classes, including *LCT* as selective sweep, the *HLA* region as balancing selection, and *BNC2* as adaptive introgression. Furthermore, the model identified several novel candidate genes subject to these selective forces. *SHADES* is available as open-source software to facilitate research and insights into the complex interplay of evolutionary forces shaping diverse species at https://github.com/md-bd/SHADES.

## Results

### Models to characterize adaptive evolution

We trained four CNN models (*M*_1_, *M*_2_, and *SHADES* [*M*] for *M* ∈ {*M*_1_*, M*_2_}) using windows of size 11, 21, and 51 SNPs to distinguish among neutrality, sweeps, adaptive introgression, and balancing selection (see *Model architecture* and *Training* subsections of the *Methods*). *M*_1_ is a relatively shallow CNN, whereas *M*_2_ uses a significantly deeper CNN architecture. *M*_1_ and *M*_2_ architectures are trained separately at each window size, while their corresponding *SHADES* [*M*_1_] and *SHADES* [*M*_2_] models integrate information across window sizes (Figure 1). The models were trained using haplotype data generated with the forward-time simulator SLiM [Haller and Messer, 2019], evolving sequences of length two megabases (Mb) under a demographic history consistent with the European CEU human population [Terhorst et al., 2017], and mutation and recombination rates consistent with empirical human estimates [Payseur and Nachman, 2000, Scally and Durbin, 2012]. The simulated replicates were each processed to generate three images summarizing haplotype variation for each replicate, which were later used for model training and evaluation (see *Input image generation* subsection of the *Methods*). These images capture of haplotype frequency distributions and differences at multiple genomic scales to tease apart subtle signals differentiating evolutionary forces. The *M*_1_ and *M*_2_ models follow a conventional CNN architecture [Krizhevsky et al., 2012, Szegedy et al., 2017], whereas the *SHADES* model incorporates a stacking approach. The *SHADES* [*M*] model, for *M* ∈ {*M*_1_*, M*_2_}, concatenates the penultimate layer activations from three *M*_1_ or *M*_2_ models pretrained on images with windows of size 11, 21, and 51 SNPs, followed by input to a two-layer fully connected model (see *Model architecture* subsection of the *Methods*).

**Figure 1:**
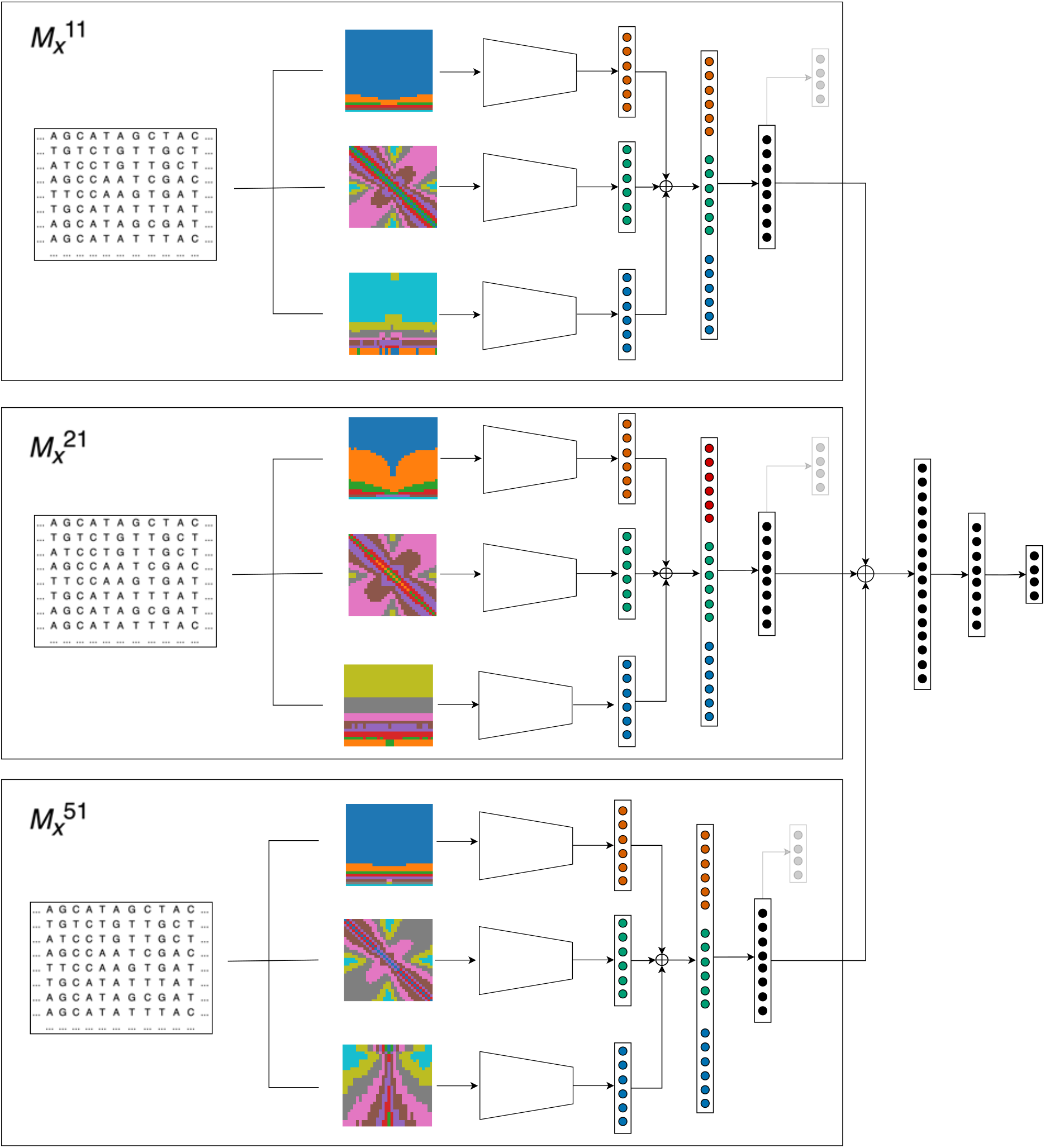
High-level schematic of the stacked *SHADES* models. Each stacked model integrates information across three genomic window sizes (11, 21, and 51 SNPs) by combining three independently trained base models of the same type (*M_x_*; architecture shown in Figure S1). For a given *x* 1, 2, inputs from each window size are processed by the corresponding *M_x_* sub model (*M*_*x*_^11^, *M*_*x*_^21^, and *M*_*x*_^51^) to produce a learned feature representation from its penultimate layer. The three penultimate layer feature vectors (one per window size) are concatenated (⊕) to form a multiscale feature vector, which is then passed through two fully connected layers with ReLU activation. A final four-dimensional output layer with softmax activation produces class probabilities for neutrality, sweep, adaptive introgression, and balancing selection. The stacked *SHADES* [*M_x_*] models differ only in the underlying base architecture *M_x_* used for each window size, while sharing the same multiscale concatenation and classifier head.

### Performance of *M*_1_, *M*_2_, and *SHADES* models

To evaluate the classification performance of our models, we generated 1,000 test replicates of neutral, sweep, adaptive introgression, and balancing selection settings with identical distributions as used for training replicates. We find that our tested models demonstrate strong ability in distinguishing among the four classes. While the *M*_1_ and *M*_2_ models performed well individually (Figures S2 and S3, respectively), stacking the best models from different window sizes further improved accuracy in both *SHADES* models (Figure 2). In particular, the *M*_1_ models attained accuracies of 74.93, 71.23, and 74.23% respectively for window sizes of 11, 21, and 51 (Figure S2A), with *SHADES* [*M*_1_] outperforming them at 80.75% (Figure 2A). The *M*_2_ models reached accuracies of 86.10, 84.08, and 80.85% for window sizes of 11, 21, and 51 SNPs, respectively (Figure S3A), and stacking these models further improved performance, with *SHADES* [*M*_2_] achieving 90.85% accuracy (Figure 2).

**Figure 2:**
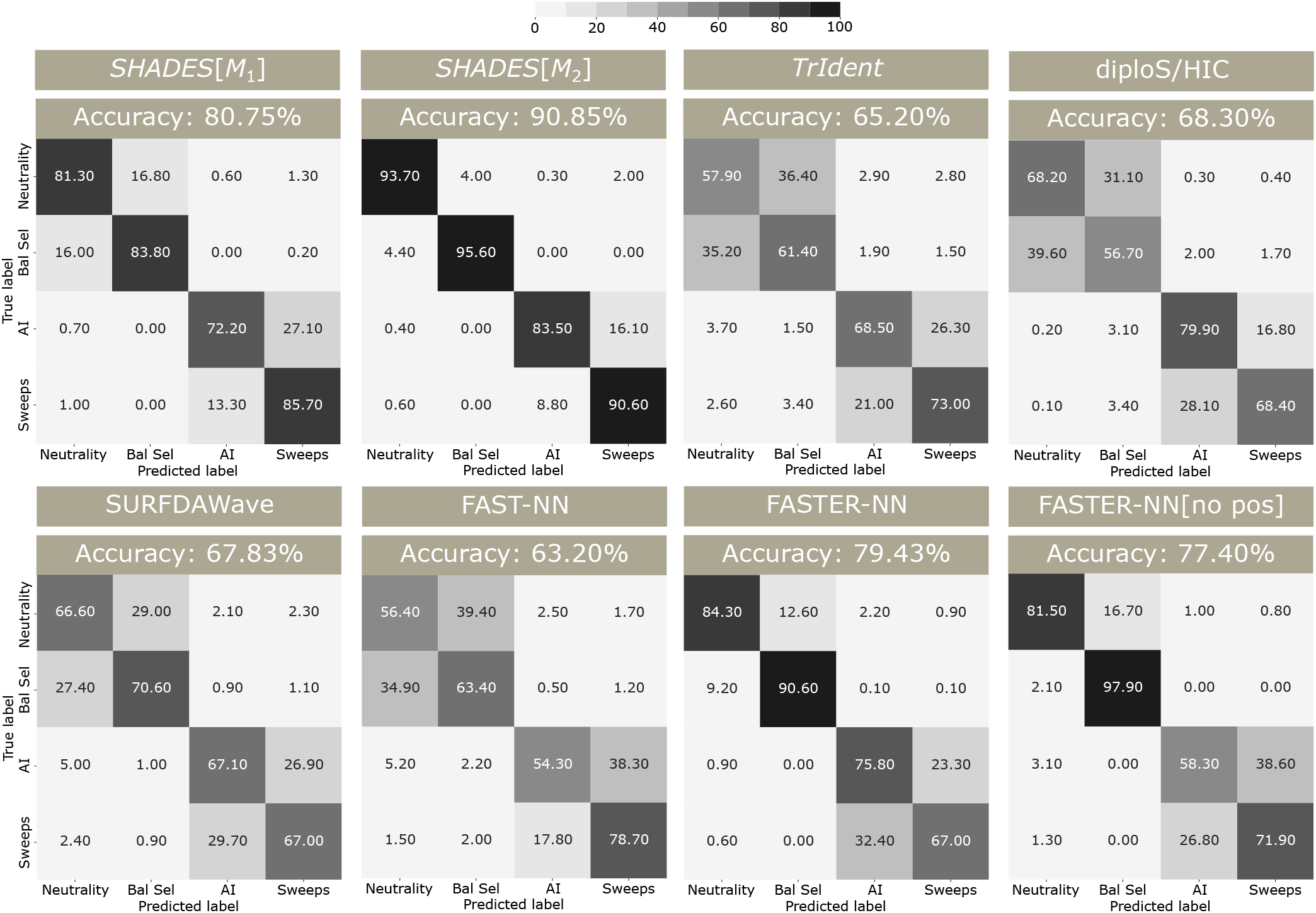
Classification rates and accuracies as depicted by confusion matrices for the *SHADES* [*M*_1_], *SHADES* [*M*_2_], diploS/HIC, *TrIdent*, *SURFDAWave*, FAST-NN, and FASTER-NN models. In the confusion matrices, AI denotes adaptive introgression and Bal Sel denotes balancing selection. The *TrIdent* model was trained and tested using the central 3,115 SNPs from each test genome replicate while retaining singletons and doubletons, as its performance improved under such settings (Figure S11). The diploS/HIC model was trained and tested with 12 summary statistics computed on 21 contiguous windows of length approximately 95 kilobases across the two Mb simulated region. The *SURFDAWave* model was trained and tested with nine summary statistics computed in 256 overlapping windows of length 10 SNPs with a stride of five SNPs. FAST-NN was trained and tested on derived allele frequencies alone, whereas FASTER-NN additionally incorporated genomic position information. FASTER-NN[no pos] was trained without the pairwise SNP distance information.

Interestingly, across all *M*_1_, *M*_2_, and *SHADES* model variants, the models frequently confused sweeps and adaptive introgression, as well as balancing selection and neutrality (Figures 2, S2A, and S3A). This misclassification pattern is reflected in the one-versus-rest receiver operating characteristic (ROC) curves, which illustrate classification power (true positive rate) as a function of false positive rate (Figures 3B, S2B, and S3B). All models demonstrated high classification power in distinguishing both sweeps and adaptive introgression from neutrality (Figures 3A, S2C, and S3C). However, the power to detect balancing selection from neutrality was significantly lower (Figures 3A, S2C, and S3C), which echos the frequent misclassifications between these two classes (Figures 2, S2A, and S3A).

**Figure 3:**
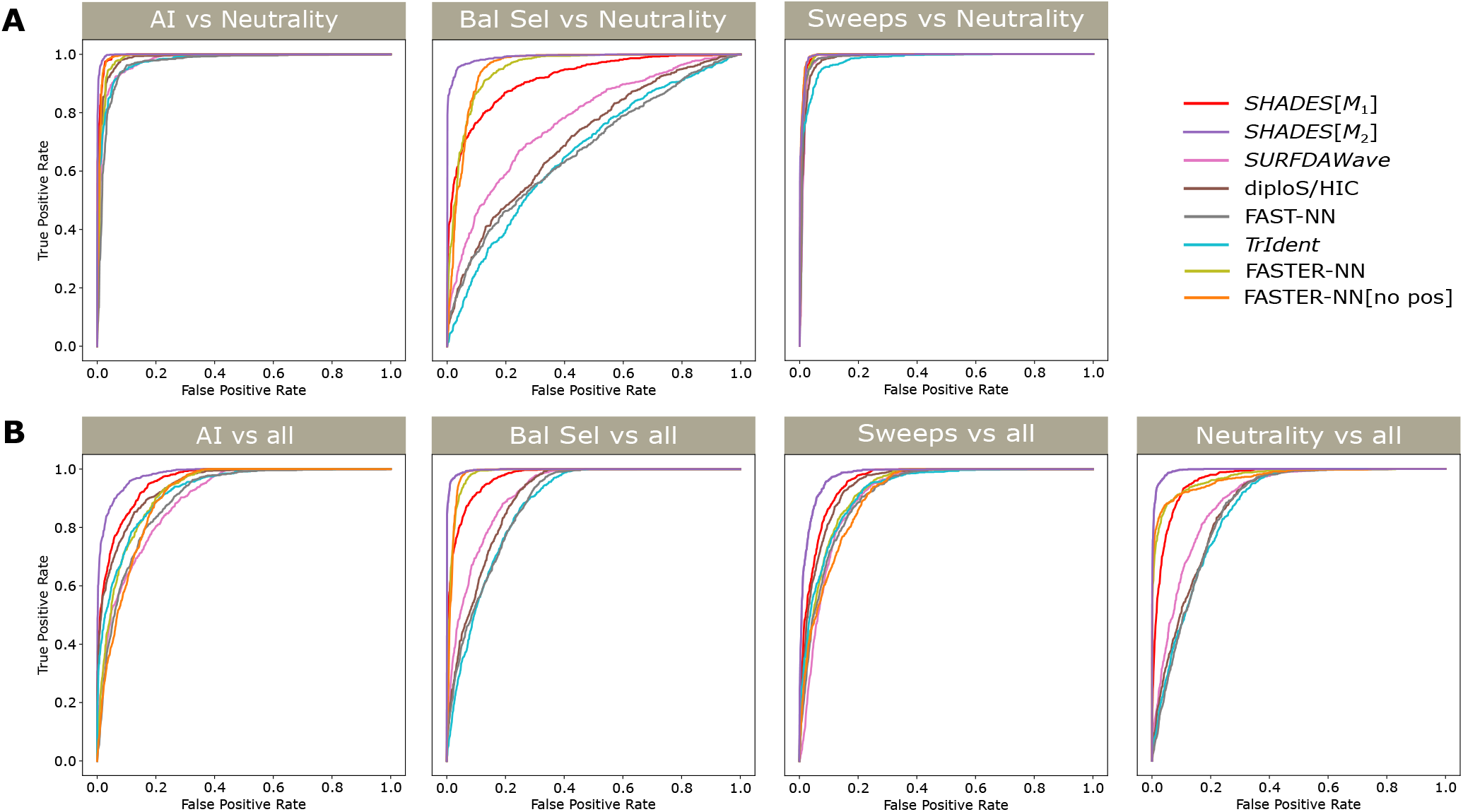
Receiver operating characteristic curves comparing *SHADES* [*M*_1_], *SHADES* [*M*_2_], diploS/HIC, *TrIdent*, *SURFDAWave*, FAST-NN, and FASTER-NN models across two evaluation settings. Here, AI denotes adaptive introgression, and Bal Sel denotes balancing selection. (A) True positive rate for each selective class as a function of the false positive rate from neutrality. (B) True positive rate for each class as a function of the false positive rate computed from all remaining classes.

Different window sizes in the *M*_1_ and *M*_2_ models demonstrate varying strengths and weaknesses in detecting evolutionary classes. By jointly considering outputs from multiple window sizes, the *SHADES* models improve upon the performance of single-window models. Therefore, we selected *SHADES* [*M*_1_] and *SHADES* [*M*_2_] for further analysis. The *M*_2_ architecture contains more trainable parameters than *M*_1_, and accordingly, *SHADES* [*M*_2_] also has more trainable parameters than *SHADES* [*M*_1_]. Shallow *SHADES* [*M*_1_] reaches overall classification accuracy of 80.75% and in contrast, the deeper *M*_2_ architecture enables substantially improved performance, with the stacked *SHADES* [*M*_2_] model achieving an accuracy of 90.85% (Figure 2). Given the markedly higher accuracy of *SHADES* [*M*_2_], we subjected all *SHADES* models to further robustness testing to identify the best-performing model.

To further examine how well the *SHADES* models distinguishes among evolutionary classes, we visualized both the input statistic space and the learned embedding space of *SHADES* [*M*_2_] using multidimensional scaling (MDS) [Kruskal, 1964, Borg and Groenen, 2005] and t-distributed stochastic neighbor embedding (t-SNE) [Van der Maaten and Hinton, 2008] (Figure 4). For the input statistics, we first computed pairwise differences among samples separately for each statistic and window size, normalized these differences to ensure equal contribution from all statistics, and then averaged them to obtain a single sample-by-sample distance matrix. For the learned representation, pairwise differences were computed directly from the penultimate layer embeddings produced by *SHADES* [*M*_2_]. The resulting distance matrices were projected using MDS, with the first two components used for direct visualization and the first 50 components used as input to t-SNE to generate an alternative two-dimensional representation. When applied to the input statistics, both methods showed substantial overlap among classes (particularly high overlap between neutrality and balancing selection and between sweeps and adaptive introgression), indicating that the raw statistical representations do not naturally separate the four evolutionary settings. In contrast, projections based on the *SHADES* [*M*_2_] embeddings exhibited markedly improved class separation, with distinct clustering patterns emerging for neutrality, selective sweeps, adaptive introgression, and balancing selection. Together with the classification accuracy and robustness analyses, these visualizations demonstrate that *SHADES* [*M*_2_] not only achieves high predictive accuracy but also learns an embedding space in which the four evolutionary settings become substantially more separable.

**Figure 4:**
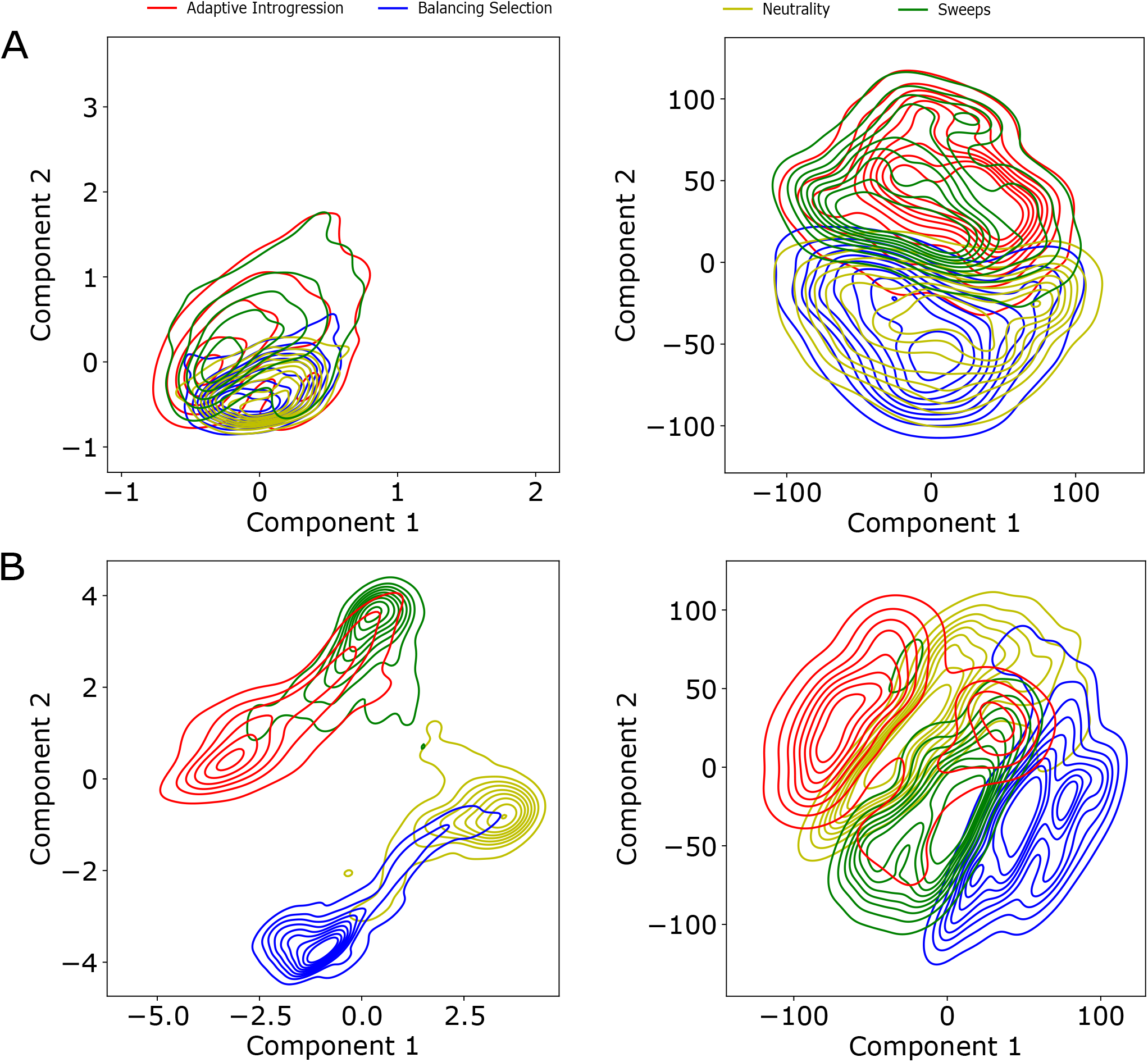
Two-dimensional kernel density visualizations of dimensionality reduction derived from pairwise differences among input image statistics (**I**^(1)^, **I**^(2)^, and **I**^(3)^) of three windows across all test samples. Pairwise differences were normalized, averaged across statistics, and used to construct a sample-by-sample difference matrix, which was then projected by multidimensional scaling (MDS). (A) Contour plot of the first two MDS components (left) and of a two-dimensional t-SNE embedding computed from the first 50 MDS components (right). (B) Contour plots of the first two MDS components (left) and of a two-dimensional t-SNE embedding computed from the first 50 MDS components (right) based on the penultimate-layer outputs of the *SHADES* [*M*_2_] model, representing learned embeddings for the same test samples. Pairwise differences among these embeddings were computed and projected as in panel A. Compared with the raw input statistics, the learned embedding space shows substantially improved separation of the four classes. Scatter plots showing the distributions that the kernel densities approximate can be found in Figure S10.

### Robustness to mutation and recombination rate variation

Detecting natural selection from genomic data requires careful consideration of the inherent variability in mutation and recombination rates across the genome [Kumar and Subramanian, 2002, Bromham, 2011, Smukowski and Noor, 2011, Cutter and Payseur, 2013, Bromham et al., 2015, Peñalba and Wolf, 2020]. Among and within species, these rates are not uniformly distributed across chromosomes and genomic regions. The recombination landscape exhibits a complex distribution, with isolated regions of extremely high recombination activity and regions of low activity, and this influences different patterns of haplotype variation [Petes, 2001, Hey, 2004, Myers et al., 2005, Grey et al., 2009, Booker et al., 2020]. For example, low recombination rates can reduce local haplotypic variation, mimicking the signature of a selective sweep, whereas high recombination rates may elevate variation and obscure sweep signals. Similarly, genomic regions with low mutation rates may appear sweep-like due to reduced variation, whereas regions with high mutation rates may mask historical selective events by inflating haplotypic variation [Harpak et al., 2016, Harris and Pritchard, 2017, Castellano et al., 2020]. Given that recombination and mutation rate heterogeneity is expected to be ubiquitous across genomes, it is essential to conduct comprehensive robustness tests under such settings.

To assess model robustness under mutational and recombination rate heterogeneity, we simulated an additional 1,000 test replicates per class for each of three settings in which genetic parameters are drawn from modified distributions. Specifically, we considered two recombination rate variation settings, with one in which recombination rates were drawn from an exponential distribution with a 10-fold lower mean of 10*^−^*^9^ per site per generation and the other in which the rates were drawn from an exponential distribution with a 10-fold higher mean of 10*^−^*^7^. For the third setting, we introduced mutation rate variability. In particular, instead of fixing the mutation rate to *µ* = 1.25 × 10*^−^*^8^ per site per generation, we sampled the mutation rate uniformly between *µ/*2 and 2*µ*.

For the low recombination rate setting, the *SHADES* [*M*_1_] and *SHADES* [*M*_2_] models achieved test accuracies of 73.65% and 82.25%, respectively (Figure S4A). In particular, the models showed improved detection of balancing selection or lower reduction in accuracy in detecting balancing selection compared to other classes, whereas performance on other classes declined (Figure S4B). Most notably is the drop in the neutral detection rate, which is surprisingly due to an increase in neutral regions misclassified as balancing selection rather than as positive selection (sweeps or adaptive introgression), which may be expected to lead to similar drops in haplotype variation as in low recombination regions. Moreover, sweeps and adaptive introgression are increasingly misclassified as the other under this setting.

In contrast to the low recombination rate scenario, the *SHADES* [*M*_1_] and *SHADES* [*M*_2_] models performed significantly poorer on the high recombination rate setting, dropping to accuracies of 57.60% and 66.33%, respectively (Figure S5A). *SHADES* [*M*_1_] tended to misclassify most cases as balancing selection, whereas *SHADES* [*M*_2_] primarily misclassified them as selective sweeps. The lower accuracies may reflect the breakdown of linkage among nearby variants under high recombination rates, which can weaken informative haplotype patterns. Consequently, misclassifications among the four classes increased for all models in genomic regions with high mean recombination rates.

Finally, under mutation rate variability, the *SHADES* [*M*_1_] and *SHADES* [*M*_2_] models were fairly robust, attaining test accuracies of 80.33% and 90.40%, respectively (Figure S6A). Both *SHADES* models had relatively minor changes in detection rates across classes (Figure S6B). Though both *SHADES* models exhibited similar patterns of vulnerability across varying parameter settings in the simulated data, *SHADES* [*M*_2_] with its ultra deep CNN architecture consistently achieved higher accuracy in all scenarios compared to its shallow architecture counterpart. This consistent performance suggests that *SHADES* [*M*_2_] is the more robust model overall.

### Resilience to missing genomic segments

To evaluate model robustness, we introduced missing data into each test replicate and assessed model performance. Because our models will eventually be applied to empirical data, where regions with low CRG (Centre for Genomic Regulation) scores [Talkowski et al., 2011, Derrien et al., 2012, Mughal et al., 2020] of poor mappability and alignability are removed, we aimed to mimic this setting in the test dataset. Specifically, we randomly selected a region of length two Mb from one of the 22 human autosomes, weighted by SNP density within the autosome. From each selected two Mb region, we identified the relative positions of the five blocks with the lowest CRG scores and removed the corresponding SNPs from the test replicate that intersected those coordinates. After introducing missing data into the test set, we regenerated the three input statistics using the same procedure outlined in the *Input image generation* subsection of the *Methods* and evaluated model performance on this modified dataset.

The classification accuracy of *SHADES* [*M*_1_] and *SHADES* [*M*_2_] remained relatively stable when genomic segments were removed, respectively decreasing by only 1.22% and 1.20% relative to its accuracy on the test set without missing regions (Figure S7). Though these overall reductions were similar in magnitude, the class specific results revealed clearer differences among the models. For *SHADES* [*M*_2_], the classification rates for neutrality and balancing selection remained largely unchanged, while only small decreases were observed for adaptive introgression and selective sweeps. In contrast, *SHADES* [*M*_1_] showed greater variation in classification accuracy across all four classes, with accuracy increasing for only neutrality and decreasing for others after genomic segments were removed. These results indicate that *SHADES* [*M*_2_] provides more stable classifications across evolutionary classes when portions of the genomic inputs are missing and it is the best fit of the three models to discriminate among neutrality, selective sweeps, balancing selection, and adaptive introgression.

### Resilience to background selection

Background selection is another evolutionary phenomenon that is the result of negative selection acting on deleterious alleles, reducing genetic variation at the selected loci and at nearby neutral sites due to linkage disequilibrium [Charlesworth et al., 1993, McVicker et al., 2009, Charlesworth, 2012]. This phenomenon resembles genetic hitchhiking, as seen in selective sweeps, and can lead to allele frequency shifts that mimic signals of positive selection [Charlesworth et al., 1995, Schrider, 2020]. In particular, regions of low recombination are more susceptible to the confounding effects of background selection, as reduced recombination intensifies linkage among sites [Hudson and Kaplan, 1995, Nicolaisen and Desai, 2013]. Though our models were not explicitly trained on data incorporating background selection, it is important to evaluate their robustness under such conditions, given how pervasive this force is within the genome [McVicker et al., 2009, Comeron, 2014].

To evaluate model resilience to background selection, we simulated an additional 1,000 replicates under the CEU demographic history with [Terhorst et al., 2017]. Inspired by the setup from Cheng et al. [2017], each replicate comprised a two megabase sequence containing a centrally located 55 kilobase protein-coding gene, with the remainder of the sequence simulated as neutral. All evolutionary parameters were kept identical to the neutral setting from the original training set, with the addition of deleterious mutations arising inside the gene, which had different proportions of deleterious mutations in exons, introns, and UTRs. The gene was modeled with 50 exons of 100 bases each, 49 introns of one kilobase each, a 5*^′^* untranslated region (UTR) of 200 bases, and a 3*^′^* UTR of 800 bases, approximating mean values from human genome [Mignone et al., 2002, Sakharkar et al., 2004]. Recessive (*h* = 0.1) deleterious mutations arose within the gene with selection coefficient *s* drawn from a gamma distribution with mean of -0.0294 and shape parameter of 0.184 [Boyko et al., 2008, Schrider and Kern, 2017]. The proportions of deleterious mutations were set to 75% in exons, 10% in introns, and 50% in UTRs. Each replicate was evolved for 2 × 10^5^ generations, including a burn-in of 10*N_e_* = 10^5^ generations followed by 10^5^ generations prior to sampling. We sampled *n* = 198 haplotypes (99 diploid individuals) per replicate and applied a scaling factor *λ* = 20 to accelerate simulations.

Under this setting across both *SHADES* models, we find that the proportion of false signals in the background selection replicates was linearly correlated and tracking the *y* = *x* line with the false positive rate of the neutral dataset across all classes. Hence, the distributions of predicted selection probabilities closely follow those of the original neutral test set, with the fraction of false selection signals under background selection comparable to the false positive rate observed from neutral simulations (Figure S8). This agreement indicates that background selection and neutral simulations produced similar rates of false classification as selective sweeps, adaptive introgression, or balancing selection. Thus, under the conditions examined here, background selection did not substantially increase false adaptive classifications for any of the three *SHADES* models.

### Performance on unphased multilocus genotype data

Unphased multilocus genotype (MLG) data inherently contains less information than haplotypes, as it represents a collapsed view of a set of underlying haplotypes without preserving phase information. While phased data retains the arrangement of alleles along individual chromosomes, unphased data obscures this structure, limiting downstream analyses including detection of selection signals. While genetic hitchhiking directly alters haplotypic variation, its effects are only indirectly reflected in unphased multilocus genotype patterns [Harris et al., 2018, Kern and Schrider, 2018, Arnab et al., 2023, 2025]. Moreover, phasing MLGs into haplotypes is often challenging or infeasible for a number of study systems, particularly for those of non-model organisms [Browning and Browning, 2011, O’Connell et al., 2014, Castel et al., 2016, Laver et al., 2016, Loh et al., 2016, Zhang et al., 2017, Sun and Li, 2025]. Because our models were originally trained on simulated haplotype data, we sought to evaluate performance on unphased MLG data to assess generalizability.

To generate unphased MLG data, we combined consecutive pairs of haplotypes within each simulation and treated them as a single unphased MLG representation. Specifically, these MLGs were represented as strings of integers (zero, ones, or two), where each value represents the number of copies of the minor allele in the diploid genotype at a given SNP. We retrained our best-performing model architecture *SHADES* [*M*_2_] using 5,000 replicates per class and evaluated it on 1,000 replicates per class, consistent with the phased haplotype setup. Due to the structural differences between phased haplotype and unphased MLG data, we modified the third summary statistic accordingly. Specifically, instead of using the Hamming distance to compute differences between unique haplotypes, we computed the *L*_1_-norm scaled by two times the window length between the *i*th and (*i* + 1)th most frequent unphased MLGs within each window, maintaining a maximum of *k_max_* = 10 unique unphased MLGs (see *Methods* for descriptions of these quantities). Scaling was applied to the third summary statistic to ensure its magnitude is comparable to that of the Hamming distance used in phased haplotype data.

Despite the inherent limitations of unphased MLG data, the *SHADES* [*M*_2_] model demonstrated strong performance, achieving a test accuracy of 86.55%, which is comparable to its accuracy on phased data (Figure S9A). The ROC curves further illustrate the ability of our *SHADES* [*M*_2_] model to distinguish among evolutionary classes with high power in both the one-versus-all setting (Figure S9B) and the selection-versus-neutrality setting (Figure S9C). Consistent with results from the model trained on phased data, selective sweeps and adaptive introgression exhibit stronger discriminative power compared to balancing selection when distinguishing from neutrality (Figure S9C). This result highlights the robustness of our approach and its potential applicability to datasets lacking phase information.

### Benchmarking classification performance

After evaluating the performance of our models, we benchmarked their classification accuracies and detection powers with five existing methods: *SURFDAWave* [Mughal et al., 2020], diploS/HIC [Kern and Schrider, 2018], *TrIdent* [Arnab et al., 2025], FAST-NN [van den Belt et al., 2024], and FASTER-NN [van den Belt and Alachiotis, 2025]. This comparison allows us to assess how well our approaches perform against published methods that have been demonstrated to be effective for similar classification tasks, with each method offering unique strengths and model inputs. *SURFDAWave* applies functional data analysis to arrays of summary statistics computed across overlapping genomic windows and models spatial autocovariation of summary statistics as a continuous function to differentiate between sweeps and neutrality. In contrast, diploS/HIC transforms summary statistics from genomic windows into image representations for CNN-based classification, effectively capturing spatial relationships among features. Originally designed to distinguish neutrality and selective sweeps (both hard and soft), it demonstrates strong performance in these domains. *TrIdent* employs transfer learning from deep pre-trained CNNs to detect adaptive genomic regions from image-based representations of genomic variation. It utilizes the Inception-ResNet-v2 [Szegedy et al., 2017] architecture pretrained on the ImageNet database as a feature extractor, offering high accuracy in discriminating sweeps from neutrality. FAST-NN is a highly scalable one-dimensional CNN for selective-sweep classification that avoids expensive preprocessing by operating on vectors of derived allele frequencies rather than raw haplotypes or summary statistics. FASTER-NN extends FAST-NN for fine-grained, sliding-window scans by combining allele frequency and positional information with a deeper, dilated convolution architecture designed to increase receptive field and reduce redundant computation across overlapping windows. The complementary strengths of these methods make them well-suited for evaluating the performance of our CNN-based framework.

*SURFDAWave* is a penalized logistic regression (linear) classifier for detecting evolutionary events from genomic summary statistics, and was developed for both binary and multi-class classification. Though originally developed to use nine summary statistics computed across 128 overlapping SNP window each of length 10 SNPs and stride of five SNPs, we expanded it to 256 windows as our simulated sequences are roughly twice as long as those used by Mughal et al. [2020]. These nine summary statistics were constructed to make *SURFDAWave* effective at discriminating between sweeps and neutrality, and to some extent adaptive introgression. However, it still did well in detecting the additional balancing selection class, with the highest accuracy among the four classes (Figure 2). However, the overall classification accuracy of *SURFDAWave* was modest at 67.83% (Figure 2).

Another classifier for detecting evolutionary events with summary statistic inputs is diploS/HIC as well, which uses a multi-branch, multi-layer CNN architecture (nonlinear model). This model computes 12 summary statistics in contiguous physical-sized genomic windows, with the summary statistics chosen to achieve high classification accuracy to distinguish among five classes: neutrality, hard sweeps, soft sweeps, regions linked to hard sweeps, and regions linked to soft sweeps. Though these statistics were computed in 11 windows of 1.1 Mb regions (100 kilobases for each window), we found that using a greater number of windows (21 windows, each spanning approximately 95 kilobases, to cover two megabase regions) permitted diploS/HIC to reach higher accuracy in our setting, and so we retooled the model for our four classes using a greater number of windows. The overall classification accuracy of diploS/HIC of 68.3% is higher than *SURFDAWave*, but significantly lower than *SHADES* [*M*_2_] (Figure 2). Interestingly, despite being developed for sweep detection, diploS/HIC yielded the best performance for adaptive introgression among the published comparison methods (Figure 2).

The *TrIdent* model differs from *SURFDAWave* and diploS/HIC in several ways. The first is that it operates on image representations of sorted haplotype alignments, and automatically extracts features from such images rather than useful features given to the model in the form of summary statistics. It also uses a highly deep pre-trained CNN, compared to the relatively shallow CNN of diploS/HIC, for the feature extractor, and this CNN component does not require training. Finally, it uses penalized logistic regression, like *SURFDAWave*, and is therefore a linear model, though it *TrIdent* was developed for the binary classification problem of distinguishing between sweeps and neutrality. We have retooled it as a multi-class classification, have adjusted the image preprocessing to retain single singletons and doubletons in our calculations for fair comparison with the other models, and we expanded the number of SNPs used to process the input images to the 3,115 central

SNPs for a similar comparison similar to our models. These adjustments to *TrIdent* result in having elevated classification accuracy on our test sets (Figure S11), with an overall classification accuracy of 65.2% (Figure 2), which is also significantly lower than the *SHADES* [*M*_2_] model.

We evaluated FAST-NN and FASTER-NN by adapting both methods from their original binary tasks to our four class setting. In both FAST-NN and FASTER-NN training, we report results from the best performing model, which occurred well before the default maximum epoch as the performance plateaued considerably earlier, suggesting that extended training was unnecessary under our settings. FAST-NN attained an overall test accuracy of 63.2%, the lowest among all compared models and showed the weakest performance for neutrality and adaptive introgression among the compared models. FASTER-NN significantly improved on FAST-NN, reaching an impressive 79.43% test accuracy (Figure 2), but remained well below the *SHADES* [*M*_2_] model. Moreover, FASTER-NN achieved the second highest classification accuracy for balancing selection. Removing pairwise SNP distance information slightly reduced its overall classification accuracy to 77.4% (Figure 2). Despite the reduction in overall accuracy, the modified model achieved a higher classification accuracy for balancing selection but showed a substantial decrease in adaptive introgression. These results suggest that derived allele frequency representations, while scalable and useful to balancing selection, may be less sensitive to adaptive introgression without positional density context.

The *SHADES* [*M*_2_] model outperformed all comparison methods in terms of overall classification accuracy (Figure 2). This advantage is also reflected in the ROC curves (Figure 3), where *SHADES* [*M*_2_] maintains higher true positive rates than competing models across a wide range of false positive rates. In particular, the ROC separation is most pronounced for balancing selection versus neutrality (Figure 3A) and across all one versus rest evaluations (Figure 3B), indicating that the performance gains are not confined to a single confidence threshold. Together, these results show that *SHADES* [*M*_2_] more reliably distinguishes neutrality from adaptive classes, and motivate its application to empirical human genomic datasets, where the data can be more challenging with real world complexities.

### Considering a more complicated demographic history

To evaluate how a more complex demographic setting would influence the performance of *SHADES* [*M*_2_] and other comparison methods, we added gene flow with an unsampled population to inject population structure into the demographic model. Specifically, we added a second population of size 10,000 diploid individuals that split from the sampled (CEU) population approximately 5,600 generations before sampling [Gutenkunst et al., 2009]. The two populations exchanged migrants at a rate of 2.5 × 10*^−^*^4^ per generation [Gutenkunst et al., 2009], while all other simulation parameters followed those used for the CEU demographic model. Using this demographic model, we trained and evaluated *SHADES* [*M*_2_] and each benchmarking method following the same procedure used for the CEU demographic model.

Among the evaluated methods, *SHADES* [*M*_2_] achieved the highest overall classification accuracy at 83.13% (Figure S12). Though this accuracy was 7.72% lower than that obtained under the CEU demographic model, which is as expected, *SHADES* [*M*_2_] maintained strong performance under a more complex demographic history incorporating population structure and continuous migration. The overall classification accuracies of *TrIdent*, diploS/HIC, *SURFDAWave*, FAST-NN, and FASTER-NN were 65.13%, 66.13%, 63.05%, 62.10%, and 77.73%, respectively, and all bench-marking methods showed lower accuracy than under the CEU demographic model, as expected. The patterns of misclassification were also consistent with those observed under the CEU model, with methods frequently confusing selective sweeps with adaptive introgression and balancing selection with neutrality, with a notable increase in balancing selection detection rate for FASTER-NN of about 7%. Interestingly, the reduction in overall accuracy for *SHADES* [*M*_2_] and *SURFDAWave* were distributed relatively evenly across classes. In contrast, the smaller changes in overall accuracy for the remaining methods concealed class specific tradeoffs, with gains in some classes offset by reductions in others. Thus, despite its larger reduction in overall accuracy relative to some bench-marking methods, *SHADES* [*M*_2_] retained the highest overall accuracy and comparatively balanced performance across evolutionary classes.

### Footprints in human genomes

To ensure the applicability of *SHADES* [*M*_2_] to real, non-synthetic data, we next applied the trained *SHADES* [*M*_2_] (denoted *SHADES* in this subsection for brevity) model to empirical data to a European human population (CEU) from the 1000 Genomes Project dataset [The 1000 Genomes Project Consortium, 2015]. In particular, we computed the same three statistical representations for each of the 22 human autosomes, with these computations were performed across 3,115 contiguous SNPs using a sliding window approach with a stride of 21 SNPs (see *Input Image Generation* subsection of the *Methods*), yielding 183,558 partially overlapping windows genome-wide. For each window, *SHADES* assigned a confidence score to each evolutionary class as its class probability, and these scores were subsequently used to identify and prioritize genomic regions with candidate signatures of selective sweeps, long-term balancing selection, and adaptive introgression. The genome-wide distributions of all window-and gene-level confidence scores were strongly concentrated near zero for selective sweeps, balancing selection, and adaptive introgression (Figure S14). Only a small subset of windows and genes received confidence scores greater than 0.9 for a given non-neutral class, yielding a relatively small number of windows and genes across the empirical genome scan with well-supported evidence of adaptation (Table S1).

#### Selective sweeps

We examined high-confidence protein-coding genes to characterize individual candidate loci contributing to the inferred selection signals. As expected, *SHADES* identified high sweep support at the well-established region on chromosome 2 that is involved in the lactase persistence phenotype in humans [Bersaglieri et al., 2004, Sabeti et al., 2006, Tishkoff et al., 2007], successfully recapitulating this classical selective sweep target [Bersaglieri et al., 2004, Gerbault et al., 2009]. The core of this signal centered on *LCT* (confidence = 0.808), its antisense RNA *LCT-AS1* (0.808), and its upstream neighbor *MCM6* (0.878). The *LCT* gene codes for the lactase enzyme that hydrolyzes lactose and its continued expression into adulthood represents one of the most extensively documented examples of local adaptation [Boll et al., 1991, Itan et al., 2010]. Upstream of this locus, *MCM6* harbors an enhancer that acts as a regulatory mechanism to control *LCT* expression [Labrie et al., 2016, Anguita-Ruiz et al., 2020]. The development of animal domestication and dairying created cultural and dietary conditions that favored the lactase-persistence phenotype [Scrimshaw and Murray, 1988, Ségurel and Bon, 2017]. Because of this selective pressure, it generated a hitchhiking footprint that depleted genetic variation across the surrounding genomic region. Accordingly, *SHADES* detected high-confidence sweep signals radiating across nearly a megabase of sequence, capturing both upstream and downstream neighboring genes. Downstream of the *LCT* target locus, *SHADES* identified a suite of candidate genes with strong sweep support, including *UBXN4* (0.986), *R3HDM1* (0.886), *ZRANB3* (0.996), and *RAB3GAP1* (0.832). Notably, *ZRANB3* has previously been highlighted as a target with high sweep support that helps repair DNA lesions during DNA replication [Weston et al., 2012, Liu et al., 2013, Sebesta et al., 2017, Amin et al., 2024]. Upstream of *LCT*, the extended sweep signature also detected *DARS1* (0.865). The simultaneous identification of *LCT* and neighboring genes is consistent with the broad linkage disequilibrium and lost haplotypic variation expected under a strong, recent selective sweep (Figure S15).

Beyond the *LCT* region, *SHADES* uncovered several additional candidates that have previously been implicated in studies of human adaptation. Notably, *OCA2*, which is a major determinant of pigmentation and eye color variation in Europeans, was classified as a sweep candidate with a high confidence score of 0.99 (Figure S15). The *OCA2* /*HERC2* region has repeatedly been highlighted in genome-wide scans of positive selection, serving as an additional positive control in European populations [Eiberg et al., 2008, Sturm et al., 2008]. Furthermore, *SHADES* detected *ABCA12*, a gene essential to skin barrier function and lipid transport, which has been previously linked to environmental ultraviolet exposure adaptation in Eurasians [Akiyama, 2014, Colonna et al., 2014, Arnab et al., 2026]. Additional sweep candidates included *SERINC2*, which has been associated with alcohol dependence in Europeans [Zuo et al., 2013, Arnab et al., 2025], and *PCDH15*, which encodes a protein critical for sensory function in the retina and cochlea where mutations are known to cause Usher syndrome type 1F [Ahmed et al., 2001, Alagramam et al., 2001, Sabeti et al., 2007, Qiu et al., 2022]. Putative sweep signatures were also found in *SIGLEC10* and *SIGLEC12*, which are crucial components of human sialic acid biology and have been identified as targets of recent selective sweeps [Nielsen et al., 2009, Angata, 2018, Moon et al., 2018]. Other high-confidence candidates included *BCAS3* and *PDPN* [Sabeti et al., 2007, Arnab et al., 2023]. *PDPN* acts as a marker for lymphatic vessels and is implicated in cancer metastasis [Kitano et al., 2010]. The *BCAS3* gene drives angiogenesis and directional endothelial cell migration [Köttgen et al., 2010, Jain et al., 2012]. It is also described as a biomarker of breast cancer [Gururaj et al., 2006]. The detection of cancer-associated genes like *BCAS3* and *PDPN* aligns with a recurring pattern in recent machine learning scans, which suggests that alleles historically targeted by positive selection may contribute modern carcinogenesis [Lou et al., 2014, Schrider and Kern, 2017, Mughal and DeGiorgio, 2019, Amin et al., 2023, Arnab et al., 2023].

We also identified a set of novel selective sweep candidates from our scan. Specifically, *SHADES* identified *IRF2* which is a key mediator of Type I interferon responses [Harada et al., 1989]. Pathogen-driven selection has recurrently shaped genes involved in type I interferon responses during primate evolution, providing a broader functional context for *IRF2* as a candidate of selective sweep [Judd et al., 2021]. Furthermore, *SHADES* detected pronounced signatures of selection at the *PADI1* gene, which encodes an enzyme that modifies filaggrin and keratin K1 during epidermal differentiation, supporting skin hydration and barrier function [Nachat et al., 2005, Méchin et al., 2005]. Previous evidence of selective sweeps affecting epidermal regulatory elements in European populations indicates that skin barrier function has been a target of human adaptation [Ju and Mathieson, 2021, Mathyer et al., 2021]. These observations provide a possible functional context for selective selection at *PADI1*. The predicted selective sweep probabilities as a function of genomic position around a subset of the candidate genes discussed are shown in Figures 5 and S15.

**Figure 5:**
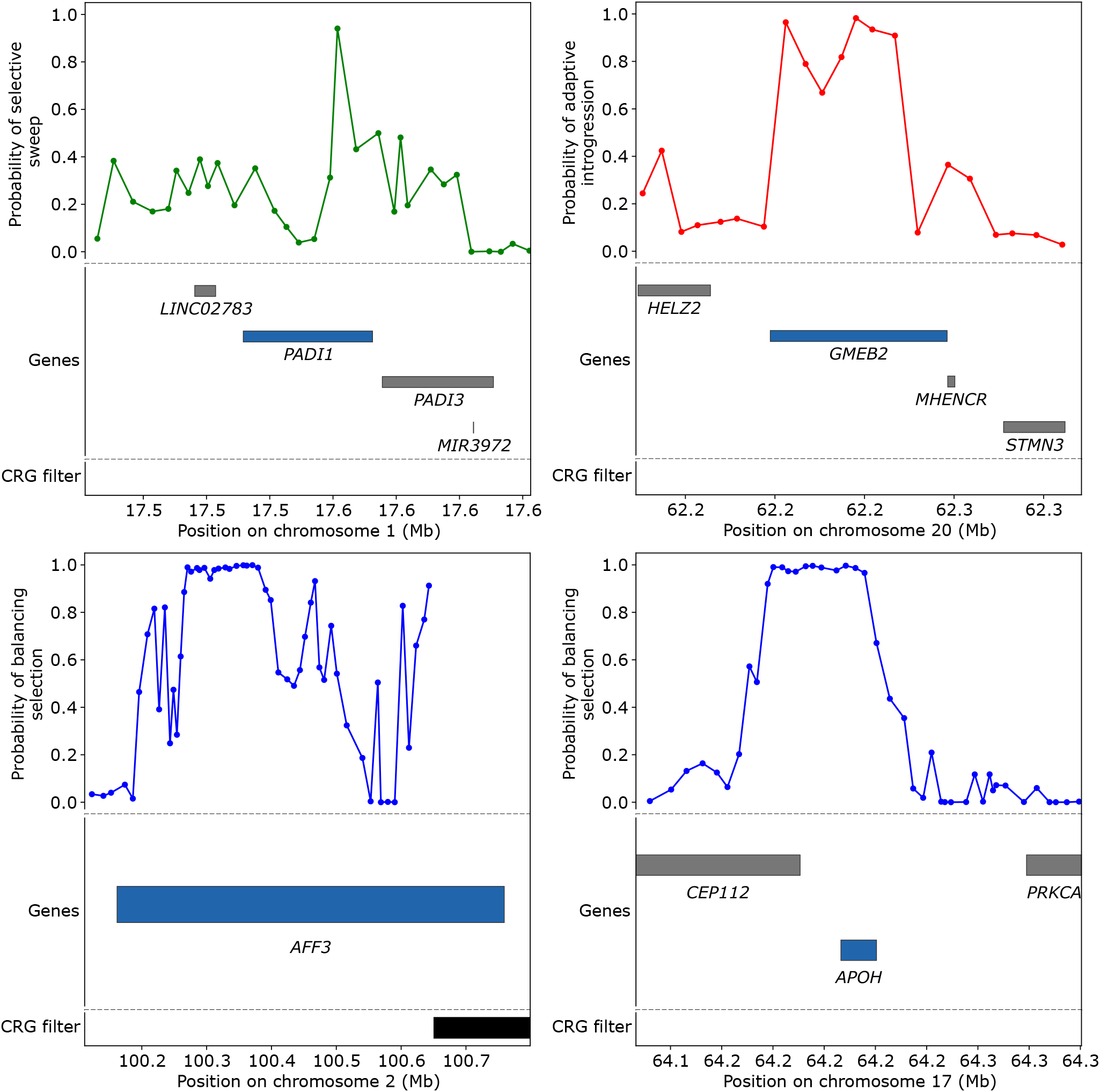
Genomic spatial pattern of *SHADES* [*M*_2_] predicted probabilities for selective sweeps (green curve), balancing selection (blue curves), and adaptive introgression (red curve) across a set of highlighted candidate genes within the empirical CEU scan. Protein coding genes and their genomic coordinates overlapping each highlighted region are shown below the predicted probabilities as horizontal gray bars, with the focal gene at which the signal is centered shown as a horizontal blue bar. CRG filtered intervals are displayed at the bottom as horizontal black bars to indicate genomic segments containing filtered or missing data.

#### Balancing selection

Beyond the selective sweep candidates described above, *SHADES* also identified genomic regions with signatures of long-term balancing selection. We recovered a dense cluster of candidates within and flanking the classical major histocompatibility complex (MHC) region of chromosome 6, consistent with previous reports of balancing selection in this region [Andŕes et al., 2009, Bitarello et al., 2018]. This cluster included classical and non-classical *HLA* loci (*HLA-F, HLA-H, HLA-J, HLA-L, HLA-DQA2, HLA-DQB2, HLA-F-AS1*), MHC-linked lncRNAs (*HCG17, HCG18, HCG22, HCG27*), butyrophilin genes (*BTN2A1, BTN3A1, BTN3A3*), a cluster of TRIM genes (*TRIM10, TRIM15, TRIM26, TRIM31, TRIM38, TRIM39, TRIM40*), and PSORS1-region genes (*CDSN, PSORS1C1, PSORS1C2, PSORS1C3, CCHCR1, TCF19*) (Figure 6). Many of these loci play critical, independent roles in immune defense and autoimmunity. For example, *CDSN* is essential for maintaining structural integrity in the skin barrier [Oji et al., 2010], a trait hypothesized to reflect an evolutionary trade-off between cutaneous pathogen resistance and psoriasis susceptibility [Kere, 2005, Andŕes et al., 2009, Cagliani et al., 2011, Lee and McGrath, 2021]. Similarly, *PSORS1C1* has been associated with susceptibility to psoriasis and systemic sclerosis [Allanore et al., 2011] and TRIM family members (e.g., *TRIM10* and *TRIM40*) regulate innate antiviral responses [Zhao et al., 2017, Shen et al., 2021, Kong et al., 2023] and exhibit strong signatures of balancing selection [Cagliani et al., 2011]. Furthermore, non-classical genes such as *HLA-DQA2* and *HLA-DQB2* contribute to broader antigen presentation pathways [Lenormand et al., 2012] and demonstrate established balancing selection footprints [Bitarello et al., 2018]. The robust recovery of this extended MHC signal serves as an internal positive control, demonstrating high sensitivity of *SHADES* in detecting complex, biologically validated targets for balancing selection.

**Figure 6:**
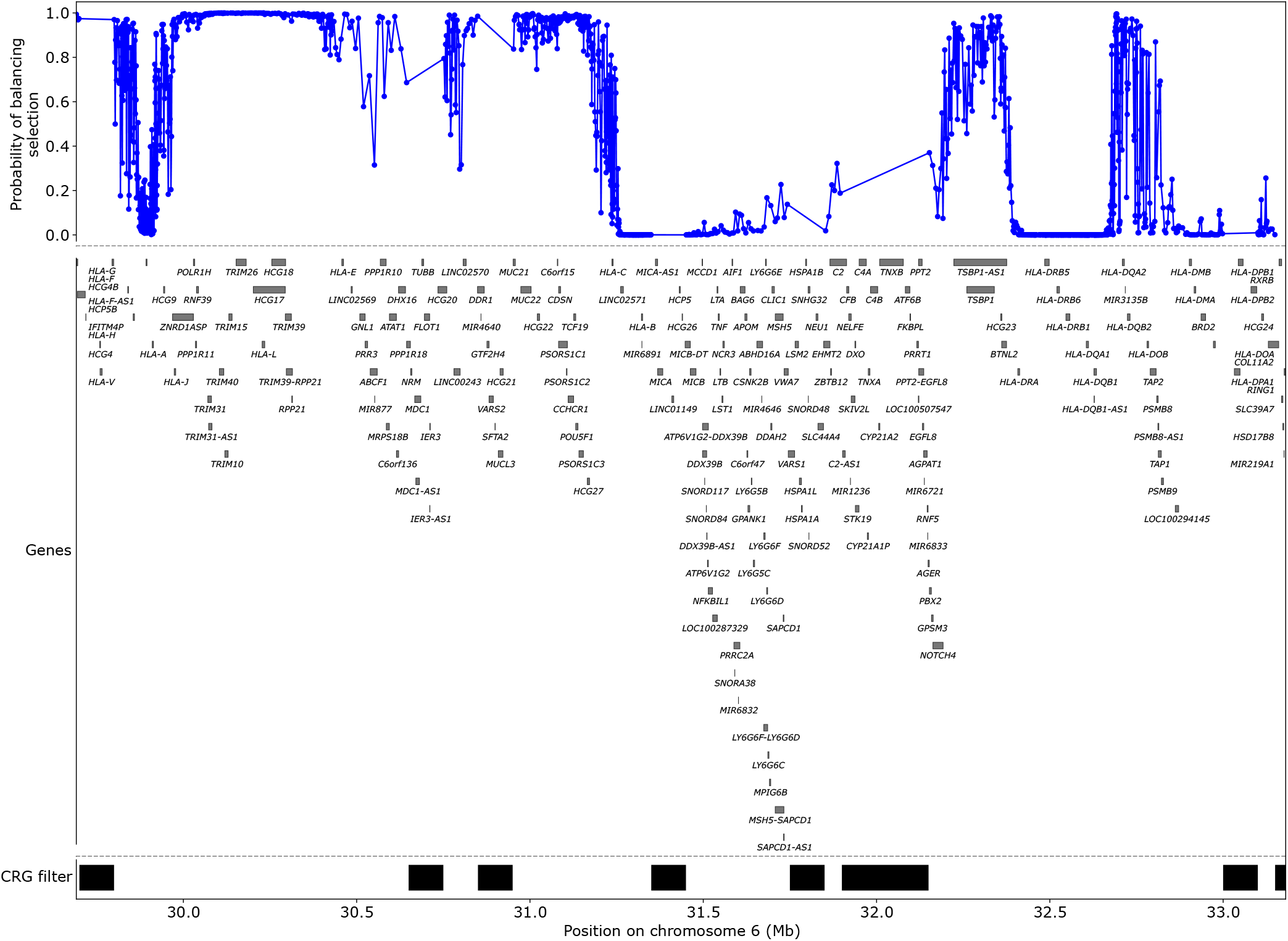
Genomic spatial pattern of *SHADES* [*M*_2_] predicted balancing selection probabilities across the major histocompatibility complex (MHC) region on chromosome 6 within the empirical CEU scan. Protein coding genes and their genomic coordinates overlapping the region are shown below the predicted probability as horizontal gray bars. CRG filtered intervals are displayed at the bottom as horizontal black bars to indicate genomic segments containing filtered or missing data.

The genome-wide scan also recovered several previously reported genes with strong support for balancing selection outside the extended MHC region. Prominent among these is *IL37*, a critical anti-inflammatory cytokine [Nold-Petry et al., 2015, Su and Tao, 2021] where ancient balanced haplotypes are estimated to have been maintained since before the divergence of modern and archaic humans to modulate innate immune responses [Nold et al., 2010, Kang et al., 2015]. *SHADES* detected *CADM2*, which has also been found as a balancing selection candidate in several genomic scans [DeGiorgio et al., 2014, Siewert and Voight, 2017, Cheng and DeGiorgio, 2020]. *CADM2* is a synaptic cell-adhesion molecule frequently implicated in behavioral traits [Boutwell et al., 2017, Sanchez-Roige et al., 2023, Chen et al., 2024]. In addition, neuropathogenic measles virus strains use *CADM2* as a neuronal receptor to trigger membrane fusion, facilitate spread between neurons, and promote neurovirulence [Shirogane et al., 2021, Takemoto et al., 2023]. The exploitation of cell-adhesion proteins by viruses may impose competing pressures to preserve their cellular functions while limiting viral infection, providing a plausible evolutionary context for the maintenance of variation at *CADM2* [Wang et al., 2020]. Furthermore, our candidate list intersects with the top previously-identified empirical candidates [DeGiorgio et al., 2014], such as *STPG2*, a poorly characterized gene encoding sperm-tail PG-rich repeat-containing protein 2 that is expressed in testis [Uhlén et al., 2015], *ANK3*, which organizes axonal membranes [Kizhatil et al., 2007, Huang and Rasband, 2018], alongside broadly acting loci like the tumor suppressor *LRP1B* [Liu et al., 2000, 2001, Bu et al., 2006], the structural ciliary protein *AXDND1* [Houston et al., 2024], and the cell-cell adhesion regulator *CTNNA3* [Janssens et al., 2001, Fanjul-Fernández et al., 2013]. Our scan also recovered several other genes reported in the same study, including *CEP112*, *SPEF2*, *SH3RF3*, *ART3*, *STK32A*, and *ZNF568*. The overlap with previously reported empirical candidates identifies these genes as potentially informative targets for further investigation of balancing selection.

In addition to previously reported genes, *SHADES* identified several novel candidates with high balancing selection confidence scores. *SHADES* detected *APOH*, a glycoprotein that facilitates the clearance of viral and bacterial pathogens [Nilsson et al., 2008, Laird et al., 2014] while simultaneously acting as the primary autoantigen in antiphospholipid syndrome [Schwarzenbacher et al., 1999, Sikara et al., 2010]. The involvement of *APOH* in both pathogen clearance and autoantigen recognition suggests a potential trade-off between host defense and self-tolerance [Leffler et al., 2013, Brinkworth and Barreiro, 2014]. Another candidate, *AFF3*, facilitates immunoglobulin class-switch recombination and contributes to antibody-mediated resistance against *Plasmodium* infection [Tsukumo et al., 2022]. Conversely, genetic variation at *AFF3* has been associated with susceptibility to autoimmune diseases, including rheumatoid arthritis and type 1 diabetes [Barton et al., 2009, Tsukumo et al., 2022]. Balancing selection has been implicated in maintaining variation at other loci associated with autoimmune disease, suggesting a broader evolutionary connection between immune diversity and autoimmune susceptibility [Cagliani et al., 2011, Azevedo et al., 2015]. Whether this mechanism explains the signal at *AFF3* remains unknown. The predicted balancing selection probabilities as a function of genomic position around a subset of the previously reported and novel candidate genes discussed are shown in Figures 5 and S16.

#### Adaptive introgression

*SHADES* recovered several genes previously reported as adaptive introgression, including *STAT1*, *IL22RA2*, *SLC28A1*, *ESRRB*, and *PRKACA*. Among these genes, *STAT1* and *IL22RA2* contribute to immune signaling [Levy and Darnell Jr, 2002, Khawar et al., 2016] and host defense [Sikorski et al., 2011, Mühl and Bachmann, 2019], whereas *SLC28A1*, *ESRRB*, and *PRKACA* have roles in nucleoside transport [Mackey et al., 1999], transcriptional regulation [Zhang et al., 2008], and hepatocellular carcinoma [Honeyman et al., 2014], respectively. We also identified *MUC4*, *CDH13*, *KCNE4*, and *ITSN2*, which were previously reported as candidates for adaptive introgression in Europeans [Setter et al., 2020]. These genes contribute to epithelial function [Vancamelbeke et al., 2017], cell adhesion [Qian et al., 2007], ion channel regulation [Abbott, 2016], and endocytosis [Pucharcos et al., 2000], respectively. The functional diversity of these genes is consistent with previous evidence that archaic introgression was involved in adaptation to pathogen exposure and other environmental pressures [Gouy and Excoffier, 2020]. We also found *GMEB2* which encodes a transcription factor that binds regulatory DNA elements to modulate gene expression [Kaul et al., 2000]. Interestingly, adaptive introgression signals at the *GMEB2* gene have been associated with autoimmune disorders [Reilly et al., 2022]. In addition, *BNC2* was detected with a moderate confidence score of 0.63. This pigmentation associated gene contains Neanderthal derived variation in European populations and may be relevant to phenotypic adaptation following the expansion of modern humans into Eurasia [Dannemann and Kelso, 2017, Yuan et al., 2021].

*SHADES* identified several novel gene candidates for adaptive introgression as well. Among the novel adaptive introgression candidates, *CRISPLD2* regulates pulmonary inflammatory responses to bacterial lipopolysaccharide [Zhang et al., 2016], whereas *IL15RA* contribute to T-cell, natural killer cell, and antiviral immunity [Verbist and Klonowski, 2012]. These immune functions are consistent with established cases in which Neanderthal introgression contributed functional variation to microbial and antiviral immune pathways [Dannemann et al., 2016, Sams et al., 2016]. Moreover, likely adaptive Neanderthal-derived segments in Europeans are enriched for genes encoding proteins that interact with RNA viruses [Enard and Petrov, 2018]. These established examples provide a functional precedent for interpreting the novel immune candidates detected by *SHADES*, though the specific pathogens and selected phenotypes underlying these signals remain unresolved. The predicted adaptive introgression probabilities as a function of genomic position around a subset of the previously reported and novel candidate genes discussed are shown in Figures 5 and S17.

#### Gene Ontology analysis

We performed Gene Ontology (GO) enrichment analyses on *SHADES* inference results using both ranked gene lists and two unranked gene lists with *GOrilla* [Eden et al., 2007, 2009]. For each ranked list analysis, genes were ordered by decreasing *SHADES* confidence score for the corresponding selection class. *GOrilla* then identifies GO terms significantly enriched toward the top of the ranked gene list [Eden et al., 2009]. For each unranked analysis, the target list consisted of genes with a confidence score greater than 0.9 for the corresponding selection class, whereas the background list included all genes evaluated by *SHADES*. Thus, the ranked analyses used the relative ordering of all scored genes without requiring a predefined confidence threshold, whereas the unranked analyses tested whether GO terms were overrepresented among the predefined high-confidence genes relative to the background. We evaluated all three ontologies, which describe gene products according to their associated biological processes, molecular functions, and cellular components. GO terms with false discovery rate adjusted *q*-values less than 0.05 were considered significantly enriched (Figure 7). For adaptive introgression, GO enrichment analyses of the ranked and unranked analyses consistently highlighted pathways associated with xenobiotic metabolism, detoxification, and immune function (Tables S2-S5). The strongest signal was driven by the UGT1A gene cluster, which showed significant enrichment for biological processes related to flavonoid metabolism, xenobiotic glucuronidation, cellular glucuronidation, and glucuronate metabolism (Tables S4 and S5). The repeated identification of glucuronidation-related processes across both ranked and unranked analyses suggests that introgressed variants may have contributed to metabolic adaptation through the processing of dietary and environmental compounds. Enrichment of immune-related categories (Table S3) further suggests a role for adaptive introgression in shaping host defense and pathogen-response pathways. In contrast, GO analyses of balancing selection for both ranked and unranked analyses revealed highly consistent enrichment of pathways related to keratinization and intermediate filament organization, together with enrichment of interleukin-1 receptor binding (Tables S6-S10). These findings suggest that balancing selection may have preferentially maintained variation in biological functions associated with epithelial barrier integrity, skin and hair phenotypes, and immune regulation. In contrast, neither the ranked nor the unranked GO analyses for selective sweeps identified significantly enriched terms at the FDR-adjusted *q*-value threshold of 0.05, indicating that the genes receiving sweep support were not significantly concentrated within particular biological processes, molecular functions, or cellular components under our analysis.

**Figure 7:**
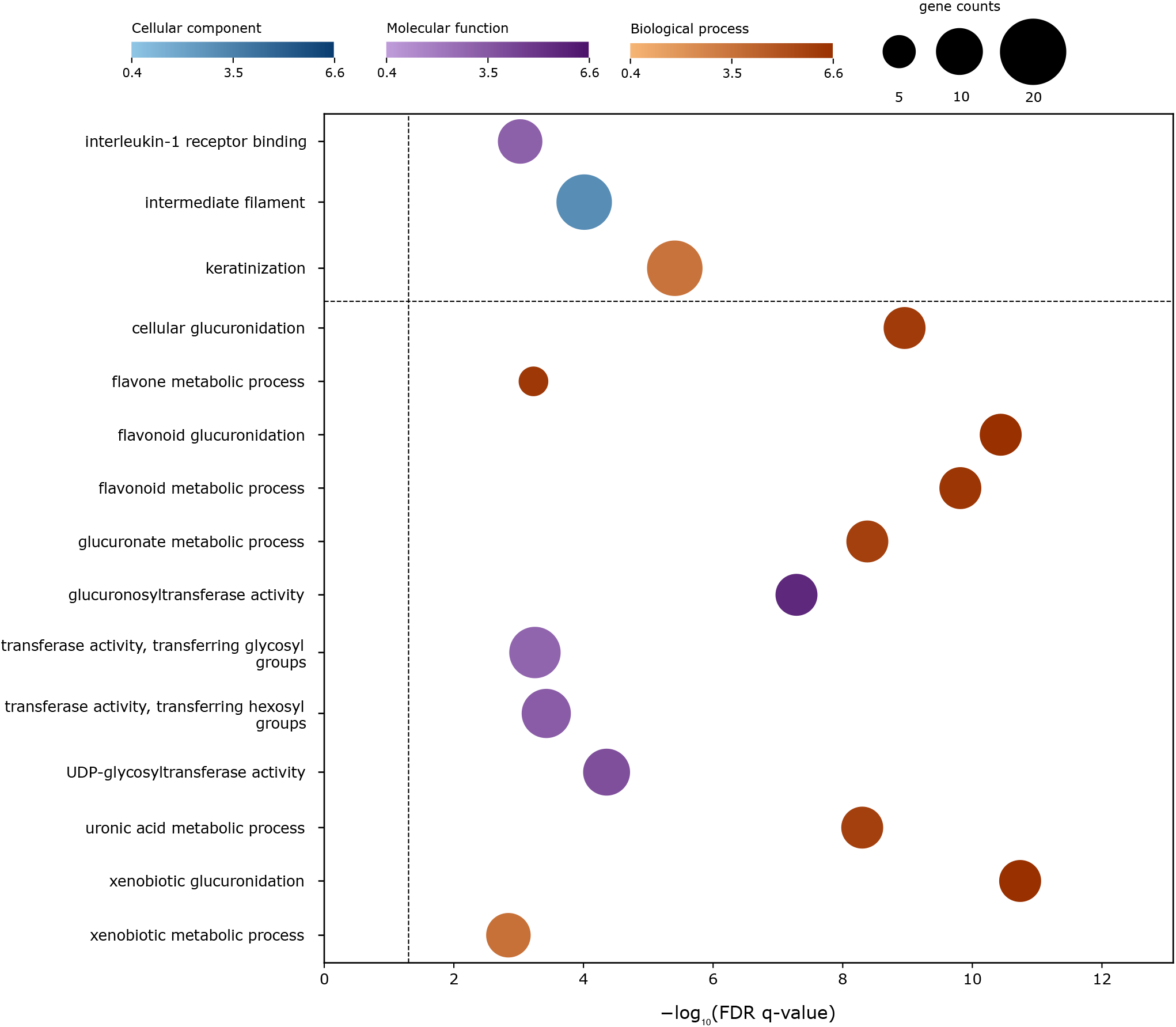
Gene Ontology (GO) enrichment among genes associated with balancing selection and adaptive introgression. The top 15 GO terms were selected by comparing FDR *q*-values from all *GOrilla* analyses of adaptive introgression, balancing selection, and selective sweeps, including both the ranked list and paired unranked list analyses. For each term, the result with the lower *q*-value was retained and all displayed results originated from ranked-list analyses. Terms are ordered alphabetically within each gene set, with balancing selection terms above the horizontal dashed line and adaptive introgression terms below, with no selective sweep terms provide as that class did not yield significant GO terms at FDR *q*-value below 0.05. The *y*-axis lists GO term descriptions, and the *x*-axis shows log_10_(FDR *q*-value), with larger values indicating stronger statistical significance. The vertical dashed line marks FDR *q* = 0.05. Each bubble represents a GO enrichment result, with area proportional to the number of total genes in the chosen ranked GO term. Blue, purple, and orange colors indicate cellular component, molecular function, and biological process, respectively. Within each color gradient, darker shades indicate higher log_2_(enrichment). Complete enrichment results and corresponding gene lists are provided in Tables S2-S10.

## Discussion

We introduced *SHADES*, a CNN-based framework for distinguishing selective sweeps, balancing selection, adaptive introgression, and neutrality based on images summarizing haplotype variation. By integrating haplotype frequencies, changes in haplotype frequency distributions, and haplotype differences, the models capture both the relative abundance of haplotypes and the sequence divergence among them. To capture evolutionary signatures at multiple genomic scales, *SHADES* evaluates these quantities at several window sizes. Smaller windows were chosen to capture finer-scale haplotypic variation that may help identify balancing selection, whereas larger windows summarize variation across broader regions that may help identify sweeps and adaptive introgression. The images summarize these statistics across many SNPs to retain the surrounding genomic context. While single window approaches are susceptible to localized noise, the stacked multiscale *SHADES* models integrate learned representations across window widths, significantly improving overall classification accuracy and detection power relative to models trained at a single scale.

The two *SHADES* architectures offer different tradeoffs between classification accuracy and computational requirements. The shallow *SHADES* [*M*_1_] architecture contains relatively few trainable parameters and can therefore be trained using modest computational resources. In contrast, *SHADES* [*M*_2_] uses the substantially larger Inception-ResNet-v2 architecture, providing greater representational capacity at a higher computational cost. To mitigate the extensive training time and data requirements typically associated with this larger model, we initialized *SHADES* [*M*_2_] with weights pretrained on ImageNet. This transfer learning approach allowed the model to reach its best validation result within a relatively small number of fine-tuning epochs.

An important consideration for any selection detection framework is robustness to empirical genomic confounders that can mimic or obscure evolutionary signatures. Despite being trained with a fixed mutation rate, when evaluated against variable mutation rates, which can easily alter baseline haplotypic variation, *SHADES* [*M*_2_] demonstrated exceptional stability. The model also remained highly effective when subjected to missing genomic segments modeled after filtering strategies to remove empirical regions with poor mappability and alignability, which can reduce local haplotype variation and mislead selection detectors into generating false positives. Furthermore, all three *SHADES* architectures were robust to background selection under the scenario examined here, as background selection did not substantially increase false classifications as selective sweeps, adaptive introgression, or balancing selection compared to false positive rate under neutrality. Recombination rate heterogeneity presented a greater challenge than other cases. Elevated mean recombination rates rapidly eroded the haplotypic footprints of past selection, causing all evaluated models to struggle with detection. Conversely, in low mean recombination settings, *SHADES* [*M*_2_] experienced a general reduction in accuracy across all classes but still maintained a higher overall performance than the other benchmarking methods. Finally, while *SHADES* [*M*_2_] successfully analyzes both phased and unphased genomic data using identical inputs, the inherent loss of haplotype phase information in unphased multilocus genotypes results in modest reductions in classification accuracy. We further evaluated the classification methods under a more complex demographic model that incorporated population structure through gene flow with an unsampled population. Though the overall classification accuracy of *SHADES* [*M*_2_] decreased by 7.72% relative to its accuracy under the CEU demographic model, it remained higher than the evaluated benchmarking methods and showed a comparatively balanced reduction across evolutionary classes, whereas several benchmarking methods gained accuracy for some classes at the expense of others. This reduction in accuracy may be attributable to increased overlap among the evolutionary classes, as this demographic scenario is more complex than the CEU model and blurs class boundaries. Together, these results indicate that the multiscale haplotype based representation in *SHADES* provides robustness to several empirical genomic confounders, experiences a modest reduction in accuracy with the loss of haplotype phase information, and retains strong comparative performance under the more complex demographic scenario examined here.

Application of *SHADES* to European human genomic data recovered several well documented selection signals, including sweep support at the region containing *LCT* and *MCM6* and at the *OCA2* gene, widespread balancing selection support among genes in the MHC region, and adaptive introgression support at *BNC2* gene. The recovery of these established genes demonstrates that *SHADES* can identify empirical genomic patterns associated with distinct evolutionary processes. Beyond these well established examples, the previously reported empirical candidates recovered by *SHADES* revealed distinct biological themes. Candidate sweep genes included genes involved in tissue remodeling, angiogenesis, and cancer related processes, whereas balancing selection candidates were enriched for immune regulation and host-pathogen interactions. Adaptive introgression candidates spanned a broad array of functions including immune signaling, epithelial biology, cellular regulation, and pigmentation. Collectively, these findings are consistent with growing evidence that adaptation in human populations has frequently involved interactions with pathogens, environmental challenges, and physiological interfaces between the organism and its environment.

In addition to recovering previously reported candidates, *SHADES* prioritized several novel genes by using a stringent confidence score threshold of 0.9. Though these novel candidates span different evolutionary classes, many are associated with immune defense, inflammatory regulation, or epithelial barrier biology. Specifically, *IRF2*, *APOH*, *AFF3*, *CRISPLD2*, and *IL15RA* have roles in immune and inflammatory processes, whereas *PADI1* contributes to epidermal barrier function. These recurring functional themes parallel those observed among previously reported candidates and suggest that host-environment interactions may represent a common feature of several signals identified by *SHADES*.

## Methods

We have developed four different convolutional neural network (CNN) models to discriminate among evolutionary processes when scanning genomes for adaptive events, specifically focusing on distinguishing among neutrality, sweeps, adaptive introgression, and long-term balancing selection. For ease of reference, we denote these models as *M*_1_, *M*_2_ and *SHADES* [*M*] with *M* ∈ {*M*_1_*, M*_2_}, and the input for each model consists of three images summarizing haplotype variation across a stretch of a chromosome. In particular, the first image depicts haplotype frequencies across overlapping windows, the second image represents probabilistic distances among the distribution of haplotype frequencies between these genomic windows, and the third image portrays nucleotide differences between haplotypes within a window ranked by their frequencies. In this section, we describe data processing and generation, network architectures, and model training.

### Simulation protocol

For the purpose of training CNN models with images summarizing haplotype variation, we calculated these statistics from simulated genomic data using the forward-time simulator SLiM 3 [Haller and Messer, 2019]. For each of the four classes (neutral, sweep, adaptive introgression, and balancing selection), we generated 6,000 replicates under a European (CEU) human demographic history [Terhorst et al., 2017], with per-site per-generation mutation rate *µ* = 1.25 × 10*^−^*^8^ [Scally and Durbin, 2012] and per-site per-generation recombination rate *r* drawn from an exponential distribution with mean 10*^−^*^8^ and truncated at three times the mean [Payseur and Nachman, 2000, Schrider and Kern, 2017]. We ran each simulated replicate for 2 × 10^5^ generations, which included a burn-in period of 10*N_e_* = 10^5^ generations, followed by an additional 10^5^ generations of evolution prior to sampling. To match empirical CEU data from the 1000 Genomes Project [The 1000 Genomes Project Consortium, 2015], we sampled *n* = 198 haplotypes to create *n/*2 = 99 diploid individuals, where these haplotypes derived from two Mb-long sequences. As is common with forward-time simulations, we applied a scaling factor of *λ* = 20 for speedup [Yuan et al., 2012, Ruths and Nakhleh, 2013].

For simulations of sweeps, adaptive introgression, and balancing selection, we introduced beneficial mutations at the center of simulated sequences (one Mb) with per-generation selection strength *s* ∈ [0.005, 0.5] drawn uniformly at random. For sweeps and adaptive introgression replicates, we introduced an initially neutral mutation at the center of a simulated sequence at a time drawn uniformly at random from 3,000 to 1,020 generations prior to sampling. We then allowed the simulation to run for 40 generations. From this point onward, if the standing allele frequency of the mutation exceeded a threshold randomly drawn on a logarithmic scale within the interval [0.001, 0.1], we altered the fitness of this mutation to be beneficial with selection coefficient *s*, leading to relative fitness values of 1, 1 +*s*, and 1 +*s* for homozygous ancestral, heterozygous, and homozygous derived allele genotypes at this selected locus, respectively. However, for adaptive introgression, in addition to the CEU population, we simulated a second unsampled population that represents a population similarly diverged with humans as Neanderthals [Kuhlwilm et al., 2016], having a split time from the human representative (CEU) drawn uniformly at random from 27,000 to 13,000 generations prior to sampling. Under adaptive introgression, the beneficial allele is introduced in the unsampled Neanderthal representative, and then haplotypes are introgressed into the sampled population from the Neanderthal representative 20 generations afterward at admixture rate that is between 1% and 8% and is drawn uniformly at random. For balancing selection, a single selected mutation was introduced earlier compared to other selection simulations, such that the time of introduction is drawn uniformly at random from 80,000 to 40,000 generations prior to sampling with dominance parameter *h* = (*x* − 1)*/*(2*x* − 1), where equilibrium minor allele frequency *x* ∈ [0.2, 0.5) was drawn uniformly at random. This parameterization resulted in relative fitness of 1, 1 + *hs*, and 1 + *s* for homozygous ancestral, heterozygous, and homozygous derived allele genotypes at this balanced locus, respectively.

### Input image generation

For the purpose of training CNN models that can uncover and learn the signals of evolutionary processes to make inferences about their action on genomes, we compute images summarizing three distinct statistics across a sample of *n* haplotypes (*n/*2 diploid individuals). The first statistic is the representation of haplotype frequencies in a window of size *s* SNPs, and computed across *m* windows. Let **I**^(1)^ ∈ ℝ*^n×m^* be a matrix that will be used to created the first image. Consider window *j*, which corresponds to column *j* of **I**^(1)^ and has *k* distinct haplotypes observed with counts *h*_1_*, h*_2_*, . . . , h_k_* ∈ {1, 2*, . . . , n*} and are ordered based on their counts *h*_1_ ≥ *h*_2_ ≥ *. . .* ≥ *h_k_*. Let the column vector of ones of dimension *h_k_* be denoted by **1***_hk_* ∈ ℝ^*h_k_*^, and thus *k***1***_h_* is a column vector of dimension *h_k_* with all values equal to *k*. Using this notation, we then represent the *j*th column of **I**^(1)^ as

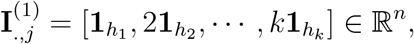

which can be used to fill the entire matrix **I**^(1)^ for columns *j* ∈ {1, 2*, . . . , m*}. This computation generates an image where for each column, the number one at the top is repeated for *h*_1_ rows, followed by the number two for *h*_2_ rows, and continuing until the last *h_k_* rows, which each have the number *k*.

The second statistic is the Hellinger distance [Hellinger, 1909] computed between the haplotype frequency distributions of two windows. Consider a pair of distinct windows *j*_1_*, j*_2_ ∈ {1, 2*, . . . , m*} for *j*_1_ ≠ *j*_2_ with *k*_1_ and *k*_2_ distinct haplotypes observed at windows *j*_1_ and *j*_2_, respectively. Denote the frequencies of haplotypes at window *j*_1_ by 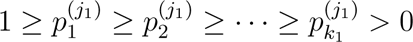 and the frequencies of haplotypes at window *j*_2_ by 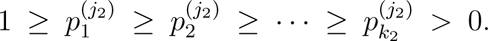 Assuming the maximum number of haplotypes observed between the two windows is *k* = max(*k*_1_*, k*_2_), define the following two frequency vectors

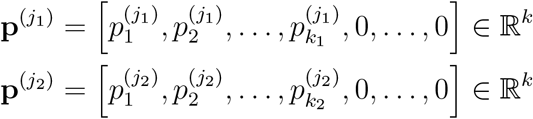

that have been appended with zeros to account for the unequal number of haplotypes between the windows. Given these two vectors, we compute the Hellinger distance between the two discrete haplotype distributions as [Le Cam, 2012]

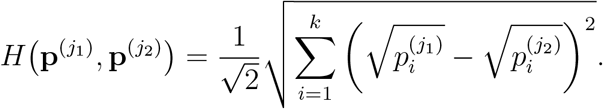

Let **I**^(2)^ ∈ ℝ*^m×m^* be a matrix that will be used to create the second image. The value in row *i* and column *j* of this matrix is computed as

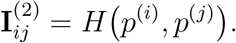

The third statistic is the Hamming distance [Hamming, 1986] (number of differing SNPs) between the *i*th most frequent haplotype and the (*i* + 1)th most frequent haplotype within each window, where for simplicity we have truncated the number of unique haplotypes to consider at *k*_max_ = 10. Consider a window *j* ∈ {1, 2*, . . . , m*} in the analyzed genomic region, which contains *s* SNPs. Further, suppose that this window has *k* distinct haplotypes ordered by counts *h*_1_ ≥ *h*_2_ ≥ · · · ≥ *h_k_ >* 0. Let the Hamming distance between haplotypes *i*_1_ and *i*_2_ be denoted by *D*(*i*_1_*, i*_2_) ∈ {1, 2*, . . . , s*} and let **I** ∈ R be a matrix that will be used to create the third image. Defining *K* = min(*k, k*_max_), we then represent the *j*th column of **I**^(3)^ as

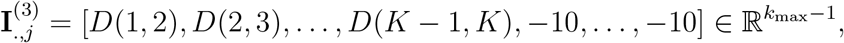

which has been appended with large magnitude negative values (-10) if there were not enough distinct haplotypes to fully compute the Hamming distances.

From our set of 4 × 6, 000 = 24, 000 replicates, the smallest number of SNPs in a given replicate flanking either side of the center SNP of the sequence was found to be 1,557, where the center SNP has position closest to one Mb. We therefore chose 3,115 SNPs (center SNP and closest 1,557 SNPs from each side of the center SNP) as the number of SNPs to construct the matrices **I**^(1)^, **I**^(2)^, and **I**^(3)^ that are used to create images for training downstream CNN models.

To capture fine scale patterns of variation, we considered a window size of *s* = 11 SNPs with a stride of 10 SNPs, which resulted in *m* = 311 windows. We also evaluated more moderate scale patterns of variation by employing a window of *s* = 21 SNPs with a stride of 10 SNPs, yielding *m* = 309 windows. Finally, we also interrogated coarser scale patterns of variation by using a window of *s* = 51 SNPs with a stride of 10 SNPs, which requires *m* = 307 windows. Each of these different window sizes was used to create a set of images derived from the matrices **I**^(1)^, **I**^(2)^, and **I**^(3)^ for CNN training and inference.

For the purpose of inference on empirical data from trained models, we computed the same three statistics for each of the 22 human autosomes from the 1000 Genomes Project dataset, retaining only biallelic SNPs with minor allele count of at least three that did not fall within regions of poor alignability and mappability according to a mean CRG (Centre for Genomic Regulation) score of 0.9 or higher within a non-overlapping 100 kilobase genomic segments [Talkowski et al., 2011, Derrien et al., 2012, Mughal et al., 2020]. For each autosome, we computed the three input matrices across 3,115 contiguous SNPs. We then create these three input matrices again at subsequent regions of 3,115 contiguous SNPs by sliding across the chromosome by a stride of 21 SNPs. The position of each focal region of 3,115 SNPs was taken as the chromosomal position of the central SNP in that region.

### Model architecture

The *M*_1_ model architecture is a simple CNNs inspired by the AlexNet architecture [Krizhevsky et al., 2012], which utilizes convolutional and max pooling layers for feature extraction and fully connected layers for class prediction. In contrast, *M*_2_ is a deeper convolutional model, based on the Inception-ResNet-v2 architecture [Szegedy et al., 2017]. The *SHADES* [*M*] model, for *M* ∈ {*M*_1_*, M*_2_}, is a network with the outputs of three CNNs stacked as input to a multilayer perceptron (MLP) model. These three CNNs are three *M*_1_ or *M*_2_ models trained on the three images, *i.e.*, **I**^(1)^, **I**^(2)^, and **I**^(3)^, that are constructed from windows of size *s* = 11, 21, and 51 SNPs. To preprocess the images, we scale the images and apply a channel-wise standardization operation. Specifically, we scale each image *i.e.*, **I**^(1)^, **I**^(2)^, and **I**^(3)^ for a genome replicate to a size of 32 × 32, which is the input dimension of the models. We also calculate the per-channel mean and standard deviation across the entire training dataset, and then standardize the images by subtracting the per-channel mean from each pixel value in the corresponding image channel and dividing by the per-channel standard deviation [Huang et al., 2017]. This approach ensures that the input data are centered and scaled, which aids in stabilizing and accelerating the training process of the neural network [Dai and Heckel, 2019].

For the *M*_1_ model (Figure S1), the inputs are single-channel images of size 32 × 32 pixels. This small input results in model architectures that are relatively shallow, with only a few convolutional and max pooling operations required to reduce the feature space to a small number. The model takes three single-channel images as input to its three parallel branches, with these three images rescaled depictions of **I**^(1)^, **I**^(2)^, and **I**^(3)^. Each branch takes a tensor of size *N* × 32 × 32 as input, where *N* is the sample size. Each branch follows the same sequential structure of a convolution layer, a rectified linear unit (ReLU) activation layer, and finally a max pooling layer, and this structure is repeated. The convolution layer enables the network to extract and recognize spatial features from input images by applying filters (kernels) that slide over the input [O’shea and Nash, 2015]. The ReLU activation layer is used for adding nonlinearity [Nair and Hinton, 2010]. The max pooling layer is used to downsample the spatial dimensions (height and width) of feature maps while retaining the most important information [O’shea and Nash, 2015]. We will consider this sequence of convolution, ReLU activation, and max-pooling as a convolution block and in each convolution block we will increase the number of channels for extracting a set of features from the input. For convolution, we employed a kernel size of 5 × 5 and for max pooling we used a kernel size of 2 × 2. We used two convolution blocks to reduce the size from 32 × 32 to 5 × 5, and increased the number of channels to 16, where each channel is a feature map corresponding to a particular kernel.

For the first convolution block, each input image of size 32 × 32 goes through a convolution layer with six kernels of size 5 × 5 that are applied with a stride of two with zero padding along the image. The output size after this convolution operation is 15 × 15. Then, in sequence, we use ReLU activation and a max pooling layer with a 2 × 2 kernel. Max pooling will reduce the size from 15 × 15 to 7 × 7, such that after the first convolution block, the output is a four-dimensional array of size *N* × 6 × 7 × 7. The second convolution block contains a convolution layer with 16 kernels of size 5 × 5, and is applied with a stride of one with zero padding. Other parameters are identical to those of the first convolution block. The output is of size 5 × 5 after the convolution layer in the second convolution block, and is finally reduced to a size of 2 × 2 after the max pooling layer. After the second convolution block, the output is a four-dimensional array of size *N* × 16 × 2 × 2, which is then flattened and concatenated with the flattened output from the other two images that have gone through the same feature extraction procedure. The flattened vector is of length 16 × 2 × 2 = 64 for each of the three parallel branches, which results in a concatenated vector of length 64 × 3 = 192. We then use this concatenated vector as input to a fully connected layer with ReLU activation for nonlinear modeling, followed by a second fully connected layer with ReLU activation, and finally an output layer for prediction. The second fully connected layer takes as input a vector of length 40, and the output layer contains is of four dimensions, with each dimension corresponding to the predicted probability of each of the four classes. These predicted probabilities are computed using a softmax activation [Bridle, 1990]. This neural network architecture results in the *M*_1_ model having 15,600 trainable parameters.

To extend beyond the relatively shallow architecture of *M*_1_, we introduced a deeper convolutional model (*M*_2_; Figure S1) based on the Inception-ResNet-v2 architecture [Szegedy et al., 2017]. It requires an input size of 299 × 299 and has 217,779,420 trainable parameters. Because *M*_2_ contains substantially more parameters than our custom models, we employed a transfer learning strategy using weights pretrained on the large-scale ImageNet natural image dataset [Deng et al., 2009]. We fine-tuned *M*_2_ separately for each genomic window size (11, 21, and 51 SNPs), allowing the pretrained filters to adapt to the structure of the input images in different scales.

The other two models (*SHADES* [*M*]; Figure 1) uses a stacking approach [Hastie et al., 2009] with the outputs of the last fully connected layers of three *M*_1_ or *M*_2_ models concatenated as input, where *M* ∈ {*M*_1_*, M*_2_}. Here, *SHADES* [*M*_1_] and *SHADES* [*M*_2_] represent the stacking of three *M*_1_ or three *M*_2_ models, respectively. The three *M*_1_ or *M*_2_ networks are trained beforehand, with their penultimate layer outputs extracted, rather than jointly training all the parameters of the *SHADES* [*M*] model. We consider training three *M*_1_ or *M*_2_ models, each corresponding to input images constructed from windows of size 11, 21, or 51 SNPs. By choosing different window sizes as input to these models, we can evaluate haplotype differences and frequency distributions at different scales, permitting better discrimination among selective events that leave genomic footprints of different sizes. By concatenating the vectors obtained from the penultimate layers of the three *M*_1_ or *M*_2_ models, we create the input to the first fully connected layer of the *SHADES* [*M*] model, which contains a ReLU activation, followed by a second fully connected layer with ReLU activation, and finally an output layer for classification. The penultimate layer of each *M*_1_ model is a vector of length 40, which yields a concatenated input vector for the first fully connected layer of the *SHADES* [*M*_1_] model of length 120. The second fully connected layer of *SHADES* [*M*_1_] takes as input a vector of length 120, which is followed by the four-dimensional output layer, with each dimension the probability of one of the four classes computed using a softmax activation. Given that the weights of the input *M*_1_ models are not re-estimated, the *SHADES* [*M*_1_] model is lightweight with only 15,004 trainable parameters. Following this approach, we constructed a stacked multi-window model from three *M*_2_ models, denoted *SHADES* [*M*_2_], which is analogous to *SHADES* [*M*_1_]. In *SHADES* [*M*_2_], the fine-tuned *M*_2_ networks for each window were frozen and used as feature extractors, and their outputs were concatenated and passed through nonlinear fully connected layers trained jointly for classification. Though *SHADES* [*M*_2_] has 653,382,064 total parameters, it has only 43,804 trainable parameters. This design employs the representational strength of a state-of-the-art vision model while preserving the complementary information captured across window sizes, resulting in substantial performance gains across all classes.

### Training

Training is accomplished using simulated data for all models. First, we train each model with 4,000 replicates per class, holding out 1,000 replicates per class as test data and 1,000 replicates per class as validation data to choose the optimal epoch for training the final model through early stopping [Prechelt, 2002]. Specifically, optimal epoch is selected as the epoch in which lowest validation loss is attained, where the loss is computed as the categorical cross entropy [Goodfellow et al., 2016b]. We then retrained the model using 5,000 observations per class for this optimal number of epochs. We estimated model parameters for each neural network using an Adam optimizer [Diederik and Jimmy, 2017], with learning rate of 0.001, exponential decay rates for the moment estimates *β*_1_ of 0.9 and *β*_2_ of 0.999 as optimizer parameters [Diederik and Jimmy, 2017]. For training the *SHADES* [*M*_1_] or *SHADES* [*M*_2_] models, the fitting process is identical to *M*_1_ or *M*_2_ models except for one difference in that the input to the neural network is not an image, but instead a stack of three *M*_1_ or *M*_2_ penultimate layer outputs of an input image. Therefore, to train the *SHADES* [*M*_1_] or *SHADES* [*M*_2_] models, it is required to train three *M*_1_ or *M*_2_ models beforehand and then start the fitting process of the *SHADES* [*M*_1_] or *SHADES* [*M*_2_] models.

### Computational Setup

We ran all computational operations on a high-performance computing cluster (HPC), using a single node with 20 CPU cores and 16 GB of memory. The parsing of the 24,000 simulated replicates from four classes using a window size of 21 SNPs to create the three input images takes around 25 hours, with roughly 22 hours to compute matrices **I**^(1)^ and **I**^(3)^, and only about three hours to compute **I**^(2)^ from the distributions displayed in **I**^(1)^. The parsing of empirical human autosomes for a window of size 21 SNPs took different amounts of time depending on chromosome size, with roughly 18 hours for the largest autosome (Chromosome 1). The training of the *M*_1_ model for 10 epochs takes roughly two hours and training of the *M*_2_ model for 10 epochs takes roughly three hours on our HPC with a graphics processing unit (GPU). Training *SHADES* [*M*_1_] and *SHADES* [*M*_2_] for 10 epochs required roughly four and five hours, respectively, on our HPC with a GPU. The inference (classification step) on different autosomes took different times depending on chromosomal size, with classification of all genomic windows on Chromosome 1 taking approximately 15 minutes for *SHADES* [*M*_1_] and 30 minutes for *SHADES* [*M*_2_] using a GPU.

## Acknowledgments

This work was supported by National Institutes of Health grant 2R35GM128590 and by National Science Foundation grant DEB-2302258. Computations for this research were performed using the services provided by Research Computing at the Florida Atlantic University.

## References

GW Abbott. Kcne4 and kcne5: K+ channel regulation and cardiac arrhythmogenesis. Gene, 593 (2):249–260, 2016.

ZM Ahmed, S Riazuddin, SL Bernstein, Z Ahmed, S Khan, AJ Griffith, RJ Morell, TB Friedman, S Riazuddin, and ER Wilcox. Mutations of the protocadherin gene pcdh15 cause usher syndrome type 1f. Am. J. Hum. Genet., 69(1):25–34, 2001.

M Akiyama. The roles of abca12 in epidermal lipid barrier formation and keratinocyte differentiation. Biochim. Biophys. Acta -Mol. Cell Biol. Lipids, 1841(3):435–440, 2014.

KN Alagramam, H Yuan, MH Kuehn, CL Murcia, S Wayne, CRS Srisailpathy, RB Lowry, R Knaus, L Van Laer, FP Bernier, et al. Mutations in the novel protocadherin pcdh15 cause usher syndrome type 1f. Hum. Mol. Genet., 10(16):1709–1718, 2001.

Y Allanore, M Saad, P Dieudé, J Avouac, JHW Distler, P Amouyel, M Matucci-Cerinic, G Riemekasten, P Airo, I Melchers, et al. Genome-wide scan identifies tnip1, psors1c1, and rhob as novel risk loci for systemic sclerosis. PLoS Genet., 7(7):e1002091, 2011.

MR Amin, M Hasan, SP Arnab, and M DeGiorgio. Tensor decomposition-based feature extraction and classification to detect natural selection from genomic data. Mol. Biol. Evol., 40(10):msad216, 2023.

MR Amin, M Hasan, and M DeGiorgio. Digital image processing to detect adaptive evolution. Mol. Biol. Evol., 41(12):msae242, 2024.

AM Andŕes, MJ Hubisz, A Indap, DG Torgerson, JD Degenhardt, AR Boyko, RN Gutenkunst, TJ White, ED Green, CD Bustamante, et al. Targets of balancing selection in the human genome. Mol. Biol. Evol., 26(12):2755–2764, 2009.

T Angata. Possible influences of endogenous and exogenous ligands on the evolution of human siglecs. Front. Immunol., 9:2885, 2018.

A Anguita-Ruiz, CM Aguilera, and Á Gil. Genetics of lactose intolerance: an updated review and online interactive world maps of phenotype and genotype frequencies. Nutrients, 12(9):2689, 2020.

SP Arnab, MR Amin, and M DeGiorgio. Uncovering footprints of natural selection through spectral analysis of genomic summary statistics. Mol. Biol. Evol., 40(7):msad157, 2023.

SP Arnab, AL Campelo dos Santos, M Fumagalli, and M DeGiorgio. Efficient detection and characterization of targets of natural selection using transfer learning. Mol. Biol. Evol., 42(5): msaf094, 2025.

SP Arnab, M Khan, ALC Dos Santos, M Fumagalli, and M DeGiorgio. Identifying adaptive footprints in the presence of demographic uncertainty. Genome biol. evol., 18(4):evag075, 2026.

Luisa Azevedo, Catarina Serrano, Antonio Amorim, and David N Cooper. Trans-species polymorphism in humans and the great apes is generally maintained by balancing selection that modulates the host immune response. Human genomics, 9(1):21, 2015.

A Barton, S Eyre, X Ke, A Hinks, J Bowes, E Flynn, P Martin, YEAR Consortium, BIRAC Consortium, AG Wilson, et al. Identification of af4/fmr2 family, member 3 (aff3) as a novel rheumatoid arthritis susceptibility locus and confirmation of two further pan-autoimmune susceptibility genes. Hum. Mol. Genet., 18(13):2518–2522, 2009.

NH Barton. Mutation and the evolution of recombination. Philos. Trans. R. Soc. B: Biol. Sci., 365 (1544):1281, 2010.

T Bersaglieri, PC Sabeti, N Patterson, T Vanderploeg, SF Schaffner, JA Drake, M Rhodes, DE Reich, and JN Hirschhorn. Genetic signatures of strong recent positive selection at the lactase gene. Am. J. Hum. Genet., 74(6):1111–1120, 2004.

D Biswas, MMM Rahman, Z Zong, and J Těsíc. Improving the energy efficiency of real-time dnn object detection via compression, transfer learning, and scale prediction. In 2022 IEEE International Conference on Networking, Architecture and Storage (NAS), pages 1–8. IEEE, 2022.

BD Bitarello, C De Filippo, JC Teixeira, JM Schmidt, P Kleinert, D Meyer, and AM Andŕes. Signatures of long-term balancing selection in human genomes. Genome Biol. Evol., 10(3):939–955, 2018.

W Boll, P Wagner, and N Mantei. Structure of the chromosomal gene and cdnas coding for lactase-phlorizin hydrolase in humans with adult-type hypolactasia or persistence of lactase. Am. J. Hum. Genet., 48(5):889, 1991.

V Bolón-Canedo, L Morán-Fernández, B Cancela, and A Alonso-Betanzos. A review of green artificial intelligence: Towards a more sustainable future. Neurocomputing, 599:128096, 2024.

TR Booker, S Yeaman, and MC Whitlock. Variation in recombination rate affects detection of outliers in genome scans under neutrality. Molecular ecology, 29(22):4274–4279, 2020.

I Borg and PJF Groenen. Modern multidimensional scaling: Theory and applications. Springer, 2005.

B Boutwell, D Hinds, M Agee, B Alipanahi, A Auton, RK Bell, K Bryc, SL Elson, P Fontanillas, NA Furlotte, et al. Replication and characterization of cadm2 and msra genes on human behavior. Heliyon, 3(7), 2017.

L Bouza, A Bugeau, and L Lannelongue. How to estimate carbon footprint when training deep learning models? a guide and review. ERC, 5(11):115014, 2023.

AR Boyko, SH Williamson, AR Indap, JD Degenhardt, RD Hernandez, KE Lohmueller, MD Adams, S Schmidt, JJ Sninsky, SR Sunyaev, et al. Assessing the evolutionary impact of amino acid mutations in the human genome. PLoS Genet., 4(5):e1000083, 2008.

JS Bridle. Probabilistic interpretation of feedforward classification network outputs, with relationships to statistical pattern recognition. In Neurocomputing: Algorithms, architectures and applications, pages 227–236. Springer, 1990.

JF Brinkworth and LB Barreiro. The contribution of natural selection to present-day susceptibility to chronic inflammatory and autoimmune disease. Curr. Opin. Immunol., 31:66–78, 2014.

L Bromham. The genome as a life-history character: why rate of molecular evolution varies between mammal species. Philosophical Transactions of the Royal Society B: Biological Sciences, 366 (1577):2503–2513, 2011.

L Bromham, X Hua, R Lanfear, and PF Cowman. Exploring the relationships between mutation rates, life history, genome size, environment, and species richness in flowering plants. The American Naturalist, 185(4):507–524, 2015.

SR Browning and BL Browning. Haplotype phasing: existing methods and new developments. Nat. Rev. Genet., 12(10):703–714, 2011.

G Bu, J Cam, and C Zerbinatti. Lrp in amyloid-*β* production and metabolism. Ann. N. Y. Acad. Sci., 1086(1):35–53, 2006.

R Cagliani, S Riva, U Pozzoli, M Fumagalli, GP Comi, N Bresolin, M Clerici, and M Sironi. Balancing selection is common in the extended mhc region but most alleles with opposite risk profile for autoimmune diseases are neutrally evolving. BMC Evol. Biol., 11(1):171, 2011.

SE Castel, P Mohammadi, WK Chung, Y Shen, and T Lappalainen. Rare variant phasing and haplotypic expression from rna sequencing with phaser. Nat. Commun., 7(1):12817, 2016.

D Castellano, A Eyre-Walker, and K Munch. Impact of mutation rate and selection at linked sites on dna variation across the genomes of humans and other homininae. Genome biology and evolution, 12(1):3550–3561, 2020.

RM Cecil and LA Sugden. On convolutional neural networks for selection inference: Revealing the effect of preprocessing on model learning and the capacity to discover novel patterns. PLoS Comput. Biol., 19(11):e1010979, 2023.

B Charlesworth. The effects of deleterious mutations on evolution at linked sites. Genetics, 190(1): 5–22, 2012.

B Charlesworth, MT Morgan, and D Charlesworth. The effect of deleterious mutations on neutral molecular variation. Genetics, 134(4):1289–1303, 1993.

D Charlesworth. Balancing selection and its effects on sequences in nearby genome regions. PLoS genet., 2(4):e64, 2006.

D Charlesworth, B Charlesworth, and MT Morgan. The pattern of neutral molecular variation under the background selection model. Genetics, 141(4):1619–1632, 1995.

Y Chen, P Liu, S Yi, C Fan, W Zhao, and J Liu. Investigating the shared genetic architecture between attention-deficit/hyperactivity disorder and risk taking behavior: A large-scale genomewide cross-trait analysis. J. Affect. Disord., 356:22–31, 2024.

X Cheng and M DeGiorgio. Detection of shared balancing selection in the absence of trans-species polymorphism. Mol. Biol. Evol., 36(1):177–199, 2019.

X Cheng and M DeGiorgio. Flexible mixture model approaches that accommodate footprint size variability for robust detection of balancing selection. Mol. Biol. Evol., 2020.

X Cheng, C Xu, and M DeGiorgio. Fast and robust detection of ancestral selective sweeps. Mol Ecol, 26(24):6871–6891, 2017.

V Colonna, Q Ayub, Y Chen, L Pagani, P Luisi, M Pybus, E Garrison, Y Xue, C Tyler-Smith, and 1000 Genomes Project Consortium. Human genomic regions with exceptionally high levels of population differentiation identified from 911 whole-genome sequences. Genome Biol., 15(6): R88, 2014.

JM Comeron. Background selection as baseline for nucleotide variation across the drosophila genome. PLoS Genet., 10(6):e1004434, 2014.

AD Cutter and BA Payseur. Genomic signatures of selection at linked sites: unifying the disparity among species. Nature Reviews Genetics, 14(4):262–274, 2013.

Z Dai and R Heckel. Channel normalization in convolutional neural network avoids vanishing gradients. arXiv preprint arXiv:1907.09539, 2019.

M Dannemann and J Kelso. The contribution of neanderthals to phenotypic variation in modern humans. Am. J. Hum. Genet., 101(4):578–589, 2017.

M Dannemann, AM Andrés, and J Kelso. Introgression of neandertal-and denisovan-like haplotypes contributes to adaptive variation in human toll-like receptors. Am. J. Hum. Genet., 98(1):22–33, 2016.

M DeGiorgio, KE Lohmueller, and R Nielsen. A model-based approach for identifying signatures of ancient balancing selection in genetic data. PLoS Genet., 10(8):e1004561, 2014.

J Deng, W Dong, R Socher, L Li, K Li, and L Fei-Fei. Imagenet: A large-scale hierarchical image database. In IEEE Conf. Comput. Vis. Pattern Recognit., pages 248–255. Ieee, 2009.

T Derrien, J Estellé, S Marco Sola, DG Knowles, E Raineri, R Guigó, and P Ribeca. Fast computation and applications of genome mappability. PloS one, 7(1):e30377, 2012.

PK Diederik and Ba Jimmy. Adam: A method for stochastic optimization, 2017. URL https://arxiv.org/abs/1412.6980.

L Duret. Neutral theory: the null hypothesis of molecular evolution. Nature education, 1(1):803–806, 2008.

E Eden, D Lipson, S Yogev, and Z Yakhini. Discovering motifs in ranked lists of dna sequences. PLoS Comput. Biol., 3(3):e39, 2007.

E Eden, R Navon, I Steinfeld, D Lipson, and Z Yakhini. Gorilla: a tool for discovery and visualization of enriched go terms in ranked gene lists. BMC Bioinform, 10(1):48, 2009.

H Eiberg, J Troelsen, M Nielsen, A Mikkelsen, J Mengel-From, KW Kjaer, and L Hansen. Blue eye color in humans may be caused by a perfectly associated founder mutation in a regulatory element located within the herc2 gene inhibiting oca2 expression. Hum. Genet., 123(2):177–187, 2008.

H Ellegren and N Galtier. Determinants of genetic diversity. Nat. Rev. Genet., 17(7):422–433, 2016.

D Enard and DA Petrov. Evidence that rna viruses drove adaptive introgression between neanderthals and modern humans. Cell, 175(2):360–371, 2018.

M Fanjul-Fernández, V Quesada, R Cabanillas, J Cadiñanos, T Fontanil, Á Obaya, AJ Ramsay, JL Llorente, A Astudillo, S Cal, et al. Cell–cell adhesion genes ctnna2 and ctnna3 are tumour suppressors frequently mutated in laryngeal carcinomas. Nat. Commun., 4(1):2531, 2013.

JC Fay and C Wu. Sequence divergence, functional constraint, and selection in protein evolution. Annu Rev Genomics Hum Genet., 4(1):213–235, 2003.

A Fijarczyk and W Babik. Detecting balancing selection in genomes: limits and prospects. Mol. Ecol., 24(14):3529–3545, 2015.

P Gerbault, C Moret, M Currat, and A Sanchez-Mazas. Impact of selection and demography on the diffusion of lactase persistence. PloS one, 4(7):e6369, 2009.

JH Gillespie. Population genetics: a concise guide. JHU press, 2004.

I Goodfellow, Y Bengio, and A Courville. Deep Learning. MIT Press, 2016a. http://www.deeplearningbook.org.

I Goodfellow, Y Bengio, A Courville, and Y Bengio. Deep learning, volume 1. MIT press Cambridge, 2016b.

A Gouy and L Excoffier. Polygenic patterns of adaptive introgression in modern humans are mainly shaped by response to pathogens. Mol. Biol. Evol., 37(5):1420–1433, 05 2020. ISSN 0737-4038. doi: 10.1093/molbev/msz306. URL 10.1093/molbev/msz306.

C Grey, F Baudat, and B De Massy. Genome-wide control of the distribution of meiotic recombination. PLoS biology, 7(2):e1000035, 2009.

AE Gururaj, RR Singh, SK Rayala, C Holm, P Den Hollander, H Zhang, S Balasenthil, AH Talukder, G Landberg, and R Kumar. Mta1, a transcriptional activator of breast cancer amplified sequence 3. Proc. Natl. Acad. Sci. U. S. A., 103(17):6670–6675, 2006.

Ryan N Gutenkunst, Ryan D Hernandez, Scott H Williamson, and Carlos D Bustamante. Inferring the joint demographic history of multiple populations from multidimensional snp frequency data. PLoS genetics, 5(10):e1000695, 2009.

BC Haller and PW Messer. SLiM 3: forward genetic simulations beyond the Wright–Fisher model. Mol. Biol. Evol., 36(3), 2019.

RW Hamming. Coding and information theory. Prentice-Hall, Inc, 1986.

H Harada, T Fujita, M Miyamoto, Y Kimura, M Maruyama, A Furia, T Miyata, and T Taniguchi. Structurally similar but functionally distinct factors, irf-1 and irf-2, bind to the same regulatory elements of ifn and ifn-inducible genes. Cell, 58(4):729–739, 1989.

A Harpak, A Bhaskar, and JK Pritchard. Mutation rate variation is a primary determinant of the distribution of allele frequencies in humans. PLoS genetics, 12(12):e1006489, 2016.

AM Harris, NR Garud, and M DeGiorgio. Detection and classification of hard and soft sweeps from unphased genotypes by multilocus genotype identity. Genetics, 210(4):1429–1452, 2018.

K Harris and JK Pritchard. Rapid evolution of the human mutation spectrum. elife, 6:e24284, 2017.

T Hastie, R Tibshirani, JH Friedman, and JH Friedman. The elements of statistical learning: data mining, inference, and prediction, volume 2. Springer, 2009.

E Hellinger. Neue begründung der theorie quadratischer formen von unendlichvielen veränderlichen. J Reine Angew Math, 136:210–271, 1909. URL http://eudml.org/doc/149313.

J Hermisson and PS Pennings. Soft sweeps: molecular population genetics of adaptation from standing genetic variation. Genetics, 169(4):2335–2352, 2005.

J Hey. What’s so hot about recombination hotspots? PLoS biology, 2(6):e190, 2004.

JN Honeyman, EP Simon, N Robine, R Chiaroni-Clarke, DG Darcy, IIP Lim, CE Gleason, JM Murphy, BR Rosenberg, L Teegan, et al. Detection of a recurrent dnajb1-prkaca chimeric transcript in fibrolamellar hepatocellular carcinoma. Science, 343(6174):1010–1014, 2014.

MR Hosseinzadeh Taher, F Haghighi, R Feng, MB Gotway, and J Liang. A systematic benchmarking analysis of transfer learning for medical image analysis. In MICCAI Workshop on Domain Adaptation and Representation Transfer, pages 3–13. Springer, 2021.

BJ Houston, J Nguyen, DJ Merriner, AE O’Connor, AM Lopes, L Nagirnaja, C Friedrich, S Kliesch, F Tüttelmann, KI Aston, et al. Axdnd1 is required to balance spermatogonial commitment and for sperm tail formation in mice and humans. Cell Death Dis., 15(7):499, 2024.

CY Huang and MN Rasband. Axon initial segments: structure, function, and disease. Ann. N. Y. Acad. Sci., 1420(1):46–61, 2018.

G Huang, Z Liu, L Van Der Maaten, and KQ Weinberger. Densely connected convolutional networks. In Proc IEEE Comput Soc Conf Comput Vis Pattern Recognit, pages 4700–4708, 2017.

X Huang, A Rymbekova, O Dolgova, O Lao, and M Kuhlwilm. Harnessing deep learning for population genetic inference. Nat. Rev. Genet., 25(1):61–78, 2024.

RR Hudson and NL Kaplan. The coalescent process in models with selection and recombination. Genetics, 120(3):831–840, 1988.

RR Hudson and NL Kaplan. Deleterious background selection with recombination. Genetics, 141 (4):1605, 1995.

RR Hudson, M Kreitman, and M Aguadé. A test of neutral molecular evolution based on nucleotide data. Genetics, 116(1):153–159, 1987.

E Huerta-Sánchez, X Jin, Asan, Z Bianba, BM Peter, N Vinckenbosch, Y Liang, X Yi, M He, M Somel, et al. Altitude adaptation in tibetans caused by introgression of denisovan-like dna. Nature, 512(7513):194–197, 2014.

Y Itan, BL Jones, CJE Ingram, DM Swallow, and MG Thomas. A worldwide correlation of lactase persistence phenotype and genotypes. BMC Evol. Biol., 10(1):36, 2010.

M Jain, GP Bhat, K Vijayraghavan, and MS Inamdar. Rudhira/bcas3 is a cytoskeletal protein that controls cdc42 activation and directional cell migration during angiogenesis. Exp. Cell Res., 318 (6):753–767, 2012.

B Janssens, S Goossens, K Staes, B Gilbert, J van Hengel, C Colpaert, E Bruyneel, M Mareel, and F Van Roy. *α*t-catenin: a novel tissue-specific *β*-catenin-binding protein mediating strong cell-cell adhesion. J. Cell Sci, 114(17):3177–3188, 2001.

JD Jensen, Y Kim, VB DuMont, CF Aquadro, and CD Bustamante. Distinguishing between selective sweeps and demography using dna polymorphism data. Genetics, 170(3):1401–1410, 2005.

D Ju and I Mathieson. The evolution of skin pigmentation-associated variation in west eurasia. Proc. Natl. Acad. Sci. U. S. A., 118(1):e2009227118, 2021.

EN Judd, AR Gilchrist, NR Meyerson, and SL Sawyer. Positive natural selection in primate genes of the type i interferon response. BMC Ecol. Evol., 21(1):65, 2021.

B Kang, S Cheng, J Peng, J Yan, and S Zhang. Interleukin-37 gene variants segregated anciently coexist during hominid evolution. Eur. J. Hum. Genet., 23(10):1392–1398, 2015.

S Kaul, JA Blackford Jr, J Chen, VV Ogryzko, and SS Simons Jr. Properties of the glucocorticoid modulatory element binding proteins gmeb-1 and-2: potential new modifiers of glucocorticoid receptor transactivation and members of the family of kdwk proteins. Mol. Endocrinol., 14(7): 1010–1027, 2000.

J Kere. Mapping and identifying genes for asthma and psoriasis. Phil. Trans. R. Soc. B, 360(1460): 1551, 2005.

AD Kern and DR Schrider. diplos/hic: an updated approach to classifying selective sweeps. G3, 8 (6):1959–1970, 2018.

MB Khawar, F Azam, N Sheikh, and K Abdul Mujeeb. How does interleukin-22 mediate liver regeneration and prevent injury and fibrosis? J. Immunol. Res., 2016(1):2148129, 2016.

M Kimura. The neutral theory of molecular evolution. Cambridge university press, 1985.

H Kitano, S Kageyama, SM Hewitt, R Hayashi, Y Doki, Y Ozaki, S Fujino, M Takikita, H Kubo, and J Fukuoka. Podoplanin expression in cancerous stroma induces lymphangiogenesis and predicts lymphatic spread and patient survival. Arch. Pathol. Lab. Med., 134(10):1520–1527, 2010.

K Kizhatil, W Yoon, PJ Mohler, LH Davis, JA Hoffman, and V Bennett. Ankyrin-g and *β*2-spectrin collaborate in biogenesis of lateral membrane of human bronchial epithelial cells. J. Biol. Chem., 282(3):2029–2037, 2007.

L Kong, C Sui, T Chen, L Zhang, W Zhao, Y Zheng, B Liu, X Cheng, and C Gao. The ubiquitin e3 ligase trim10 promotes sting aggregation and activation in the golgi apparatus. Cell Rep., 42 (4), 2023.

K Korfmann, OE Gaggiotti, and M Fumagalli. Deep learning in population genetics. Genome biol. evol., 15(2):evad008, 2023.

A Köttgen, C Pattaro, CA Böger, C Fuchsberger, M Olden, NL Glazer, A Parsa, X Gao, Q Yang, AV Smith, et al. New loci associated with kidney function and chronic kidney disease. Nat. Genet., 42(5):376–384, 2010.

A Krizhevsky, I Sutskever, and GE Hinton. Imagenet classification with deep convolutional neural networks. NIPS, 25, 2012.

JB Kruskal. Multidimensional scaling by optimizing goodness of fit to a nonmetric hypothesis. Psychometrika, 29(1):1–27, 1964.

M Kuhlwilm, I Gronau, MJ Hubisz, C De Filippo, J Prado-Martinez, M Kircher, Q Fu, HA Burbano, C Lalueza-Fox, M de La Rasilla, et al. Ancient gene flow from early modern humans into eastern neanderthals. Nature, 530(7591):429–433, 2016.

H Kumar, M Panigrahi, A Panwar, D Rajawat, SS Nayak, KA Saravanan, K Kaisa, S Parida, B Bhushan, and T Dutt. Machine-learning prospects for detecting selection signatures using population genomics data. J. Comput. Biol., 29(9):943–960, 2022.

S Kumar and S Subramanian. Mutation rates in mammalian genomes. Proceedings of the National Academy of Sciences, 99(2):803–808, 2002.

V Labrie, OJ Buske, E Oh, R Jeremian, C Ptak, G Gasiūnas, A Maleckas, R Petereit, A Z^̌^virbliene, K Adamonis, et al. Lactase nonpersistence is directed by dna-variation-dependent epigenetic aging. Nat. Struct. Mol. Biol., 23(6):566–573, 2016.

ME Laird, A Mohsen, D Duffy, R Mamdouh, L LeFouler, A Casrouge, M El-Daly, M Rafik, M Abdel-Hamid, A Soulier, et al. Apolipoprotein h expression is associated with il28b genotype and viral clearance in hepatitis c virus infection. J. Hepatol., 61(4):770–776, 2014.

ME Lauterbur, K Munch, and D Enard. Versatile detection of diverse selective sweeps with flex-sweep. Mol. Biol. Evol., 40(6):msad139, 2023.

TW Laver, RC Caswell, KA Moore, J Poschmann, MB Johnson, MM Owens, S Ellard, KH Paszkiewicz, and MN Weedon. Pitfalls of haplotype phasing from amplicon-based long-read sequencing. Sci. Rep., 6(1):21746, 2016.

L Le Cam. Asymptotic methods in statistical decision theory. Springer Science & Business Media, 2012.

JYW Lee and JA McGrath. Mutations in genes encoding desmosomal proteins: spectrum of cutaneous and extracutaneous abnormalities. Br. J. Dermatol., 184(4):596–605, 2021.

EM Leffler, Z Gao, S Pfeifer, L Ségurel, A Auton, O Venn, R Bowden, R Bontrop, JD Wall, G Sella, et al. Multiple instances of ancient balancing selection shared between humans and chimpanzees. Science, 339(6127):1578–1582, 2013.

C Lenormand, H Bausinger, F Gross, F Signorino-Gelo, S Koch, M Peressin, D Fricker, J Cazenave, T Bieber, D Hanau, et al. Hla-dqa2 and hla-dqb2 genes are specifically expressed in human langerhans cells and encode a new hla class ii molecule. J. Immunol., 188(8):3903–3911, 2012.

DE Levy and JE Darnell Jr. Stats: transcriptional control and biological impact. Nat. Rev. Mol. Cell Biol., 3(9):651–662, 2002.

C Liu, S Musco, NM Lisitsina, SY Yaklichkin, and NA Lisitsyn. Genomic organization of a new candidate tumor suppressor gene, lrp1b. Genomics, 69(2):271–274, 2000.

C Liu, Y Li, LM Obermoeller-McCormick, AL Schwartz, and G Bu. The putative tumor suppressor lrp1b, a novel member of the low density lipoprotein (ldl) receptor family, exhibits both overlapping and distinct properties with the ldl receptor-related protein. J. Biol. Chem., 276(31): 28889–28896, 2001.

X Liu, RT Ong, EN Pillai, AM Elzein, KS Small, TG Clark, DP Kwiatkowski, and Y Teo. Detecting and characterizing genomic signatures of positive selection in global populations. Am. J. Hum. Genet., 92(6):866–881, 2013.

L Loewe and WG Hill. The population genetics of mutations: good, bad and indifferent. Philos. Trans. R. Soc. B: Biol. Sci., 365(1544):1153, 2010.

Po-Ru Loh, P Danecek, PF Palamara, C Fuchsberger, Y A Reshef, H K Finucane, S Schoenherr, L Forer, S McCarthy, GR Abecasis, et al. Reference-based phasing using the haplotype reference consortium panel. Nat. Genet., 48(11):1443–1448, 2016.

DI Lou, Ross M McBee, UQ Le, AC Stone, GK Wilkerson, AM Demogines, and SL Sawyer. Rapid evolution of brca1 and brca2 in humans and other primates. BMC Evol. Biol., 14(1):155, 2014.

JR Mackey, SYM Yao, KM Smith, E Karpinski, SA Baldwin, CE Cass, and JD Young. Gemcitabine transport in xenopus oocytes expressing recombinant plasma membrane mammalian nucleoside transporters. J. Natl. Cancer Inst., 91(21):1876–1881, 1999.

ME Mathyer, EA Brettmann, AD Schmidt, ZA Goodwin, IY Oh, AM Quiggle, E Tycksen, N Ramakrishnan, SJ Matkovich, E Guttman-Yassky, et al. Selective sweep for an enhancer involucrin allele identifies skin barrier adaptation out of africa. Nat. Commun., 12(1):2557, 2021.

G McVicker, D Gordon, C Davis, and P Green. Widespread genomic signatures of natural selection in hominid evolution. PLoS genet., 5(5):e1000471, 2009.

MC Méchin, M Enji, R Nachat, S Chavanas, M Charveron, A Ishida-Yamamoto, G Serre, H Taka-hara, and M Simon. The peptidylarginine deiminases expressed in human epidermis differ in their substrate specificities and subcellular locations. Cell. Mol. Life Sci., 62(17):1984–1995, 2005.

F Mignone, C Gissi, S Liuni, and G Pesole. Untranslated regions of mrnas. Genome Biol., 3(3): reviews0004–1, 2002.

J Min, M Gupta, MM Desai, and DB Weissman. Spatial structure alters the site frequency spectrum produced by hitchhiking. Genetics, 222(3):iyac139, 2022.

J Min, Y Ning, NS Pope, F Baumdicker, and AD Kern. Neural posterior estimation for population genetics. Genetics, 233(3):iyag107, 2026.

JM Moon, DM Aronoff, JA Capra, P Abbot, and A Rokas. Examination of signatures of recent positive selection on genes involved in human sialic acid biology. G3: Genes Genomes Genet., 8 (4):1315–1325, 2018.

MR Mughal and M DeGiorgio. Localizing and classifying adaptive targets with trend filtered regression. Mol. Biol. Evol., 36(2):252–270, 2019.

MR Mughal, H Koch, J Huang, F Chiaromonte, and M DeGiorgio. Learning the properties of adaptive regions with functional data analysis. PLoS genet., 16(8):e1008896, 2020.

H Mühl and M Bachmann. Il-18/il-18bp and il-22/il-22bp: two interrelated couples with therapeutic potential. Cell. Signal., 63:109388, 2019.

S Myers, L Bottolo, C Freeman, G McVean, and P Donnelly. A fine-scale map of recombination rates and hotspots across the human genome. Science, 310(5746):321–324, 2005.

R Nachat, M Méchin, H Takahara, S Chavanas, M Charveron, G Serre, and M Simon. Peptidylarginine deiminase isoforms 1–3 are expressed in the epidermis and involved in the deimination of k1 and filaggrin. J. Invest. Dermatol., 124(2):384–393, 2005.

V Nair and GE Hinton. Rectified linear units improve restricted boltzmann machines. In ICML-10, pages 807–814, 2010.

LE Nicolaisen and MM Desai. Distortions in genealogies due to purifying selection and recombination. Genetics, 195(1):221–230, 2013.

R Nielsen, MJ Hubisz, I Hellmann, D Torgerson, AM Andrés, A Albrechtsen, R Gutenkunst, MD Adams, M Cargill, A Boyko, et al. Darwinian and demographic forces affecting human protein coding genes. Genome Res., 19(5):838–849, 2009.

M Nilsson, S Wasylik, M Mörgelin, AI Olin, JCM Meijers, RHWM Derksen, PG De Groot, and H Herwald. The antibacterial activity of peptides derived from human beta-2 glycoprotein i is inhibited by protein h and m1 protein from streptococcus pyogenes. Mol. Microbiol., 67(3): 482–492, 2008.

MF Nold, CA Nold-Petry, JA Zepp, BE Palmer, P Bufler, and CA Dinarello. Il-37 is a fundamental inhibitor of innate immunity. Nat. Immunol., 11(11):1014–1022, 2010.

CA Nold-Petry, CY Lo, I Rudloff, KD Elgass, S Li, MP Gantier, AS Lotz-Havla, SW Gersting, SX Cho, JC Lao, et al. Il-37 requires the receptors il-18r*α* and il-1r8 (sigirr) to carry out its multifaceted anti-inflammatory program upon innate signal transduction. Nat. Immunol., 16(4): 354–365, 2015.

J O’Connell, D Gurdasani, O Delaneau, N Pirastu, S Ulivi, M Cocca, M Traglia, J Huang, JE Huffman, I Rudan, et al. A general approach for haplotype phasing across the full spectrum of relatedness. PLoS Genet., 10(4):e1004234, 2014.

V Oji, K Eckl, K Aufenvenne, M Nätebus, T Tarinski, K Ackermann, N Seller, D Metze, G Nürnberg, R Fölster-Holst, et al. Loss of corneodesmosin leads to severe skin barrier defect, pruritus, and atopy: unraveling the peeling skin disease. Am. J. Hum. Genet., 87(2):274–281, 2010.

K O’shea and R Nash. An introduction to convolutional neural networks. arXiv:1511.08458, 2015.

SJ Pan and Q Yang. A survey on transfer learning. IEEE Trans. Knowl. Data Eng., 22(10): 1345–1359, 2009.

BA Payseur and MW Nachman. Microsatellite variation and recombination rate in the human genome. Genetics, 156(3):1285–1298, 2000.

JV Peñalba and JBW Wolf. From molecules to populations: appreciating and estimating recombination rate variation. Nature Reviews Genetics, 21(8):476–492, 2020.

PS Pennings and J Hermisson. Soft sweeps ii—molecular population genetics of adaptation from recurrent mutation or migration. Mol. Biol. Evol., 23(5):1076–1084, 2006.

TD Petes. Meiotic recombination hot spots and cold spots. Nature Reviews Genetics, 2(5):360–369, 2001.

L Prechelt. Early stopping-but when? In Neural Networks: Tricks of the trade, pages 55–69. Springer, 2002.

M Przeworski. The signature of positive selection at randomly chosen loci. Genetics, 160(3): 1179–1189, 2002.

C Pucharcos, X Estivill, and S de la Luna. Intersectin 2, a new multimodular protein involved in clathrin-mediated endocytosis. FEBS Lett., 478(1-2):43–51, 2000.

ZR Qian, T Sano, K Yoshimoto, SL Asa, S Yamada, N Mizusawa, and E Kudo. Tumor-specific downregulation and methylation of the cdh13 (h-cadherin) and cdh1 (e-cadherin) genes correlate with aggressiveness of human pituitary adenomas. Mod. Pathol., 20(12):1269–1277, 2007.

J Qiu, Q Zhou, W Ye, Q Chen, and Y Bao. Sweepcluster: A snp clustering tool for detecting gene-specific sweeps in prokaryotes. BMC Bioinform., 23(1):19, 2022.

F Racimo, S Sankararaman, R Nielsen, and E Huerta-Sánchez. Evidence for archaic adaptive introgression in humans. Nat. Rev. Genet., 16(6):359–371, 2015.

JS Rees, S Castellano, and AM Andrés. The genomics of human local adaptation. Trends in Genetics, 36(6):415–428, 2020.

PF Reilly, A Tjahjadi, SL Miller, JM Akey, and S Tucci. The contribution of neanderthal introgression to modern human traits. Curr. Biol., 32(18):R970–R983, 2022.

C Retel, H Märkle, L Becks, and PGD Feulner. Ecological and evolutionary processes shaping viral genetic diversity. Viruses, 11(3):220, 2019.

T Ruths and L Nakhleh. Boosting forward-time population genetic simulators through genotype compression. BMC Bioinform, 14:1–12, 2013.

PC Sabeti, SF Schaffner, B Fry, J Lohmueller, P Varilly, O Shamovsky, A Palma, TS Mikkelsen, D Altshuler, and ES Lander. Positive natural selection in the human lineage. science, 312(5780): 1614–1620, 2006.

PC Sabeti, P Varilly, B Fry, J Lohmueller, E Hostetter, C Cotsapas, X Xie, EH Byrne, SA McCar-roll, R Gaudet, et al. Genome-wide detection and characterization of positive selection in human populations. Nature, 449(7164):913–918, 2007.

MK Sakharkar, VTK Chow, and P Kangueane. Distributions of exons and introns in the human genome. In Silico Biol., 4(4):387–393, 2004.

AJ Sams, A Dumaine, Y Nédélec, V Yotova, C Alfieri, JE Tanner, PW Messer, and LB Barreiro. Adaptively introgressed neandertal haplotype at the oas locus functionally impacts innate immune responses in humans. Genome Biol., 17(1):246, 2016.

S Sanchez-Roige, MV Jennings, HHA Thorpe, JE Mallari, LC van der Werf, SB Bianchi, Y Huang, C Lee, TT Mallard, SA Barnes, et al. Cadm2 is implicated in impulsive personality and numerous other traits by genome-and phenome-wide association studies in humans and mice. Transl. Psychiatry, 13(1):167, 2023.

A Scally and R Durbin. Revising the human mutation rate: implications for understanding human evolution. Nat Rev Genet, 13(10):745–753, 2012.

DR Schrider. Background selection does not mimic the patterns of genetic diversity produced by selective sweeps. Genetics, 216(2):499–519, 2020.

DR Schrider and AD Kern. Soft sweeps are the dominant mode of adaptation in the human genome. Mol. Biol. Evol., 34(8):1863–1877, 2017.

R Schwarzenbacher, K Zeth, K Diederichs, A Gries, GM Kostner, P Laggner, and R Prassl. Crystal structure of human *β*2-glycoprotein i: implications for phospholipid binding and the antiphos-pholipid syndrome. EMBO J., 18(22):6228–6239, 1999.

NS Scrimshaw and EB Murray. The acceptability of milk and milk products in populations with a high prevalence of lactose intolerance. Am. J. Clin. Nutr., 48(4):1142–1159, 1988.

M Sebesta, CDO Cooper, A Ariza, CJ Carnie, and D Ahel. Structural insights into the function of zranb3 in replication stress response. Nat. Commun., 8(1):15847, 2017.

L Ségurel and C Bon. On the evolution of lactase persistence in humans. Annu. Rev. Genomics Hum. Genet., 18:297–319, 2017.

D Setter, S Mousset, X Cheng, R Nielsen, M DeGiorgio, and J Hermisson. Volcanofinder: genomic scans for adaptive introgression. PLoS Genet., 16(6):e1008867, 2020.

A Sharif Razavian, H Azizpour, J Sullivan, and S Carlsson. Cnn features off-the-shelf: an astounding baseline for recognition. In CVPRW, pages 806–813, 2014.

Z Shen, L Wei, Z Yu, Z Yao, J Cheng, Y Wang, X Song, and M Li. The roles of trims in antiviral innate immune signaling. Front. Cell. Infect. Microbiol., 11:628275, 2021.

Y Shirogane, R Takemoto, T Suzuki, T Kameda, K Nakashima, T Hashiguchi, and Y Yanagi. Cadm1 and cadm2 trigger neuropathogenic measles virus-mediated membrane fusion by acting in cis. J. Virol., 95(14):10–1128, 2021.

KM Siewert and BF Voight. Detecting long-term balancing selection using allele frequency correlation. Mol. Biol. Evol., 34(11):2996–3005, 2017.

MP Sikara, JG Routsias, M Samiotaki, G Panayotou, HM Moutsopoulos, and PG Vlachoyiannopoulos. *β*2 glycoprotein i (*β*2gpi) binds platelet factor 4 (pf4): implications for the pathogenesis of antiphospholipid syndrome. Blood, 115(3):713–723, 2010.

K Sikorski, A Czerwoniec, JM Bujnicki, J Wesoly, and HAR Bluyssen. Stat1 as a novel therapeutical target in pro-atherogenic signal integration of ifn*γ*, tlr4 and il-6 in vascular disease. Cytokine Growth Factor Rev., 22(4):211–219, 2011.

JM Smith and J Haigh. The hitch-hiking effect of a favourable gene. Genet. Res., 23(1):23–35, 1974.

CS Smukowski and MAF Noor. Recombination rate variation in closely related species. Heredity, 107(6):496–508, 2011.

B Star and HG Spencer. Effects of genetic drift and gene flow on the selective maintenance of genetic variation. Genetics, 194(1):235–244, 2013.

RA Sturm, DL Duffy, ZZ Zhao, FPN Leite, MS Stark, NK Hayward, NG Martin, and GW Montgomery. A single snp in an evolutionary conserved region within intron 86 of the herc2 gene determines human blue-brown eye color. Am. J. Hum. Genet., 82(2):424–431, 2008.

Z Su and X Tao. Current understanding of il-37 in human health and disease. Front. Immunol., 12:696605, 2021.

LA Sugden, EG Atkinson, AP Fischer, S Rong, BM Henn, and S Ramachandran. Localization of adaptive variants in human genomes using averaged one-dependence estimation. Nat. Commun., 9(1):703, 2018.

Q Sun and Y Li. Advances in haplotype phasing and genotype imputation. Nat. Rev. Genet., pages 1–15, 2025.

C Szegedy, S Ioffe, V Vanhoucke, and A Alemi. Inception-v4, inception-resnet and the impact of residual connections on learning. In Proc. AAAI Conf. Artif. Intell., volume 31, 2017.

F Tajima. Statistical method for testing the neutral mutation hypothesis by dna polymorphism. Genetics, 123(3):585–595, 1989.

R Takemoto, Y Hirai, S Watanabe, H Harada, T Suzuki, T Hashiguchi, Y Yanagi, and Y Shirogane. Interaction of the hemagglutinin stalk region with cell adhesion molecule (cadm) 1 and cadm2 mediates the spread between neurons and neuropathogenicity of measles virus with a hyperfusogenic fusion protein. J. Virol., 97(5):e00340–23, 2023.

ME Talkowski, C Ernst, A Heilbut, C Chiang, C Hanscom, A Lindgren, A Kirby, S Liu, B Muddukrishna, TK Ohsumi, et al. Next-generation sequencing strategies enable routine detection of balanced chromosome rearrangements for clinical diagnostics and genetic research. Am J Hum Genet, 88(4):469–481, 2011.

J Terhorst, JA Kamm, and YS Song. Robust and scalable inference of population history from hundreds of unphased whole genomes. Nat Genet, 49(2):303–309, 2017.

The 1000 Genomes Project Consortium. A global reference for human genetic variation. Nature, 526, 2015.

SA Tishkoff, FA Reed, A Ranciaro, BF Voight, CC Babbitt, JS Silverman, K Powell, HM Mortensen, JB Hirbo, M Osman, et al. Convergent adaptation of human lactase persistence in africa and europe. Nat. Genet., 39(1):31–40, 2007.

L Torada, L Lorenzon, A Beddis, U Isildak, L Pattini, S Mathieson, and M Fumagalli. Imagene: a convolutional neural network to quantify natural selection from genomic data. BMC bioinformatics, 20(Suppl 9):337, 2019.

S Tsukumo, PG Subramani, N Seija, M Tabata, Y Maekawa, Y Mori, C Ishifune, Y Itoh, M Ota, K Fujio, et al. Aff3, a susceptibility factor for autoimmune diseases, is a molecular facilitator of immunoglobulin class switch recombination. Sci. Adv., 8(34):eabq0008, 2022.

M Uhlén, L Fagerberg, BM Hallström, C Lindskog, P Oksvold, A Mardinoglu, Å Sivertsson, C Kampf, E Sjöstedt, A Asplund, et al. Tissue-based map of the human proteome. Science, 347(6220):1260419, 2015.

S van den Belt and N Alachiotis. Fast and accurate deep learning scans for signatures of natural selection in genomes using faster-nn. Commun Biol, 8(1):58, 2025.

S van den Belt, H Zhao, and N Alachiotis. Scalable cnn-based classification of selective sweeps using derived allele frequencies. Bioinformatics, 40(Supplement 2):ii29–ii36, 2024.

L Van der Maaten and G Hinton. Visualizing data using t-sne. J. Mach. Learn. Res., 9(11), 2008.

M Vancamelbeke, T Vanuytsel, R Farré, S Verstockt, M Ferrante, G Van Assche, P Rutgeerts, F Schuit, S Vermeire, I Arijs, et al. Genetic and transcriptomic bases of intestinal epithelial barrier dysfunction in inflammatory bowel disease. *nflamm*. Bowel Dis., 23(10):1718–1729, 2017.

KC Verbist and KD Klonowski. Functions of il-15 in anti-viral immunity: multiplicity and variety. Cytokine, 59(3):467–478, 2012.

JJ Vitti, SR Grossman, and PC Sabeti. Detecting natural selection in genomic data. Annu. Rev. Genet., 47(1):97–120, 2013.

W Wang, H Zhao, and G Han. Host-virus arms races drive elevated adaptive evolution in viral receptors. J. Virol., 94(16):10–1128, 2020.

R Weston, H Peeters, and D Ahel. Zranb3 is a structure-specific atp-dependent endonuclease involved in replication stress response. Genes Dev., 26(14):1558–1572, 2012.

H Xu, L Zong, DD Ray, L Lei, DR Schrider, F Baumdicker, and S Mathieson. Interpreting convolutional neural networks in population genetics. Mol. Biol. Evol., page msag256, 10 2026. ISSN 1537-1719. doi: 10.1093/molbev/msag256. URL 10.1093/molbev/msag256.

K Yuan, X Ni, C Liu, Y Pan, L Deng, R Zhang, Y Gao, X Ge, J Liu, X Ma, et al. Refining models of archaic admixture in eurasia with archaicseeker 2.0. Nat. Commun., 12(1):6232, 2021.

X Yuan, DJ Miller, J Zhang, D Herrington, and Y Wang. An overview of population genetic data simulation. J Comput Biol, 19(1):42–54, 2012.

F Zhang, L Christiansen, J Thomas, D Pokholok, R Jackson, N Morrell, Y Zhao, M Wiley, E Welch, E Jaeger, et al. Haplotype phasing of whole human genomes using bead-based barcode partitioning in a single tube. Nat. Biotechnol., 35(9):852–857, 2017.

H Zhang, AT Kho, Q Wu, AJ Halayko, K Limbert Rempel, RP Chase, NB Sweezey, ST Weiss, and F Kaplan. Crispld2 (lgl1) inhibits proinflammatory mediators in human fetal, adult, and copd lung fibroblasts and epithelial cells. Physiol. Rep., 4(17):e12942, 2016.

X Zhang, J Zhang, T Wang, MA Esteban, and D Pei. Esrrb activates oct4 transcription and sustains self-renewal and pluripotency in embryonic stem cells. J. Biol. Chem., 283(51):35825–35833, 2008.

C Zhao, M Jia, H Song, Z Yu, W Wang, Q Li, L Zhang, W Zhao, and X Cao. The e3 ubiquitin ligase trim40 attenuates antiviral immune responses by targeting mda5 and rig-i. Cell Rep., 21 (6):1613–1623, 2017.

Y Zheng and T Wiehe. Adaptation in structured populations and fuzzy boundaries between hard and soft sweeps. PLoS Comput. Biol., 15(11):e1007426, 2019.

L Zuo, K Wang, X Zhang, CR Li, F Zhang, X Wang, W Chen, G Gao, H Zhang, JH Krystal, et al. Rare serinc2 variants are specific for alcohol dependence in subjects of european descent. Pharmacogenetics Genom., 23(8):395, 2013.

