## Supplementary text for "Multi-scale haplotype signatures distinguish among evolutionary processes"

### Supplementary material

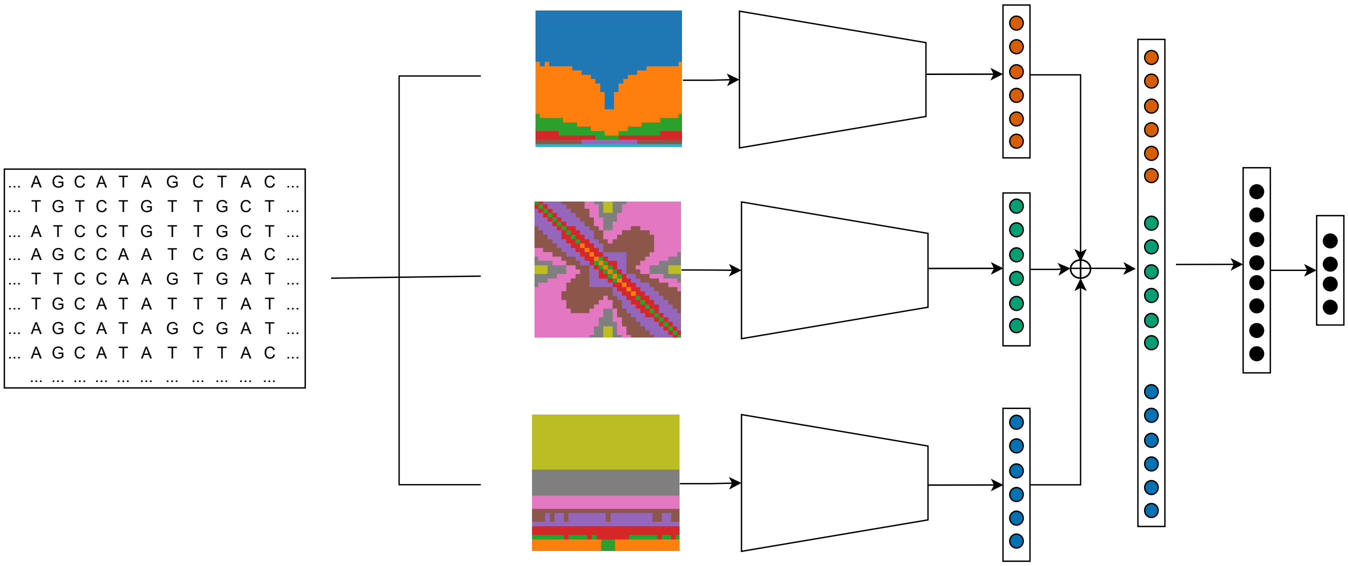


Figure S1: High-level schematic of the $M_1$ and $M_2$ models. The overall architecture of $M_2$ is similar to that of $M_1$, except that each parallel branch uses Inception-ResNet-v2 rather than a shallow CNN. Genomic sequence alignments are processed to generate images of three statistics $\mathbf{I}^{(1)}$ (top), $\mathbf{I}^{(2)}$ (middle), and $\mathbf{I}^{(3)}$ (bottom) that are each fed to separate branches of a convolutional neural network (CNN). For $M_1$, each branch receives a single channel input image with dimension $32 \times 32$ and has two convolution blocks (a sequence of convolution, ReLU activation, and max pooling), followed by a flattened layer (orange, green, and blue vectors). This step produces flattened vectors of lengths 64 for each image in the $M_1$ model. For $M_2$, each single channel image is replicated across three identical channels each producing input images of dimension $299 \times 299$, as required by the Inception-ResNet-v2. Each $M_2$ branch uses an Inception-ResNet-v2 model initialized with ImageNet pretrained weights, with the final classification layer removed to use it as a feature extractor. Each branch produces a flattened feature vector of length 1,536 (orange, green, and blue vectors). For all three models, the feature vectors from the three parallel branches are concatenated ($\oplus$) into a single vector and used as input to a fully connected layer with ReLU activation, followed by another fully connected layer with ReLU activation, and finally a four-dimensional output layer with softmax activation. This output layer predicts the probability of neutrality, sweep, adaptive introgression, and balancing selection.

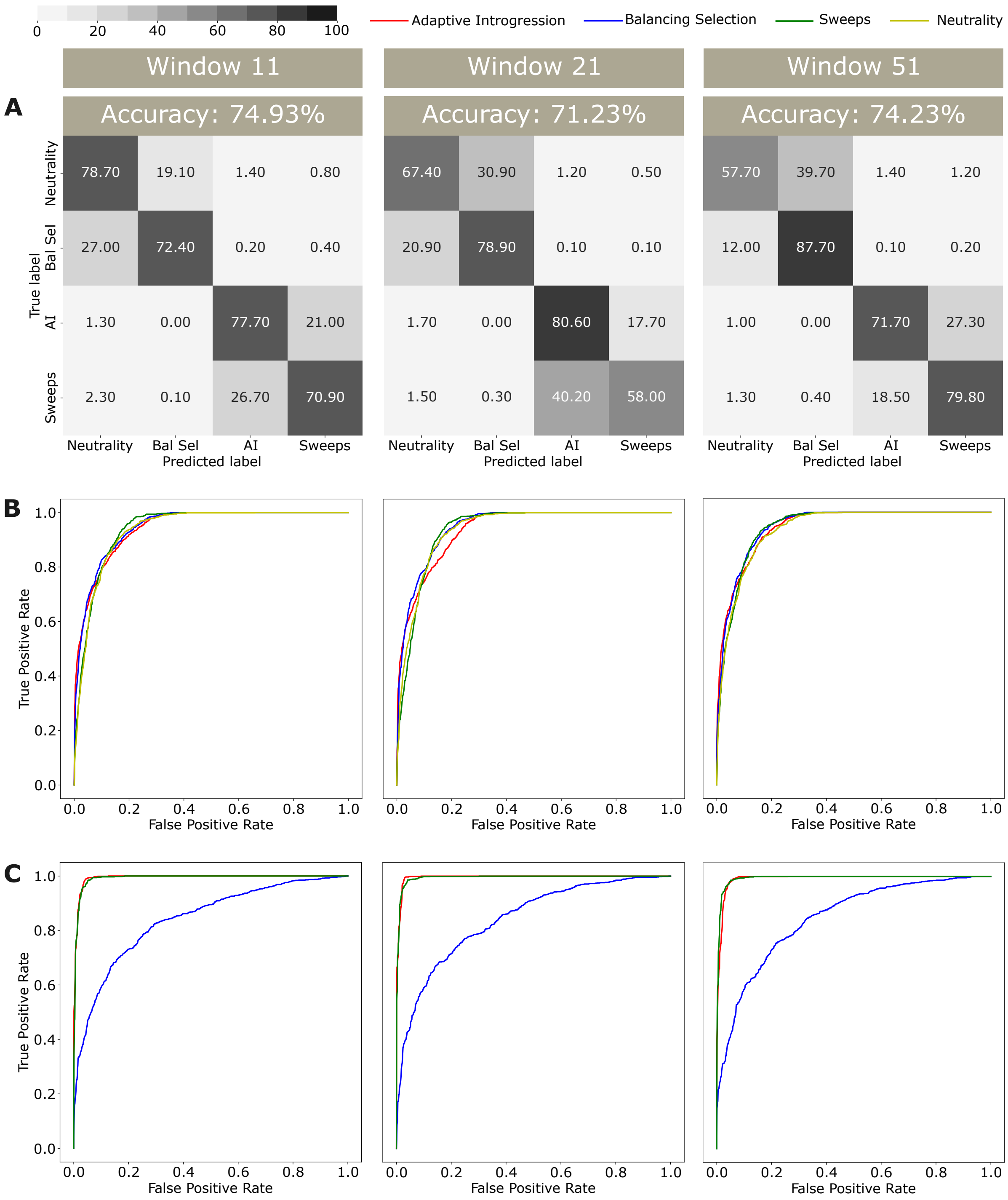


Figure S2: Classification rates and accuracies as depicted by confusion matrices (A) and power to distinguish evolutionary classes as depicted by receiver operating characteristic curves (B, C) for the $M_1$ model with windows of sizes 11 (left column), 21 (middle column), or 51 SNPs (right column). In the confusion matrices, AI denotes adaptive introgression and Bal Sel denotes balancing selection. (B) True positive rate for each class as a function of the false positive rate for all other classes. (C) True positive rate for balancing selection, adaptive introgression, and sweeps as a function of the false positive rate from neutrality.

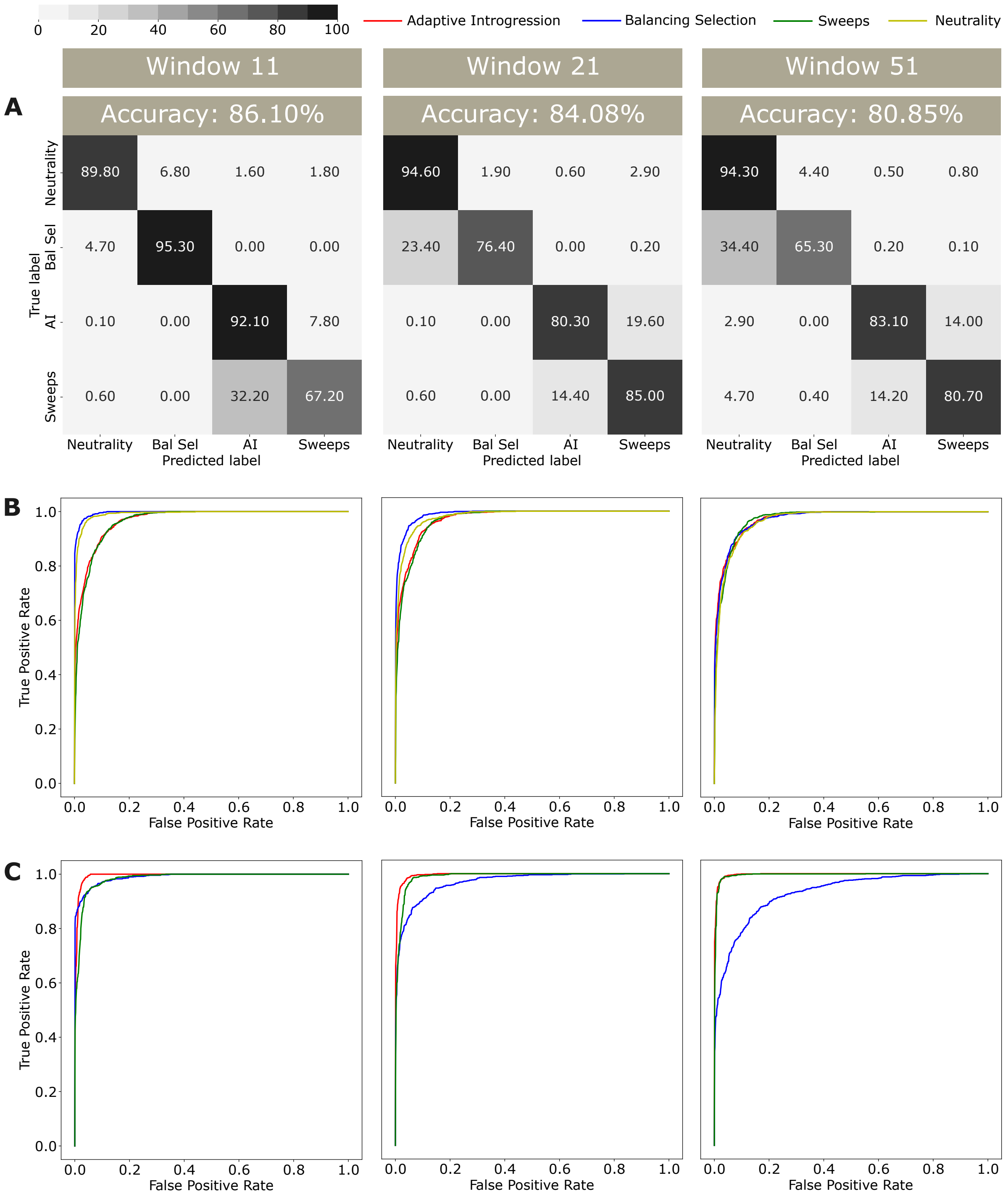


Figure S3: Classification rates and accuracies as depicted by confusion matrices (A) and power to distinguish evolutionary classes as depicted by receiver operating characteristic curves (B, C) for the $M_2$ model with windows of sizes 11 (left column), 21 (middle column), or 51 SNPs (right column). In the confusion matrices, AI denotes adaptive introgression and Bal Sel denotes balancing selection. (B) True positive rate for each class as a function of the false positive rate for all other classes. (C) True positive rate for balancing selection, adaptive introgression, and sweeps as a function of the false positive rate from neutrality.

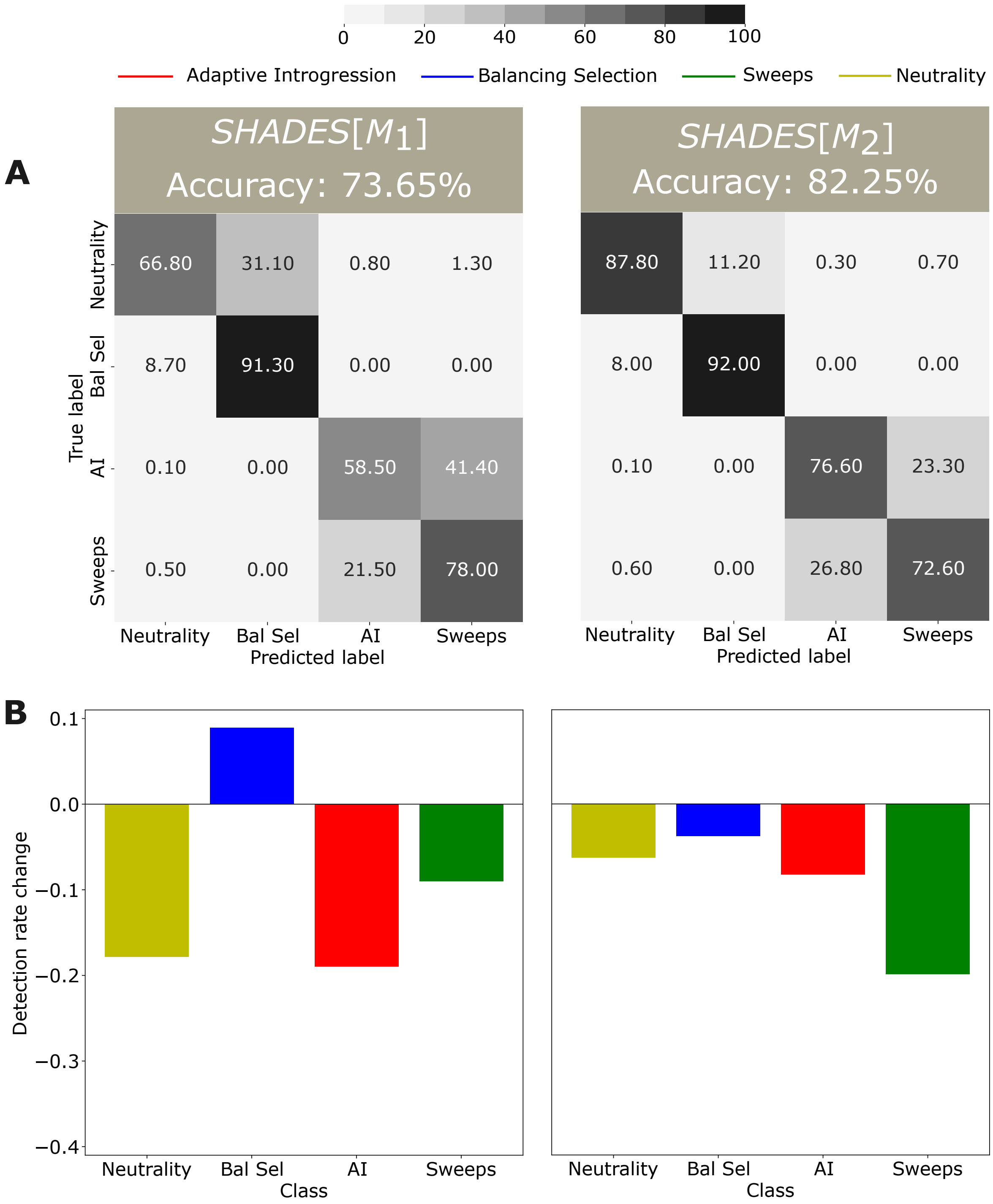


Figure S4: Classification rates and accuracies as depicted by a confusion matrix (A) and evolutionary class detection rate change (B) for the (left) $SHADES[M_1]$ and (right) $SHADES[M_2]$ models under a test setting with low mean recombination rates that are drawn from an exponential distribution with a 10-fold lower mean per-site per-generation recombination rate of $10^{-9}$ compared to the mean recombination rate of the original test dataset. Here, AI denotes adaptive introgression and Bal Sel denotes balancing selection. (B) Detection rate change is quantified as $(X - Y)/Y$, where $X$ is the detection rate for a given evolutionary class with the lower recombination rate in the test set and $Y$ is the detection rate for the same class with the original recombination rate in the test set. Positive values for detection rate change indicate higher detection accuracies under lower recombination rate, whereas negative values result from lower detection accuracies.

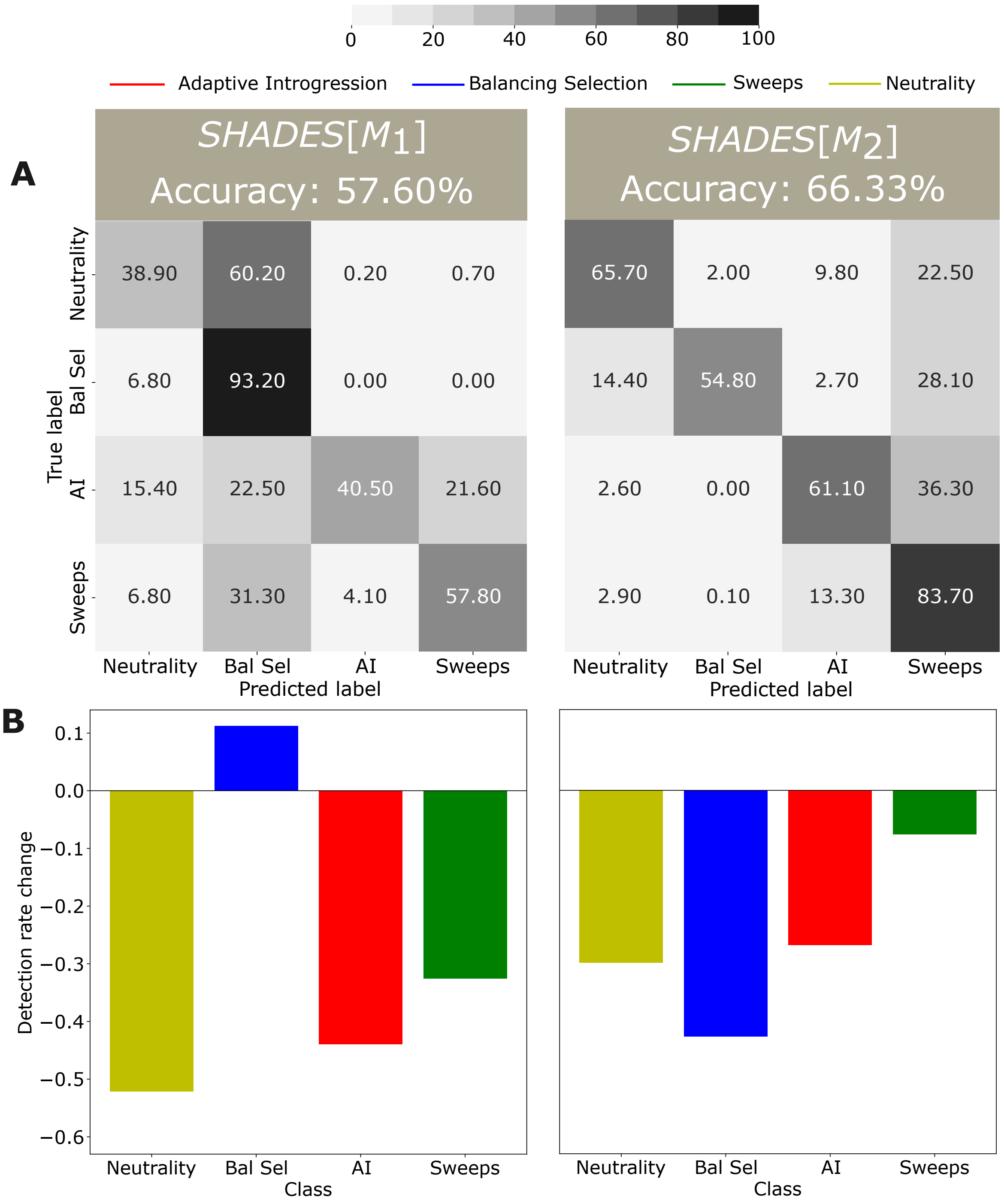


Figure S5: Classification rates and accuracies as depicted by a confusion matrix (A) and evolutionary class detection rate change (B) for the (left) $SHADES[M_1]$ and (right) $SHADES[M_2]$ models under a test setting with high mean recombination rates that are drawn from an exponential distribution with a 10-fold higher mean per-site per-generation recombination rate of $10^{-7}$ compared to the mean recombination rate of the original test dataset. Here, AI denotes adaptive introgression and Bal Sel denotes balancing selection. (B) Detection rate change is quantified as $(X - Y)/Y$, where $X$ is the detection rate for a given evolutionary class with the higher recombination rate in the test set and $Y$ is the detection rate for the same class with the original recombination rate in the test set. Positive values for detection rate change indicate higher detection accuracies under higher recombination rate, whereas negative values result from lower detection accuracies.

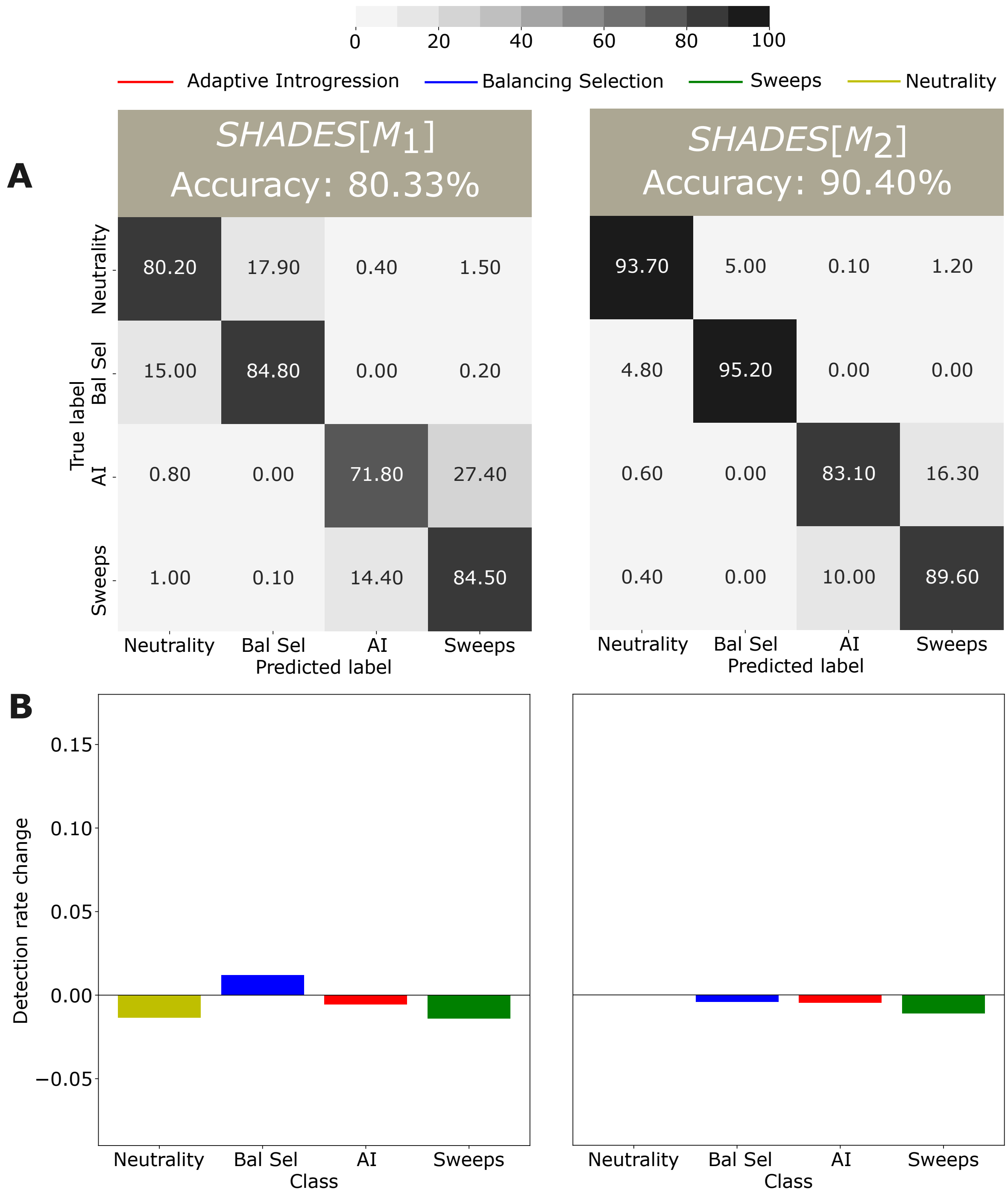


Figure S6: Classification rates and accuracies as depicted by a confusion matrix (A) and evolutionary class detection rate change (B) for the (left) $SHADES[M_1]$ and (right) $SHADES[M_2]$ models under a test with mutation rate variability in which rates are drawn uniformly between $\mu/2$ and $2\mu$ compared to the fixed per-site per-generation mutation rate of $\mu = 1.25 \times 10^{-8}$ of the original test dataset. Here, AI denotes adaptive introgression and Bal Sel denotes balancing selection. (B) Detection rate change is quantified as $(X - Y)/Y$, where $X$ is the detection rate for a given evolutionary class with a variable mutation rate in the test set and $Y$ is the detection rate for the same class with the original fixed mutation rate in the test set. Positive values for detection rate change indicate higher detection accuracies under the variable mutation rate, whereas negative values result from lower detection accuracies.

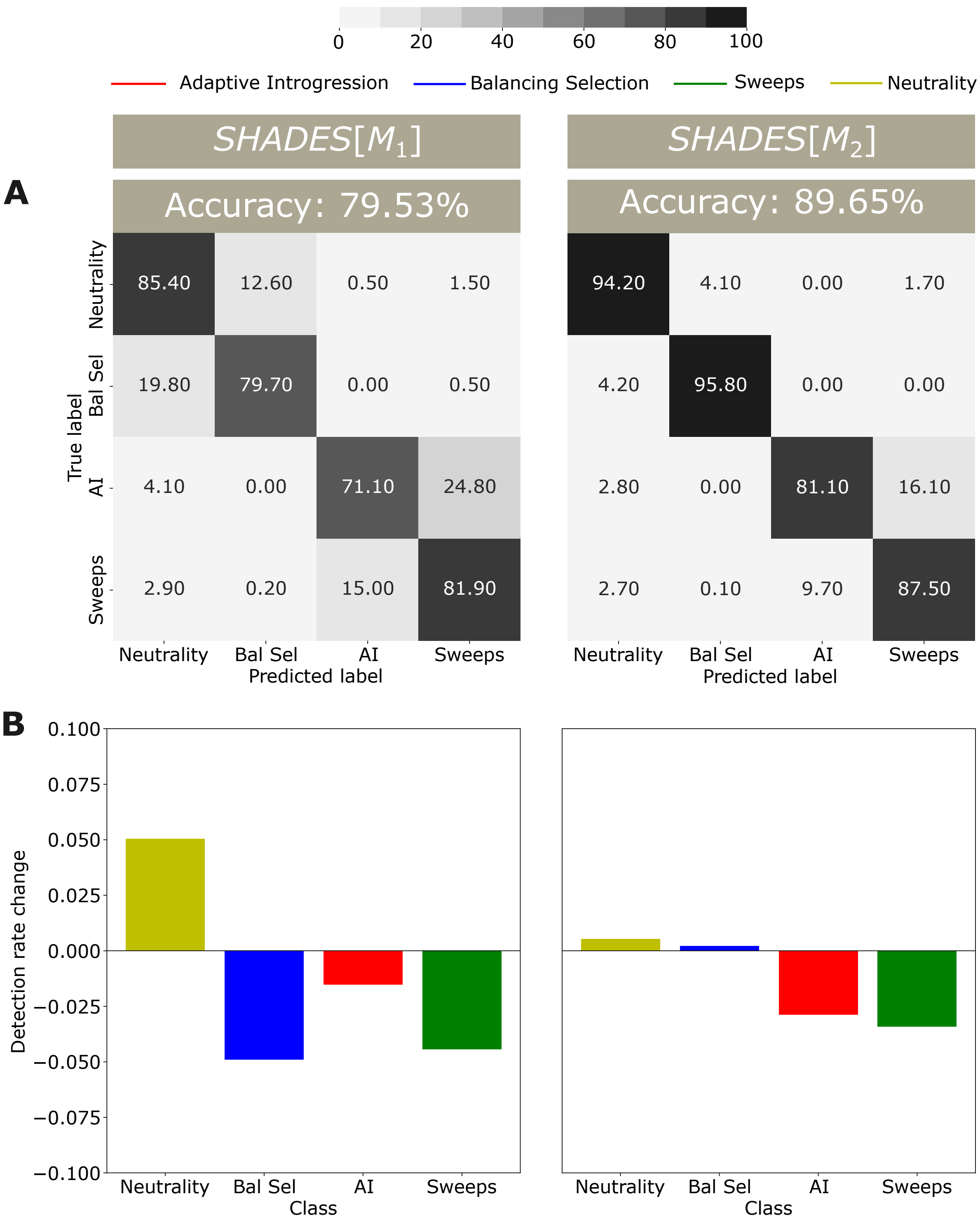


Figure S7: Classification rates and accuracies as depicted by a confusion matrix (A) and evolutionary class detection rate change (B) for the (left) $SHADES[M_1]$ and (right) $SHADES[M_2]$ models under a setting of missing genomic blocks to mimic filtering of low CRG (Centre for Genomic Regulation) score regions. Here, AI denotes adaptive introgression and Bal Sel denotes balancing selection. (B) Detection rate change is quantified as $(X - Y)/Y$, where $X$ is the detection rate for a given evolutionary class with missing genomic blocks in the test set and $Y$ is the detection rate for the same class without missing blocks. Positive values for detection rate change indicate higher detection accuracies under missing genomic blocks, whereas negative values result from lower detection accuracies.

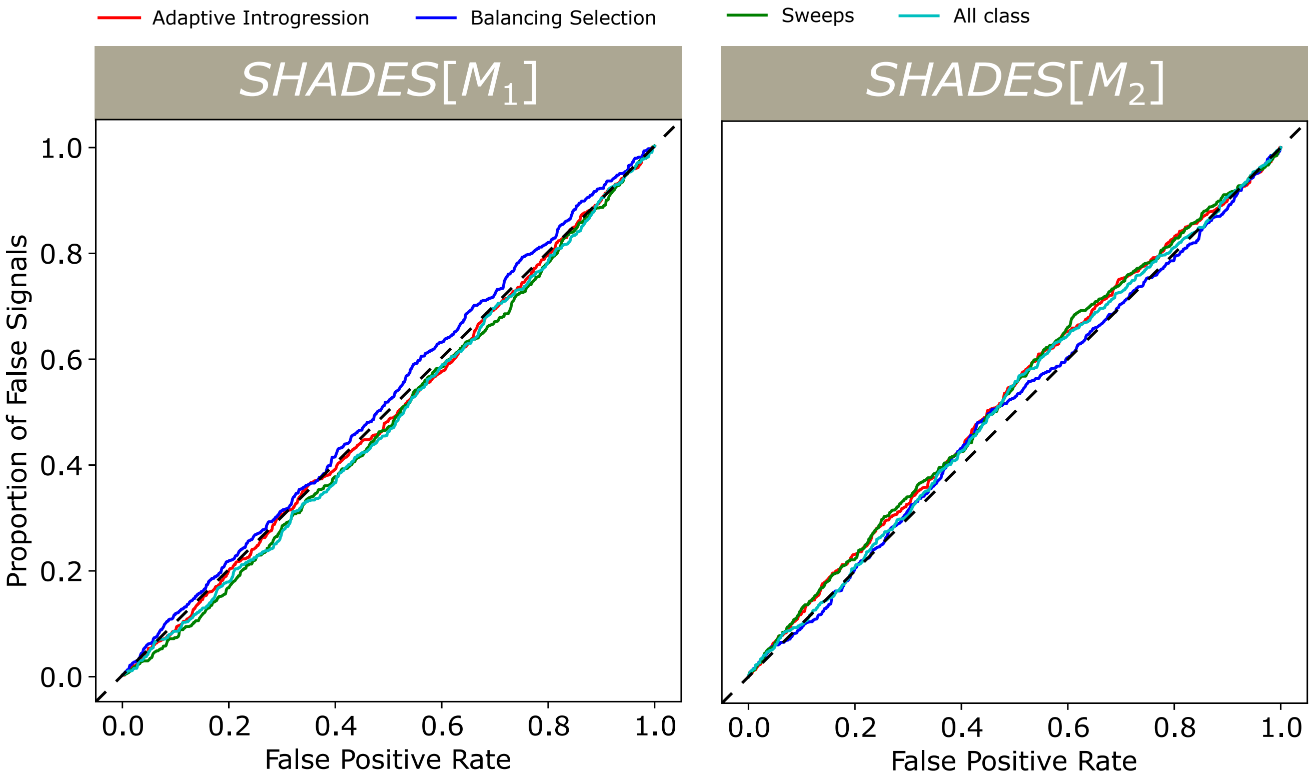


Figure S8: Proportion of false signals under background selection as a function of false positive rate under neutrality for the pretrained $SHADES[M_1]$ and $SHADES[M_2]$ models. The proportion of false signals was defined as the fraction of background selection test replicates with a class probability exceeding the threshold that yielded a given false positive rate in neutral test replicates for that class. The trained *SHADES* models are identical to those from Figure 2. Results closely track the $y = x$ line, indicating consistent classification behavior and robustness to background selection effects when evaluated for each class separately and for all selection classes combined as one class.

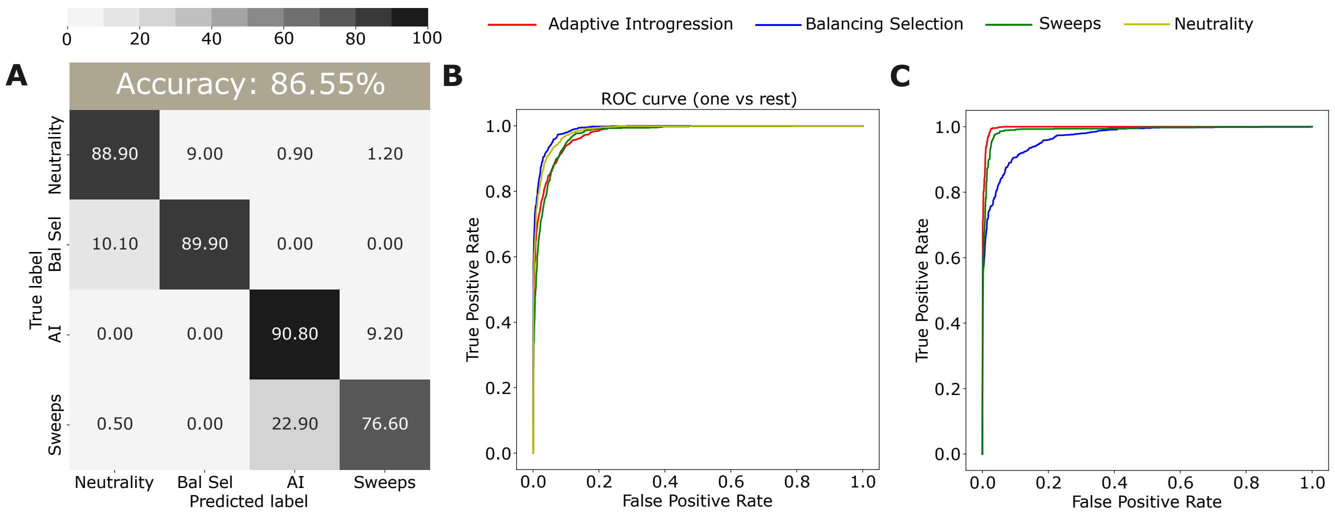


Figure S9: Classification rates and accuracy as depicted by a confusion matrix (A) and power to distinguish evolutionary classes as depicted by receiver operating characteristic curves (B, C) for the $SHADES[M_2]$ model trained and applied to unphased multilocus genotype data. In the confusion matrices, AI denotes adaptive introgression and Bal Sel denotes balancing selection. (B) True positive rate for each class as a function of the false positive rate for all other classes. (C) True positive rate for balancing selection, adaptive introgression, and sweeps as a function of the false positive rate from neutrality. Summary statistic calculations were modified to accommodate the unphased multilocus genotype structure (see *Results* for details).

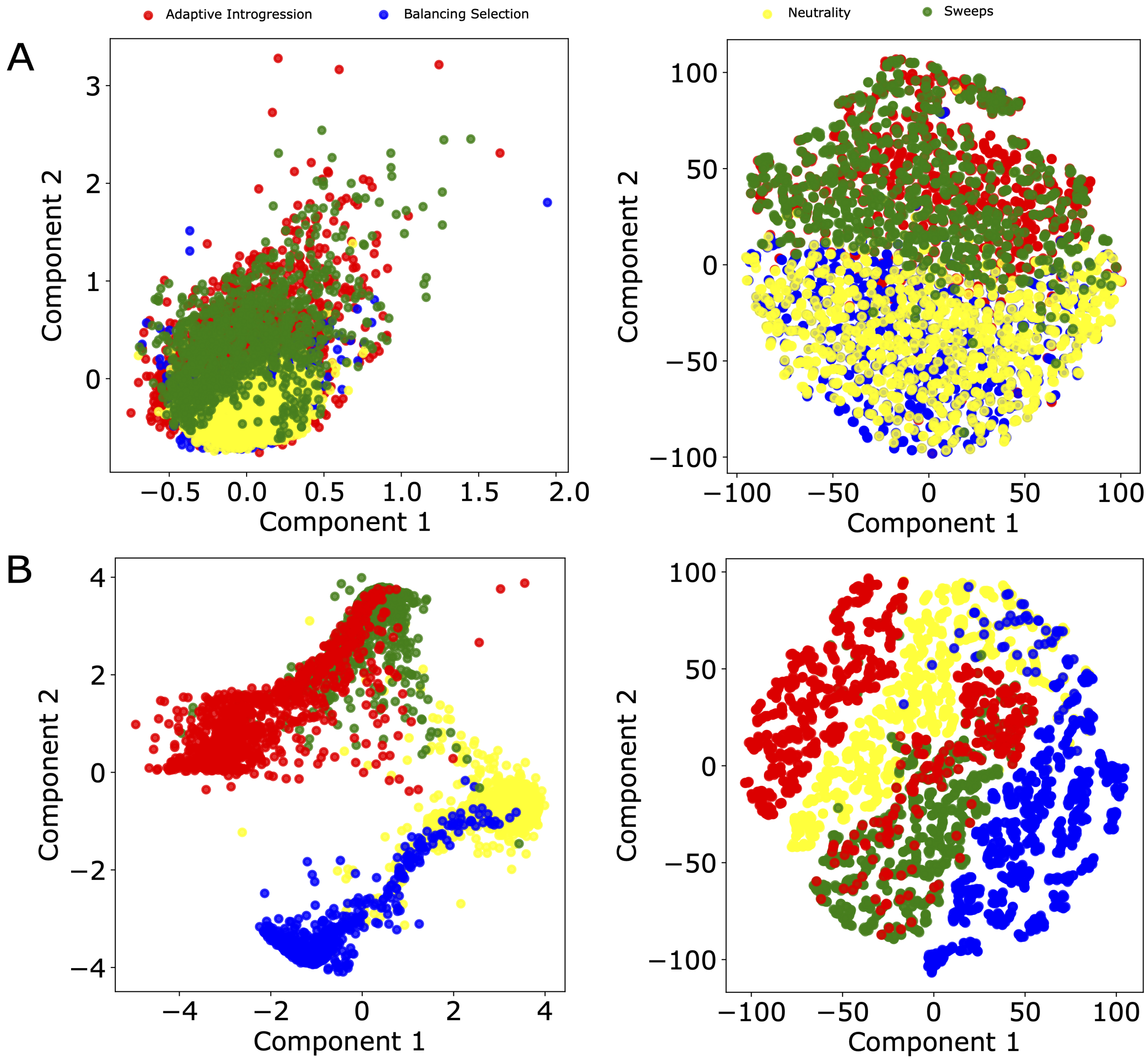


Figure S10: Two-dimensional scatter plot of dimensionality reduction derived from pairwise differences among input image statistics ($\mathbf{I}^{(1)}$, $\mathbf{I}^{(2)}$, and $\mathbf{I}^{(3)}$) of three windows across all test samples. Pairwise differences were normalized, averaged across statistics, and used to construct a sample-by-sample difference matrix, which was then projected by multidimensional scaling (MDS). (A) Scatter plot of the first two MDS components (left) and of a two-dimensional t-SNE embedding computed from the first 50 MDS components (right). (B) Scatter plots of the first two MDS components (left) and of a two-dimensional t-SNE embedding computed from the first 50 MDS components (right) based on the penultimate-layer outputs of the $SHADES[M_2]$ model, representing learned embeddings for the same test samples. Pairwise differences among these embeddings were computed and projected as in panel A. Compared with the raw input statistics, the learned embedding space shows substantially improved separation of the four classes.

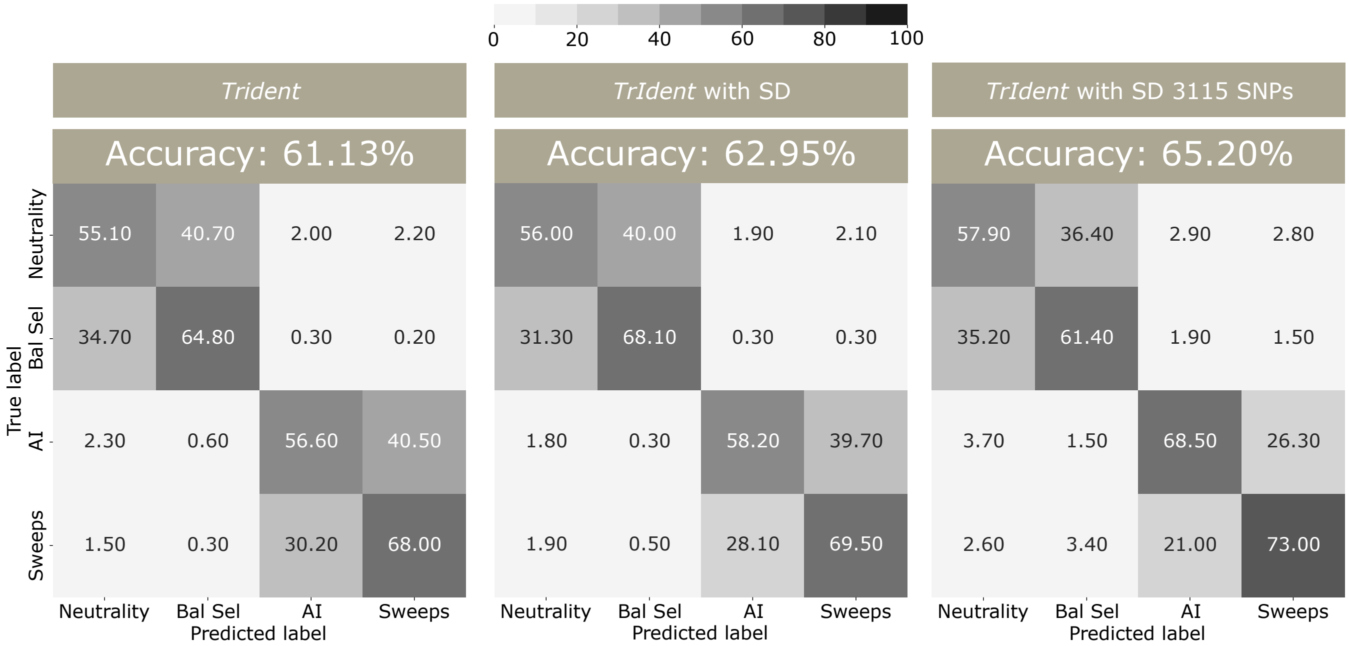


Figure S11: Classification rates and accuracies as depicted by confusion matrices for the default *TrIdent* architecture of Arnab et al. [2025] with singletons and doubletons (SDs) removed and 500 total SNPs (left), a *TrIdent* architecture with SDs retained and still with 500 total SNPs (center), and a *TrIdent* architecture with SDs retained but with 3,115 SNPs. In the confusion matrices, AI denotes adaptive introgression and Bal Sel denotes balancing selection.

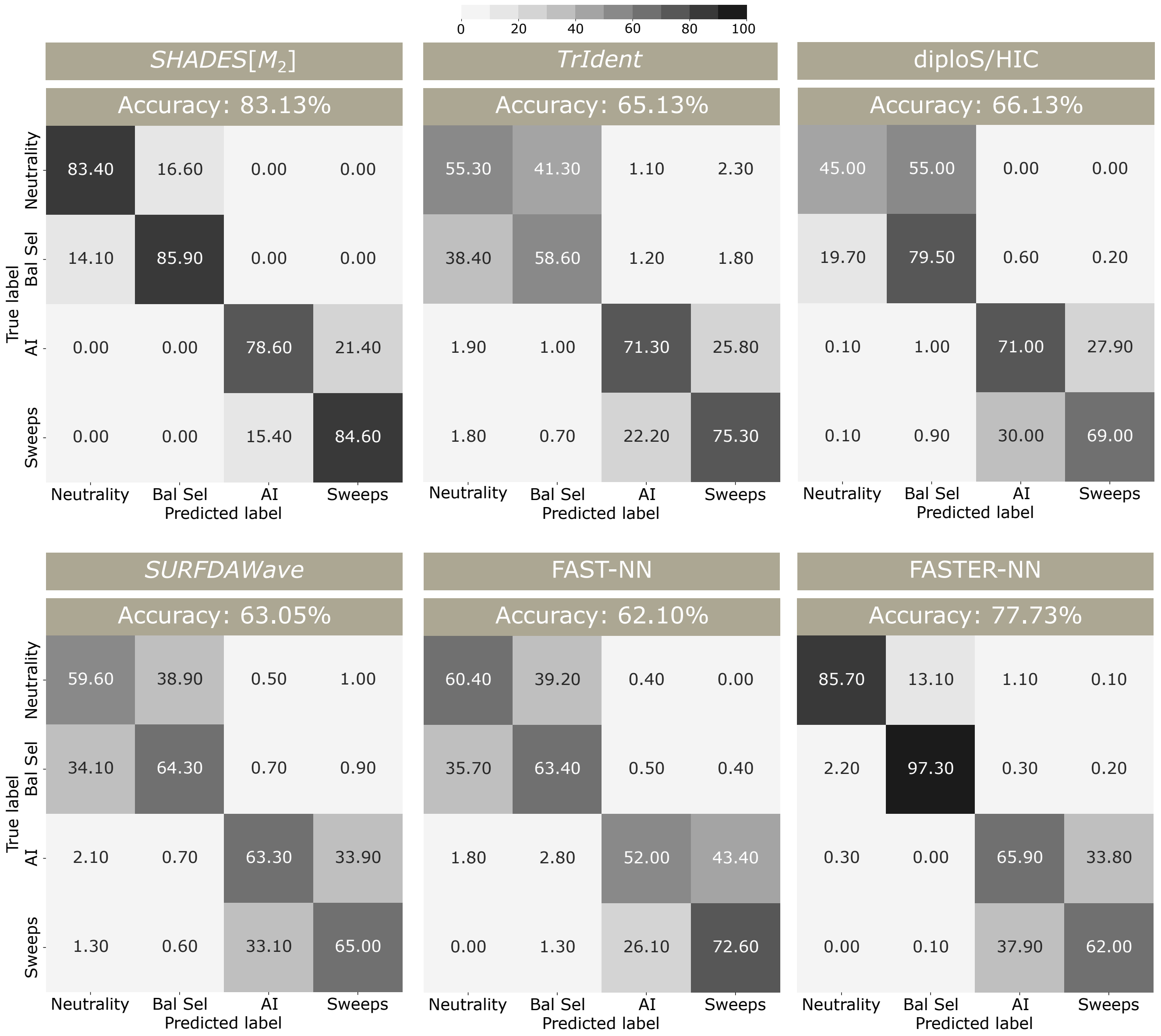


Figure S12: Classification rates and accuracies as depicted by confusion matrices for the *SHADES*[$M_2$], diploS/HIC, *TrIdent*, *SURFDAWave*, FAST-NN, and FASTER-NN models under a complex demographic history with gene flow between an unsampled population with the sampled representing the CEU demographic model. In the confusion matrices, AI denotes adaptive introgression and Bal Sel denotes balancing selection. The *TrIdent* model was trained and tested using the central 3,115 SNPs from each test genome replicate while retaining singletons and doubletons. The diploS/HIC model was trained and tested with 12 summary statistics computed on 21 contiguous windows of length approximately 95 kilobases across the two Mb simulated region. The *SURFDAWave* model was trained and tested with nine summary statistics computed in 256 overlapping windows of length 10 SNPs with a stride of five SNPs. FAST-NN was trained and tested on derived allele frequencies alone, whereas FASTER-NN additionally incorporated genomic position information.

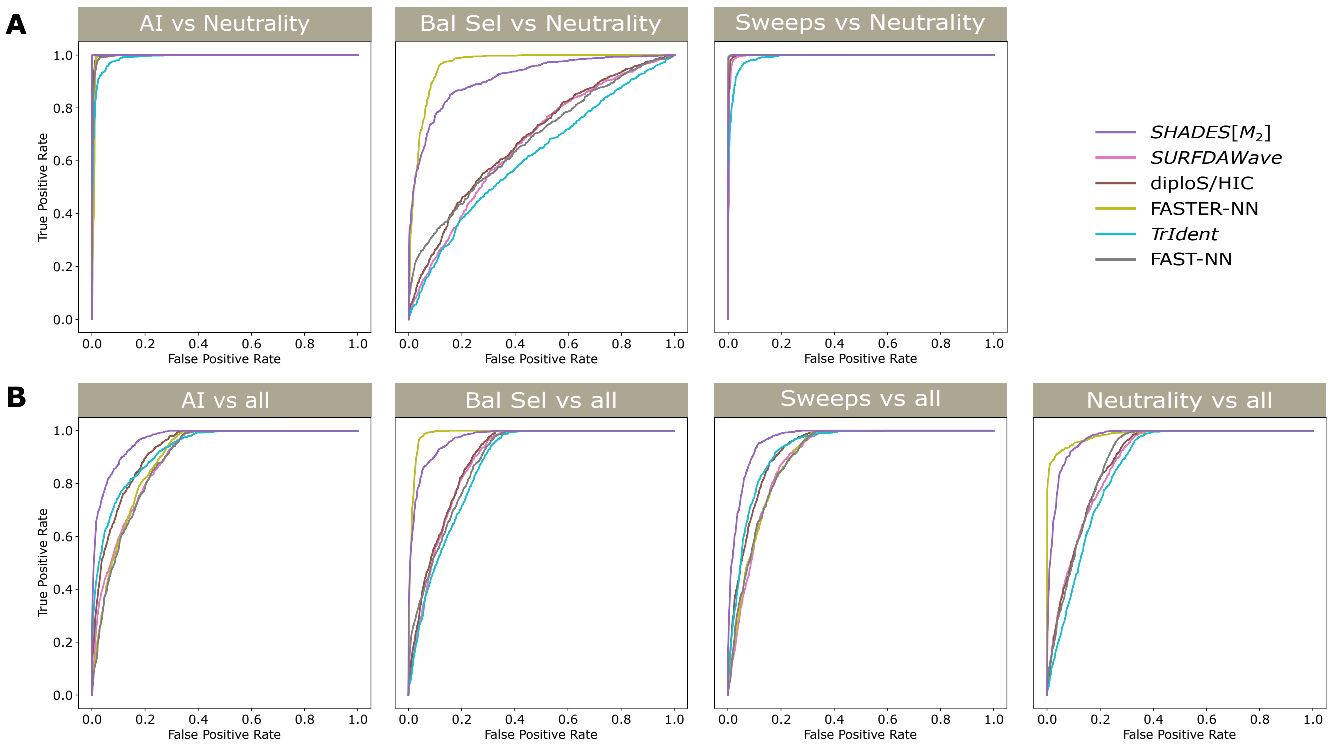


Figure S13: Receiver operating characteristic curves comparing $SHADES[M_2]$, diploS/HIC, *TrIdent*, *SURFDAWave*, FAST-NN, and FASTER-NN models across two evaluation settings under a complex demographic history with gene flow between an unsampled population with the sampled population representing the CEU demographic model. Here, AI denotes adaptive introgression and Bal Sel denotes balancing selection. (A) True positive rate for each selective class as a function of the false positive rate from neutrality. (B) True positive rate for each class as a function of the false positive rate computed from all remaining classes.

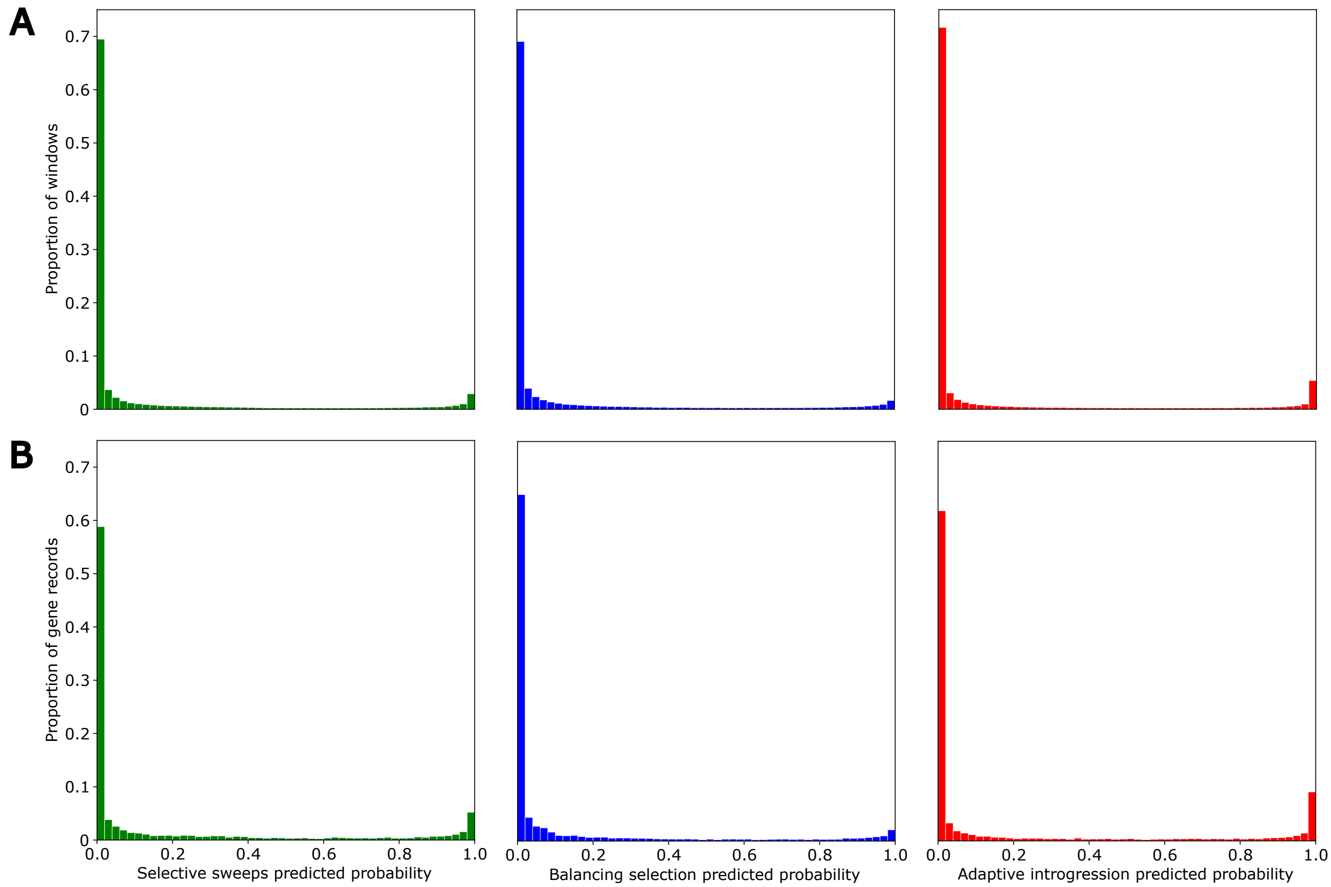


Figure S14: Genome-wide distributions of $SHADES[M_2]$ predicted probabilities for selective sweep, balancing selection, and adaptive introgression in the empirical CEU scan. (A) Distributions of predicted probabilities across all scanned genomic windows. Each distribution shows the proportion of windows within each predicted probability interval. (B) Distribution of gene-level predicted probabilities. Scores were computed as the maximum predicted probability for a given class intersecting the start and stop of a protein-coding gene, and the distribution is across all genes for which a score was assigned. All distributions sum to one, and most windows (A) and genes (B) received confidence scores at or near zero, whereas only a small subset received scores greater than 0.9.

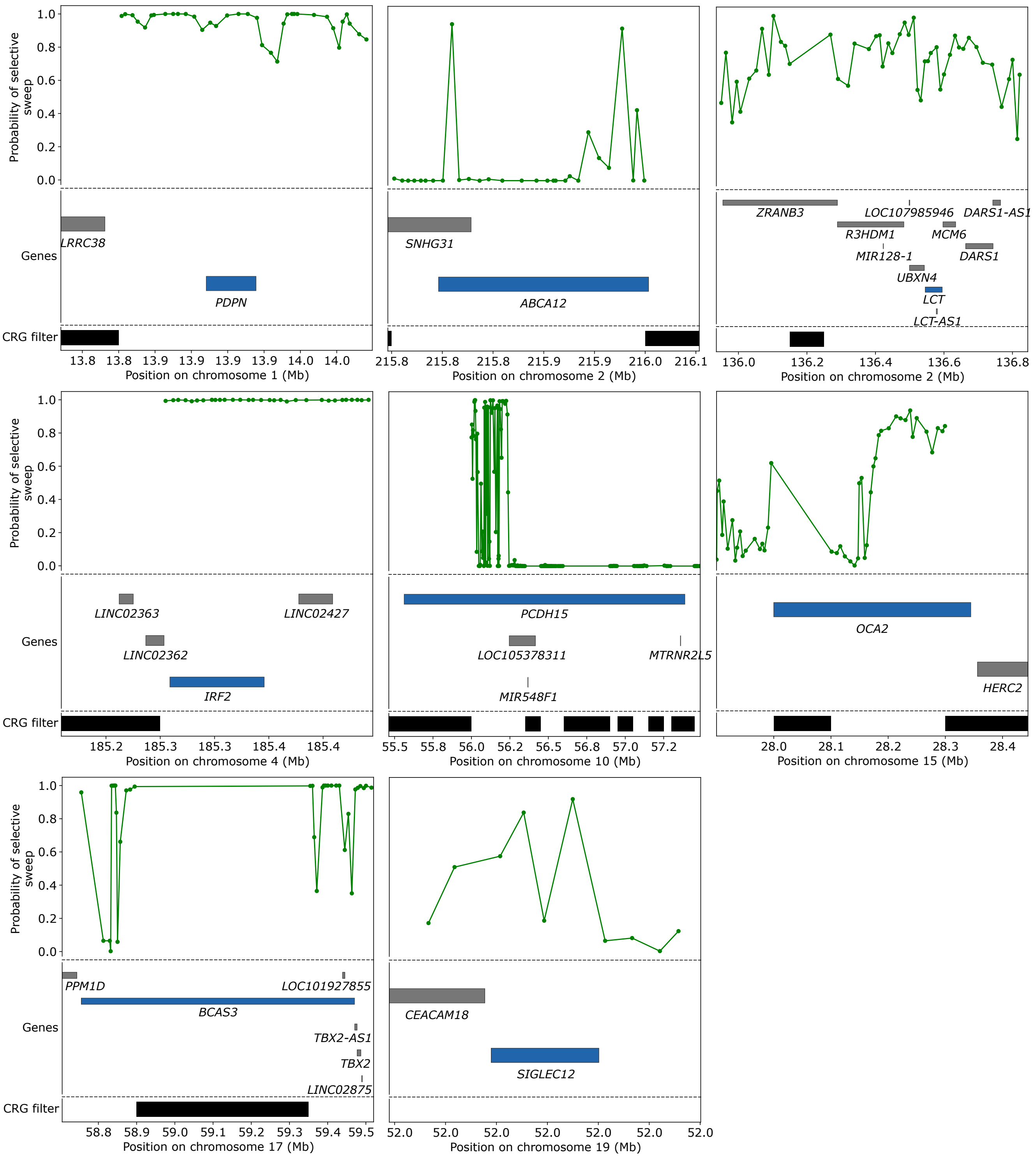


Figure S15: Genomic spatial pattern of $SHADES[M_2]$ predicted sweep probabilities across a set of highlighted candidate genes within the empirical CEU scan. Protein coding genes and their genomic coordinates overlapping each highlighted region are shown below the predicted probabilities as horizontal gray bars, with the focal gene at which the signal is centered shown as a horizontal blue bar. CRG filtered intervals are displayed at the bottom as horizontal black bars to indicate genomic segments containing filtered or missing data.

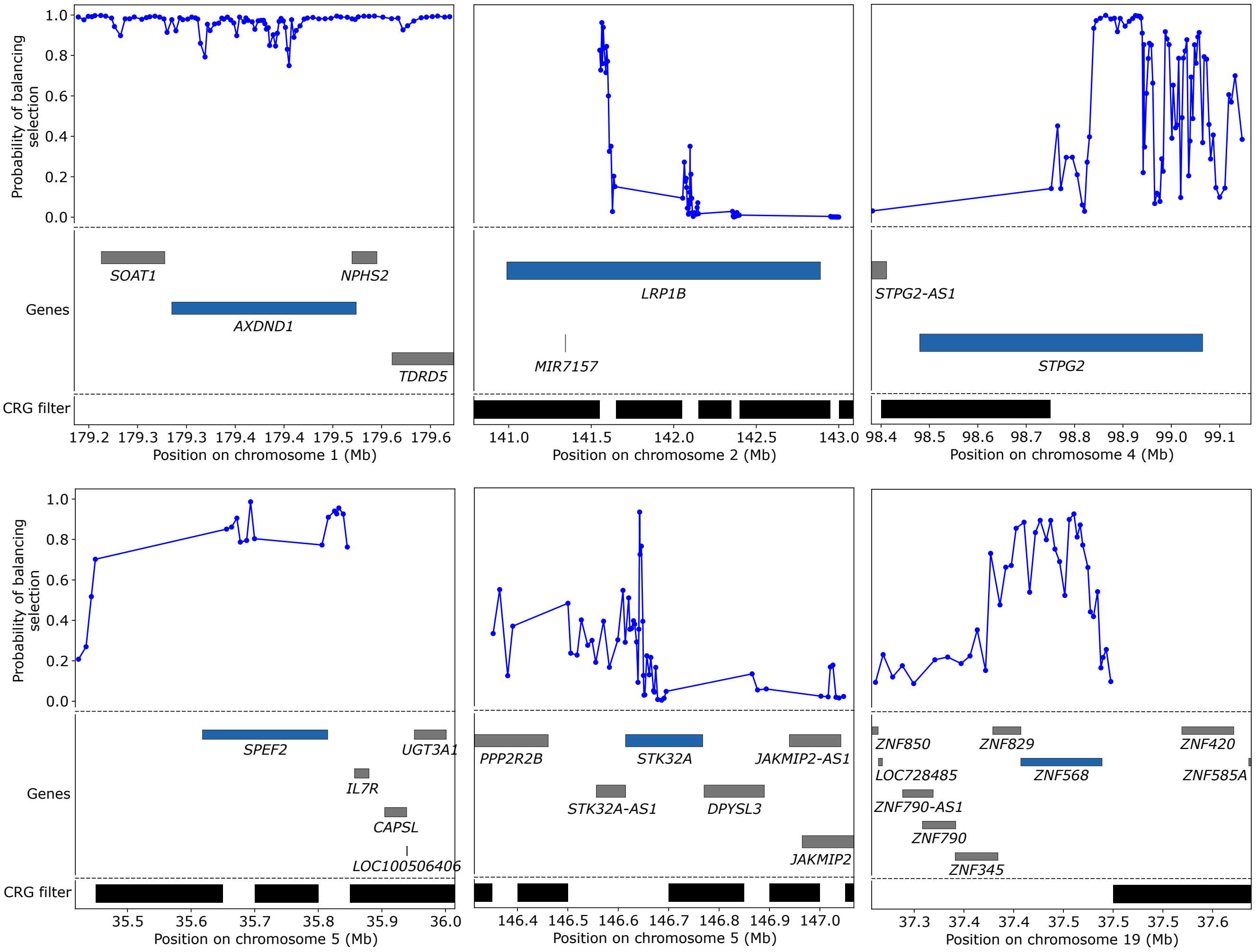


Figure S16: Genomic spatial pattern of $SHADES[M_2]$ predicted balancing selection probabilities across a set of highlighted candidate genes within the empirical CEU scan. Protein coding genes and their genomic coordinates overlapping each highlighted region are shown below the predicted probabilities as horizontal gray bars, with the focal gene at which the signal is centered shown as a horizontal blue bar. CRG filtered intervals are displayed at the bottom as horizontal black bars to indicate genomic segments containing filtered or missing data.

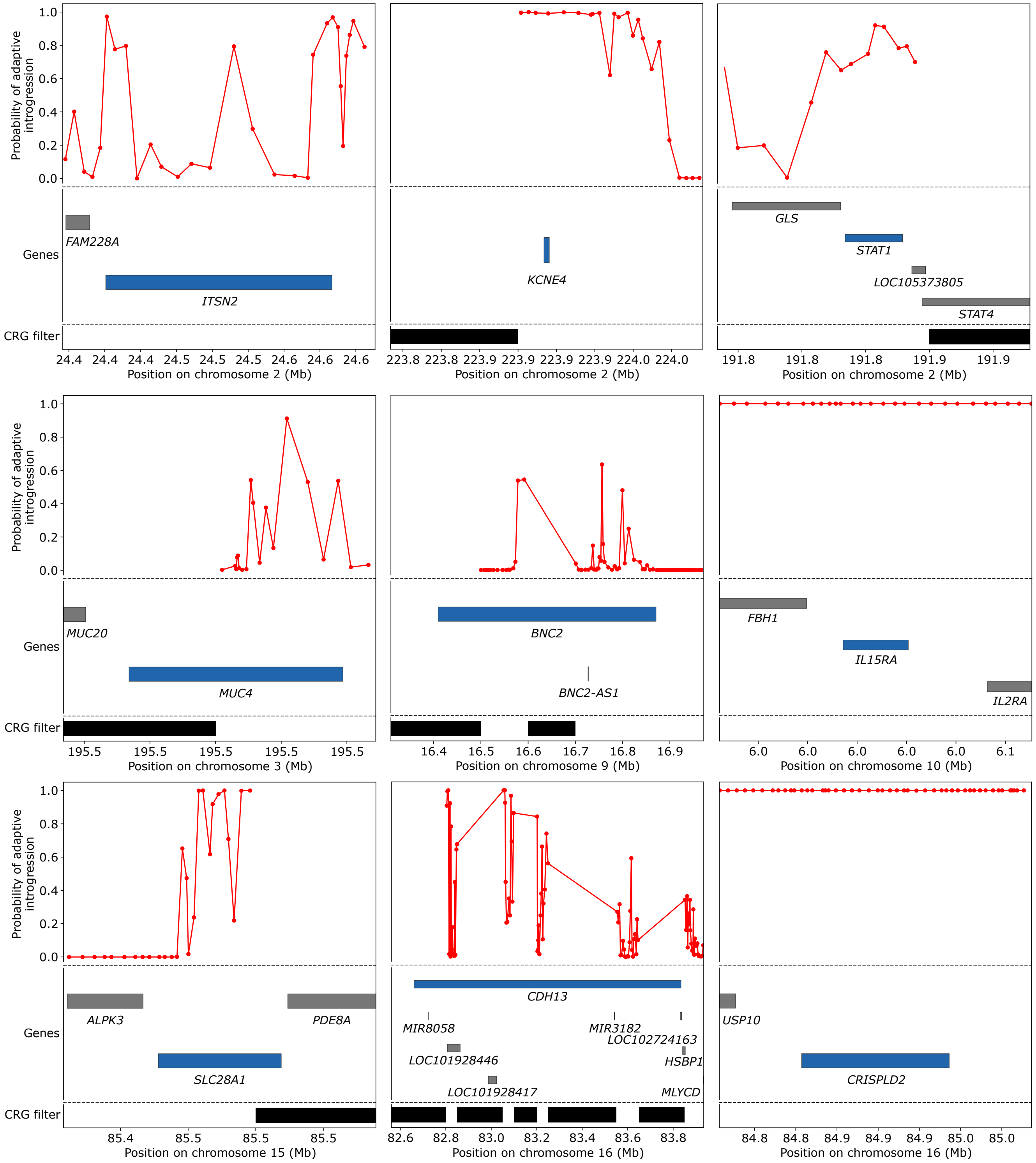


Figure S17: Genomic spatial pattern of $SHADES[M_2]$ predicted adaptive introgression probabilities across a set of highlighted candidate genes within the empirical CEU scan. Protein coding genes and their genomic coordinates overlapping each highlighted region are shown below the predicted probabilities as horizontal gray bars, with the focal gene at which the signal is centered shown as a horizontal blue bar. CRG filtered intervals are displayed at the bottom as horizontal black bars to indicate genomic segments containing filtered or missing data.

Table S1: $SHADES[M_2]$ predictions across the 22 autosomes of a European (CEU) population from the 1000 Genomes Project dataset. For each scanned gene, the highest confidence score is shown for selective sweeps, balancing selection, adaptive introgression, and neutrality. See file Table-CEU-1000-genome.csv

Table S2: Gene Ontology (GO) cellular components for genes ranked based on *SHADES*$[M_2]$ prediction output for adaptive introgression.

| GO Term | Description | FDR $q$-value | Enrichment | Genes |
|---|---|---|---|---|
| GO:0016020 | membrane | 0.0152 | 1.29 | *ERGIC1, SLC43A2, ITPR2, BIK, JAK3, ATP6V0E1, GALNT13, HK1, SAG, CD82, VPS37A, PAK2, PALM, HLA-DRB1, HLA-DRB5, KCNG1, SPAG9, KCNJ1, TBXA2R, KCNJ5, GPRC5A, PIP5K1B, KCNJ16, CCDC124, TLN2, KCNMA1, BZW2, TSPO, PN-PLA6, KCNQ2, PTBP1, NUBPL, MSR1, RUFY1, PIGZ, PALLD, MFSD12, ADAM10, DDX47, ELAVL1, HNRNPH1, SCUBE1, SULF2, ARHGAP33, MTMR7, PDE9A, LDLRAD4, B3GNT3, PTPN1, CLEC4M, PTPN3, RP-TOR, CHMP4A, MDGA2, PT-PRD, CACNB2, PTPRS, OSBP2, CHST8, SLC28A1, DGKI, GDI2, TTYH3, DLC1, CALU, MX1, MX2, DGKD, CANX, HRH1, TGM1, PARVB, FCHO1, ST3GAL4, VCAN, EPB41L4B, MICU3, CHMP6, NGEF, UGT1A10, MAGI1, UGT1A8, UGT1A7, FZR1, UGT1A6, DLGAP1, UGT1A5, ARHGAP10, EXT2, UGT1A9, SAMM50, TIMM44, UGT1A4, SLC5A5, UGT1A3, GNA12, GNA15, SULF1, LOXL2, ICA1, LPP, ATP2C2, P2RY12, ARL6IP6, SLC19A1, PLCG2, APBB2, SCFD1, NDUFS5, APLP1, TM9SF1, GPRC5D, IQCE, NRXN3,* |

| | | | | |
|---|---|---|---|---|
| | | | | *DGKB, CD209, TSPAN13, TPCN2, TM9SF2, SORBS1, CACNG4, NCLN, FHIT, IRAK3, PRPF8, NMT2, CHRNA9, RHOH, CNTN4, TJP3, NALCN, MATK, FMN1, PACSIN2, FAM174B, TMBIM4, PRODH2, MBP, ZDHHC7, TNFRSF10D, TMEM167A, TNFRSF10C, TNFRSF10B, PIP5K1C, NPHS1, GIGYF2, NTN4, IL2RA, COTL1, IL5RA, DPM1, RHBDL2, APOLD1, CARD11, GRIP1, SHANK1, MREG, LZTS2, AKAP1, GAL3ST1, IL15RA, GALNT18, SPTLC2, IL16, MEGF11, PIGX, PDZD7, MGST3, PES1, APBA3, INPP5D, PRKCA, MRPL43, EBI3, PRKCE, MACF1, GSG1L, GABBR2, ARID3A, THSD7A, CLCN2, PARD6B, MGAT5B, SLC39A14, TXNDC11, CLRN1, KIRREL3, STXBP5, DYSF, CDHR3, AKAP12, SLCO3A1, SNN, LRRC16B, MCTP2, NRROS, SLC17A9, NCOR2, SYPL1, TSPAN15, TACR2, SLC37A1* |
| GO:0071944 | cell periphery | 0.0112 | 5.42 | *MAGI1, ITGB1BP1, NME2, ATP1B2, SLC1A1, THSD1, MYO1B, PRKCE, ACTG2, MBP, NOS1* |

Table S3: Gene Ontology (GO) molecular functions for genes ranked based on $SHADES[M_2]$ prediction output for adaptive introgression.

| GO Term | Description | FDR $q$-value | Enrichment | Genes |
|---|---|---|---|---|
| GO:0015020 | glucuronosyltransferase activity | $5.17 \times 10^{-8}$ | 49.15 | *UGT1A9, UGT1A10, UGT1A8, UGT1A7, UGT1A6, UGT1A5, UGT1A4, UGT1A3* |
| GO:0008194 | UDP-glycosyltransferase activity | $4.40 \times 10^{-5}$ | 14.4 | *UGT1A9, GALNT13, UGT1A10, UGT1A8, UGT1A7, UGT1A6, UGT1A5, GALNT18, UGT1A4, UGT1A3* |
| GO:0016758 | transferase activity, transferring hexosyl groups | $3.75 \times 10^{-4}$ | 9.57 | *UGT1A9, GALNT13, UGT1A10, UGT1A8, UGT1A7, UGT1A6, UGT1A5, GALNT18, UGT1A4, DPM1, UGT1A3* |
| GO:0016757 | transferase activity, transferring glycosyl groups | $5.61 \times 10^{-4}$ | 7.26 | *UGT1A9, GALNT13, UGT1A10, UGT1A8, UGT1A7, UGT1A6, UGT1A5, GALNT18, NAMPT, UGT1A4, DPM1, UGT1A3* |
| GO:0001972 | retinoic acid binding | $9.02 \times 10^{-3}$ | 40.21 | *UGT1A9, UGT1A8, UGT1A7, UGT1A3* |
| GO:0005501 | retinoid binding | $4.32 \times 10^{-2}$ | 27.65 | *UGT1A9, UGT1A8, UGT1A7, UGT1A3* |
| GO:0019840 | isoprenoid binding | $4.67 \times 10^{-2}$ | 26.02 | *UGT1A9, UGT1A8, UGT1A7, UGT1A3* |
| GO:0042605 | peptide antigen binding | $4.96 \times 10^{-2}$ | 14.86 | *CD209, MAML1, HLA-DRB5, CLEC4M* |

Table S4: Gene Ontology (GO) biological processes genes ranked based on $SHADES[M_2]$ prediction output for adaptive introgression.

| GO Term | Description | FDR $q$-value | Enrichment | Genes |
|---|---|---|---|---|
| GO:0052696 | flavonoid glucuronidation | $3.65 \times 10^{-11}$ | 98.3 | *UGT1A9, UGT1A10, UGT1A8, UGT1A7, UGT1A6, UGT1A5, UGT1A4, UGT1A3* |
| GO:0052697 | xenobiotic glucuronidation | $1.83 \times 10^{-11}$ | 98.3 | *UGT1A9, UGT1A10, UGT1A8, UGT1A7, UGT1A6, UGT1A5, UGT1A4, UGT1A3* |
| GO:0009812 | flavonoid metabolic process | $1.53 \times 10^{-10}$ | 80.43 | *UGT1A9, UGT1A10, UGT1A8, UGT1A7, UGT1A6, UGT1A5, UGT1A4, UGT1A3* |
| GO:0052695 | cellular glucuronidation | $1.11 \times 10^{-9}$ | 68.06 | *UGT1A9, UGT1A10, UGT1A8, UGT1A7, UGT1A6, UGT1A5, UGT1A4, UGT1A3* |
| GO:0006063 | uronic acid metabolic process | $5.00 \times 10^{-9}$ | 58.98 | *UGT1A9, UGT1A10, UGT1A8, UGT1A7, UGT1A6, UGT1A5, UGT1A4, UGT1A3* |
| GO:0019585 | glucuronate metabolic process | $4.17 \times 10^{-9}$ | 58.98 | *UGT1A9, UGT1A10, UGT1A8, UGT1A7, UGT1A6, UGT1A5, UGT1A4, UGT1A3* |
| GO:0051552 | flavone metabolic process | $5.91 \times 10^{-4}$ | 76.13 | *UGT1A9, UGT1A10, UGT1A8, UGT1A7* |
| GO:0006805 | xenobiotic metabolic process | $1.44 \times 10^{-3}$ | 11.65 | *UGT1A9, UGT1A10, UGT1A8, UGT1A7, UGT1A6, UGT1A5, UGT1A4, MGST3, UGT1A3* |
| GO:0005996 | monosaccharide metabolic process | $3.00 \times 10^{-3}$ | 9.14 | *UGT1A9, HK1, UGT1A10, UGT1A8, UGT1A7, UGT1A6, UGT1A5, PFKFB3, UGT1A4, UGT1A3* |
| GO:0032787 | monocarboxylic acid metabolic process | $4.90 \times 10^{-2}$ | 4.61 | *UGT1A10, UGT1A8, LIPC, UGT1A7, UGT1A6, UGT1A5, PRODH2, UGT1A9, HK1, VCAN, MCAT, UGT1A4, UGT1A3* |

Table S5: Gene Ontology (GO) biological processes significantly enriched among genes with $SHADES[M_2]$ adaptive introgression confidence scores greater than 0.9 relative to a background set of all retained genes.

| GO Term | Description | FDR $q$-value | Enrichment | Genes |
|---|---|---|---|---|
| GO:0052696 | flavonoid glucuronidation | $1.31 \times 10^{-4}$ | 7.76 | *UGT1A9, UGT1A10, UGT1A8, UGT1A7, UGT1A6, UGT1A5, UGT1A4, UGT1A1, UGT1A3* |
| GO:0052697 | xenobiotic glucuronidation | $6.53 \times 10^{-5}$ | 7.76 | *UGT1A9, UGT1A10, UGT1A8, UGT1A7, UGT1A6, UGT1A5, UGT1A4, UGT1A1, UGT1A3* |
| GO:0009812 | flavonoid metabolic process | $5.42 \times 10^{-5}$ | 7.06 | *UGT1A9, UGT1A10, UGT1A8, UGT1A7, PPARGC1A, UGT1A6, UGT1A5, UGT1A4, UGT1A1, UGT1A3* |
| GO:0052695 | cellular glucuronidation | $1.43 \times 10^{-2}$ | 5.38 | *UGT1A9, UGT1A10, UGT1A8, UGT1A7, UGT1A6, UGT1A5, UGT1A4, UGT1A1, UGT1A3* |
| GO:0051552 | flavone metabolic process | $1.24 \times 10^{-2}$ | 7.76 | *UGT1A9, UGT1A10, UGT1A8, UGT1A7, PPARGC1A, UGT1A1* |
| GO:0019585 | glucuronate metabolic process | $4.49 \times 10^{-2}$ | 4.66 | *UGT1A9, UGT1A10, UGT1A8, UGT1A7, UGT1A6, UGT1A5, UGT1A4, UGT1A1, UGT1A3* |

Table S6: Gene Ontology (GO) cellular components for genes ranked based on $SHADES[M_2]$ prediction output for balancing selection.

| GO Term | Description | FDR $q$-value | Enrichment | Genes |
|---|---|---|---|---|
| GO:0005882 | intermediate filament | $9.75 \times 10^{-5}$ | 7.57 | *KRTAP4-8, KRT39, KRTAP2-1, KRTAP4-12, KRT38, KRTAP1-4, KRT23, KRT31, KRTAP4-11, KRT33B, KRT40, KRTAP4-9, KRT28, KRT20* |

Table S7: Gene Ontology (GO) molecular functions for genes ranked based on $SHADES[M_2]$ prediction output for balancing selection.

| GO Term | Description | FDR $q$-value | Enrichment | Genes |
|---|---|---|---|---|
| GO:0005149 | interleukin-1 receptor binding | $9.49 \times 10^{-4}$ | 8.31 | *IL1A, IL1B, IRAK4, IL1RN, IL36B, IL37, TRIP6, IL1F10, IL36RN* |

Table S8: Gene Ontology (GO) biological processes for genes ranked based on $SHADES[M_2]$ prediction output for balancing selection.

| GO Term | Description | FDR $q$-value | Enrichment | Genes |
|---|---|---|---|---|
| GO:0031424 | keratinization | $3.90 \times 10^{-6}$ | 10.93 | *KRTAP4-8, KRT39, KRTAP2-1, KRT38, KRTAP1-4, KRT23, PPHLN1, KRT31, KRTAP4-11, KRT33B, KRT40, KRTAP4-9, KRT28, KRT20* |

Table S9: Gene Ontology (GO) cellular components significantly enriched among genes with $SHADES[M_2]$ balancing selection confidence scores greater than 0.9 relative to a background set of all retained genes.

| GO Term | Description | FDR $q$-value | Enrichment | Genes |
|---|---|---|---|---|
| GO:0005882 | intermediate filament | $4.24 \times 10^{-4}$ | 4.12 | *KRTAP4-8, KRT39, KRTAP2-1, KRTAP4-12, KRT38, KRT37, LMNB1, KRTAP1-4, KRT23, KRT31, KRTAP4-11, KRT33B, KRT40, KRTAP4-9, PKP2, KRTAP17-1, KRT28, KRT20* |

Table S10: Gene Ontology (GO) biological processes significantly enriched among genes with $SHADES[M_2]$ balancing selection confidence scores greater than 0.9 relative to a background set of all retained genes.

| GO Term | Description | FDR $q$-value | Enrichment | Genes |
|---|---|---|---|---|
| GO:0031424 | keratinization | $2.26 \times 10^{-5}$ | 5.87 | *KRTAP4-8, KRT39, KRTAP2-1, KRT38, KRT37, KRTAP1-4, KRT23, PPHLN1, KRT31, KRTAP4-11, KRT33B, KRT40, PKP2, KRTAP4-9, KRTAP17-1, KRT28, KRT20* |
